# eIF4A inhibition disrupts resistance-associated translational and metabolic programs in BRAF-mutant melanoma

**DOI:** 10.64898/2026.09.03.749206

**Authors:** Alejandro Schcolnik-Cabrera, Meriem Takdenti, Anavasadat Sadr Hashemi Nejad, Zaynab Nouhi, Sarah St-Amand, Caroline Capdevielle, Donia Issa, Melody Riaud, April A. N. Rose, Mireille Khacho, Frédérick A. Mallette, Martin Roffe, Tommy Alain, Ivan Topisirovic, Laura Hulea

## Abstract

Acquired resistance to mitogen-activated protein kinase (MAPK) pathway inhibitors remains a major barrier to durable control of BRAF-mutant melanoma. Although resistance mechanisms are heterogeneous, they converge on adaptive programs that support survival, phenotypic plasticity, and metabolic fitness. We asked whether eukaryotic translation initiation factor 4A (eIF4A)-dependent mRNA translation represents a shared vulnerability of kinase inhibitor-resistant melanoma. Using matched BRAF^V600E^ A375 and BRAF inhibitor-resistant A375R cells together with additional melanoma models, we integrated pharmacological and functional assays with polysome-associated RNA sequencing, quantitative proteomics, bioenergetic profiling, metabolomics, [U-¹³C₅]glutamine tracing, and xenograft studies. Melanoma cells remained sensitive to multiple eIF4A inhibitors regardless of their responsiveness to BRAF inhibition. The eIF4A inhibitor CR-1-31-B rapidly reduced nascent protein synthesis when used alone in A375 cells and when added to the BRAF inhibitor PLX4032 in A375R cells; it also reduced BCL-2, CDK4, and cyclin D3 abundance, suppressed clonogenic growth, and induced apoptosis. Integrated analysis showed that acquired resistance involved broad RNA-abundance remodeling with superimposed changes in translational efficiency and buffering, affecting survival, extracellular-matrix and plasticity programs, and mitochondrial and metabolic functions. In resistant cells, CR-1-31-B induced early transcript-selective translational changes, accompanied at later time points by RNA-abundance and proteome remodeling. Publicly annotated 5′ untranslated regions (5′UTRs) of CR-1-31-B-sensitive transcripts were enriched for purine-rich sequence architecture and local structural complexity. eIF4A inhibition preferentially attenuated the expression of proteins acquired during resistance and imposed a lower-output metabolic state in sensitive and resistant cells, reducing tricarboxylic-acid-cycle and pentose-phosphate-pathway metabolite pools and restricting intracellular glutamine-carbon transfer downstream of uptake. In A375 xenografts, CR-1-31-B delayed tumor growth, while its combination with PLX4720 produced deeper and more sustained tumor control and prolonged tumor endpoint-free survival compared with PLX4720 alone. These findings show that multiple resistance-associated programs spanning signaling, cell survival, and metabolism share a dependency on eIF4A-dependent translation and provide a preclinical rationale to test whether adding eIF4A inhibition can prolong responses to MAPK-targeted therapy in melanoma.

## Introduction

Activating BRAF^V600^ mutations drive constitutive mitogen-activated protein kinase (MAPK) signaling in a substantial fraction of cutaneous melanomas and provide a clinically actionable therapeutic target^1,2^. Combined BRAF and MEK inhibition produces rapid and often profound tumor responses in patients with BRAF ^V600^-mutant melanoma^3,4^. However, this benefit is not uniformly durable: pooled long-term analyses of treatment with the BRAF inhibitor dabrafenib plus the MEK inhibitor trametinib reported five-year progression-free and overall survival rates of approximately 19% and 34%, respectively^4^. Thus, even when MAPK-targeted therapy initially controls disease, residual melanoma cells can persist and adapt under treatment pressure, creating a reservoir from which recurrence can emerge^5,6^.

Acquired resistance to MAPK inhibition is genetically heterogeneous^7–11^. Recurrent mechanisms include MAPK pathway reactivation, receptor tyrosine kinase rewiring, compensatory phosphoinositide 3-kinase (PI3K)-AKT signaling, and altered survival control^7–10,12,13^. Despite this diversity, these mechanisms can converge on a more restricted set of functional outputs that restore proliferation, survival, and tumor-cell fitness under treatment pressure^14,15^. MYC activation, for example, has been identified as a common output of otherwise distinct resistance mechanisms and can create accompanying dependencies on glucose, glutamine, and serine metabolism^15^.

Resistance is increasingly understood as a staged and dynamic process rather than solely the consequence of selecting a pre-existing genetic clone^5,6,16,17^. Following MAPK inhibition, melanoma cells can enter reversible drug-tolerant or minimal-residual-disease states that provide a reservoir from which stable resistance can emerge^5,6,18^. Single-cell, spatial, proteomic, and lineage-aware studies have identified coexisting persister programs with neural crest-like, stress-adaptive, PI3K-associated, lipid-metabolic, and extracellular-matrix-remodeling features^5,16,19–21^. These programs intersect with the broader differentiation trajectory of melanoma cells, which can transition from melanocytic states characterized by microphthalmia-associated transcription factor (MITF), SRY-box transcription factor 10 (SOX10), and other melanocytic-lineage genes toward neural crest-like and undifferentiated states associated with nerve growth factor receptor (NGFR) or the receptor tyrosine kinase AXL^6,16,17,22^. This plasticity can permit melanoma cells to alter their signaling, survival, and metabolic requirements without an immediate change in genotype^5,6,16,17^.

Metabolic adaptation is another important component of resistant-cell fitness. MAPK inhibition can relieve BRAF-dependent suppression of the MITF–PGC-1α (PPARγ coactivator 1α) axis, thereby promoting mitochondrial biogenesis and oxidative phosphorylation^23^. In some models, melanoma cells with acquired BRAF inhibitor (BRAFi) resistance exhibit increased mitochondrial respiration and greater bioenergetic flexibility^24,25^. Resistant melanoma cells can also acquire increased reliance on glutamine to replenish tricarboxylic acid cycle intermediates^26^. These observations indicate that resistant-cell fitness can be maintained not only by restoration of signaling but also by coordinated remodeling of the pathways that supply energy, reducing equivalents, and biosynthetic precursors^21,23–26^.

Selective mRNA translation may provide a common regulatory layer linking adaptive signaling and survival to cell-state plasticity, extracellular-matrix remodeling, and metabolic adaptation under therapeutic stress^14,27–31^. The eukaryotic translation initiation factor 4F (eIF4F) complex, composed of the cap-binding protein eIF4E, the scaffold eIF4G, and the RNA helicase eIF4A, integrates signals from the MAPK pathway and PI3K-AKT-mechanistic target of rapamycin (mTOR) axis to regulate which mRNAs are efficiently translated^27,29,32–36^. Persistent eIF4F complex assembly and eIF4A-dependent mRNA translation have been linked to resistance or persistence under BRAF and MEK inhibition, including contexts in which MAPK signaling is restored or alternative survival pathways are engaged^14,18,37^. In melanoma models, eIF4A1 depletion reduces proliferation and alters the synthesis of cell-cycle and oxidative-phosphorylation proteins, further connecting translation initiation to tumor-cell and metabolic fitness^38^.

Rocaglate eIF4A inhibitors do not simply reproduce eIF4A depletion^39–41^. Compounds such as CR-1-31-B stabilize eIF4A on susceptible RNA sequences, often enriched in purines, thereby impeding ribosomal scanning and selectively reducing the synthesis of proteins encoded by affected mRNAs^40–43^. This molecular-clamp mechanism produces transcript-selective effects rather than an indiscriminate block of protein synthesis^40–43^. Across diverse cancer contexts, CR-1-31-B has shown preclinical antitumor activity and disrupted cell-cycle, anti-apoptotic, oncogenic-signaling, redox, and bioenergetic programs, supporting the idea that eIF4A-dependent mRNA translation couples tumor-cell survival to proliferative and metabolic homeostasis^44–49^.

Several studies establish that translation remodeling contributes to melanoma plasticity and resistance. Stress-responsive regulation of mRNA translation has been linked to the MITF-low/AXL-high melanoma state^22^. BRAFi/MEK inhibitor (MEKi)-tolerant persister cells also undergo reversible changes in translation efficiency that support their survival, and eIF4A inhibition can restrict the emergence of these persisters^18^. More recently, eIF4A-dependent synthesis of p53-binding protein 1 (53BP1) was shown to promote error-prone repair and adaptive mutability during the transition from drug tolerance to stable resistance^50^. These findings establish the importance of translation control in melanoma resistance but also sharpen the remaining question: how is established resistance organized across RNA abundance, mRNA translation, protein output, and metabolism, and which components of this acquired state remain dependent on eIF4A activity?

Here, we addressed these questions using an isogenic model of acquired BRAFi resistance, complementary melanoma models, and multi-omic analyses performed at selected post-treatment intervals across translation efficiency, polysome-associated RNA, total RNA, and protein abundance. We integrated these measurements with functional bioenergetic assays, steady-state metabolomics, isotope tracing, and *in vivo* tumor-growth studies. We show that eIF4A inhibition remains active in BRAFi-resistant melanoma cells and preferentially counters multiple programs acquired with resistance, including cell-cycle, survival, extracellular-matrix, and metabolic outputs. CR-1-31-B disrupts mitochondrial and anabolic fitness, limits the downstream incorporation of glutamine-derived carbon into glutamate and the TCA cycle, and improves tumor control when combined with BRAF inhibition *in vivo*. Together, these findings identify eIF4A-dependent mRNA translation as a convergence layer linking the resistant melanoma proteome to its metabolic phenotype and support its therapeutic targeting to constrain resistant-cell fitness.

## Results

### eIF4A inhibition suppresses growth in BRAFi-sensitive and -resistant melanoma cells

Oncogenic BRAF–MAPK signaling promotes eIF4F complex formation and selective mRNA translation^14^. We therefore asked whether direct inhibition of eIF4A, a core component of eIF4F, could suppress melanoma-cell accumulation independently of responsiveness to BRAF inhibition. We first used the matched A375/A375R model, in which A375R cells were generated from parental A375 cells by stepwise selection with increasing concentrations of the BRAFi PLX4032, up to 10 µM ^51^. Both lines carry the BRAF^V600E^ mutation. PLX4032 strongly reduced the accumulation of parental A375 cells and suppressed extracellular signal-regulated kinase 1/2 (ERK1/2) and ribosomal protein S6 (RPS6) phosphorylation (Fig. 1A,B). In contrast, A375R cells remained largely refractory across the tested concentration range, and PLX4032 had limited effects on downstream signaling, confirming the resistant phenotype (Fig. 1A,B). In A375 cells, PLX4032 produced a modest, nonsignificant reduction in puromycin incorporation, a measure of nascent protein synthesis, at 1 h, followed by a larger, significant reduction at 24 h (Fig. S1A,B), consistent with time-dependent suppression of protein synthesis downstream of BRAF-pathway inhibition. In A375R cells, the PLX4032 effect was absent at 1 h and was smaller but significant at 24 h (Fig. S1A,B). As a second model, we used another independently derived BRAFi-resistant melanoma line, LU1205R, selected from BRAF^V600E^-mutant LU1205 cells with increasing concentrations of up to 5 µM PLX4032^51^. In LU1205R cells, PLX4032 similarly produced limited effects on cell accumulation, ERK1/2 and RPS6 phosphorylation, and puromycin incorporation (Fig. S1C,D).

**Fig. 1.**
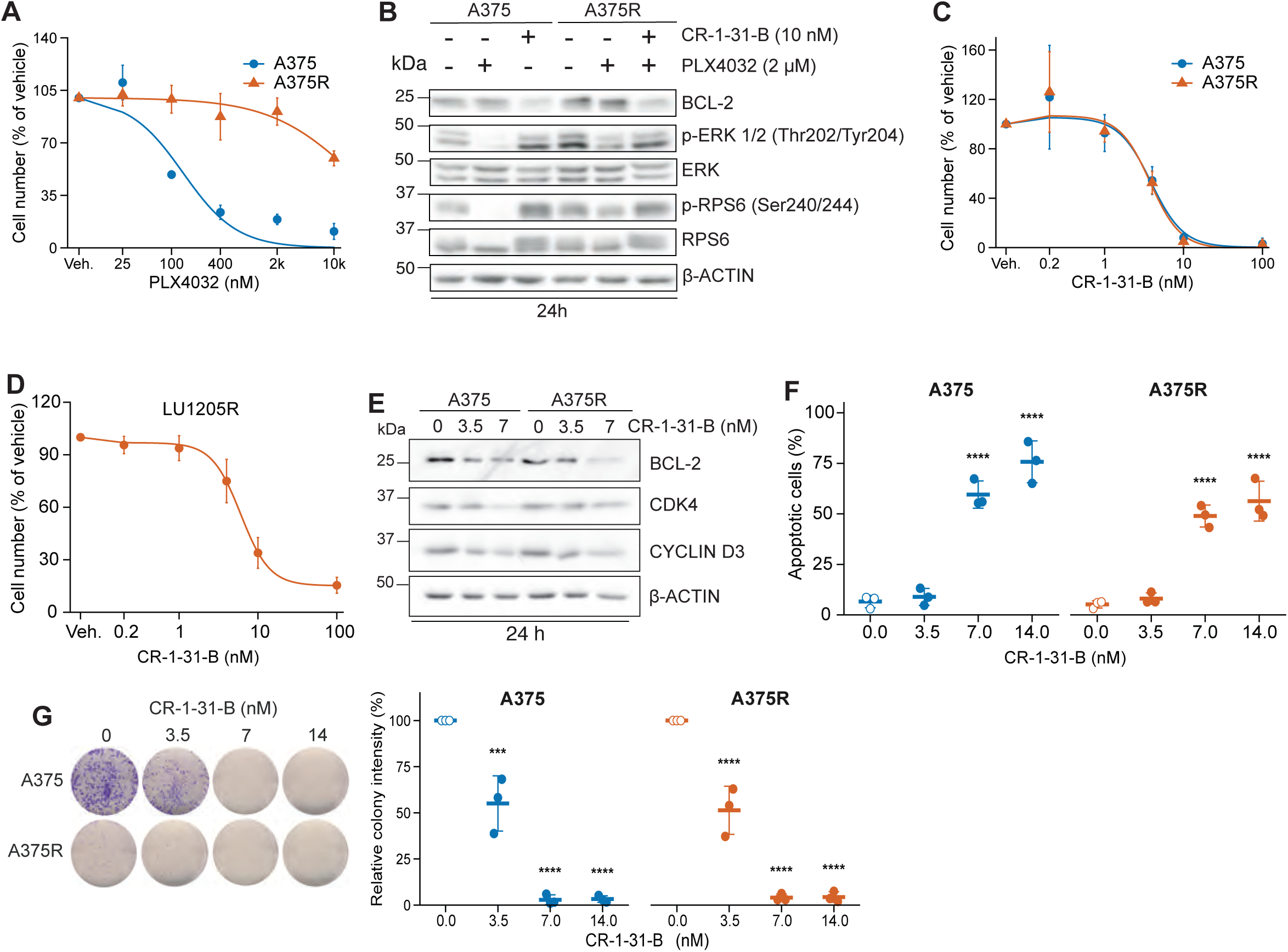
eIF4A inhibition suppresses growth and survival in BRAFi-sensitive and -resistant melanoma cells. **(A)** Representative concentration-response experiment in which A375 and A375R cells were exposed to PLX4032 for 72 h. Viable cells were counted by trypan-blue exclusion and expressed relative to vehicle-treated cells. Across three independently fitted experiments, the mean PLX4032 IC_50_ for A375 cells was 124 ± 27 nM (mean ± SD); the A375R IC_50_ was not reached within the tested range and is reported as >10 µM. **(B)** Immunoblots of indicated proteins after 24 h: A375, vehicle, PLX4032 (2 µM), or CR-1-31-B (10 nM); A375R, vehicle, PLX4032, or PLX4032 + CR-1-31-B at the same concentrations. β-actin was the loading control. **(C,D)** Representative CR-1-31-B concentration-response curves in A375 and A375R cells (C) and LU1205R cells (D) after 72 h. Cell numbers were normalized to the corresponding vehicle control. **(E)** Immunoblot analysis of BCL-2, CDK4, and cyclin D3 after 24 h exposure to increasing CR-1-31-B concentrations. **(F)** Total Annexin V-positive cells (Annexin V+/DAPI− plus Annexin V+/DAPI+) after 72 h exposure to the indicated CR-1-31-B concentrations, quantified by flow cytometry. **(G)** Representative clonogenic images and corresponding crystal-violet quantification after 72 h exposure to the indicated CR-1-31-B concentrations, followed by replating of 1,000 viable cells per well and 6 days of drug-free colony growth. Colony intensity was normalized to vehicle-treated cells. In F and G, symbols represent three independent biological experiments and bars show mean ± SD. Statistical comparisons were performed by one-way ANOVA followed by Tukey’s honestly significant difference test within each cell line; displayed comparisons are dose versus vehicle. ***P < 0.001; ****P < 0.0001.

To investigate the effect of eIF4A inhibition, we next tested a panel of four eIF4A inhibitors—CR-1-31-B, SDS-1-021^52^, silvestrol^53^ and CMLD012612^54^—in A375, A375R and LU1205R cells. All four compounds inhibited cell accumulation at low-nanomolar concentrations and retained activity in the BRAFi-resistant models, indicating that this response extended across the eIF4A-inhibitor panel (Fig. 1C,D; Fig. S1E-K). Because CR-1-31-B was the most extensively characterized compound in the panel and had established utility in both cell-based and *in vivo* studies, we selected it for subsequent mechanistic and therapeutic experiments. CR-1-31-B was also active in YUMMER1.7 murine melanoma cells, extending this observation to a Braf^V600E^-driven murine model (Fig. S1K,L). Thus, melanoma cells remain sensitive to eIF4A inhibition regardless of their responsiveness to BRAFi across the models tested.

We then examined whether this pharmacological response was accompanied by inhibition of translation-dependent molecular outputs. After 1 h, CR-1-31-B alone reduced puromycin incorporation relative to vehicle in A375 cells, whereas adding CR-1-31-B to PLX4032 reduced incorporation relative to PLX4032 alone in A375R cells, demonstrating rapid inhibition of nascent protein synthesis before overt changes in cell accumulation (Fig. S1A). We selected anti-apoptotic protein BCL-2, cyclin-dependent kinase 4 (CDK4) and cyclin D3 as established translation-sensitive outputs of mTOR complex 1 (mTORC1)/eIF4F signaling. Dose-response immunoblots showed that CR-1-31-B reduced BCL-2, CDK4, and cyclin D3 abundance in both A375 and A375R cells (Fig. 1E). Similar molecular changes were observed with SDS-1-021 and CMLD012612 in A375/A375R cells and across multiple eIF4A inhibitors in LU1205R cells (Fig. S1M-O).

We next determined whether these early molecular effects translated into loss of tumor-cell survival and long-term proliferative capacity. CR-1-31-B dose-dependently increased apoptosis and suppressed colony formation in both A375 and A375R cells (Fig. 1F,G).

To examine whether this response extended beyond BRAF^V600^ melanoma, we tested MDA-MB-231 breast cancer cells carrying the non-V600 BRAF^G464V^ mutation and a matched derivative selected for resistance to combined BRAF and MEK inhibition with encorafenib and binimetinib, respectively. Despite their different responsiveness to MAPK inhibition (Fig. S1P,Q), both cell lines were sensitive to CR-1-31-B, which reduced clonogenic growth and the abundance of BCL-2, CDK4 and cyclin D3 (Fig. S1R-T). Together, these results show that direct inhibition of eIF4A suppresses proliferative and survival outputs downstream of oncogenic signaling and retains activity across models with acquired resistance to BRAF/MAPK inhibition.

### Acquired BRAFi resistance is associated with coordinated transcriptomic, translational, proteomic, and bioenergetic remodeling

To define the molecular programs that sustain the A375R state and might explain its retained sensitivity to eIF4A inhibition, we compared A375R with parental A375 cells using matched sequencing of total and polysome-associated RNA analyzed with Anota2Seq^55^, together with quantitative proteomics (Fig. 2A). Total RNA measures transcript availability, whereas polysome-associated RNA represents transcripts engaged by multiple ribosomes and therefore reflects both transcript abundance and ribosome loading. Anota2Seq models the relationship between these two RNA pools to distinguish three regulatory modes: abundance-mode regulation, in which total and polysome-associated RNA change concordantly; translational-efficiency (TE) regulation, in which polysome association changes beyond that expected from total-RNA abundance; and translational buffering, in which a change in total RNA is offset by an opposing change in TE, limiting its propagation to the polysome-associated pool^55^. The polysome profiling analyses included six biological replicates per cell state across two experimental batches, and proteomics included four biological replicates per cell state. Baseline polysome profiles were broadly comparable, although A375R cells had a modestly higher polysome/80S ratio (1.25-fold; P = 0.034; Fig. S2A,B). Principal-component analyses (PCA) separated A375 from A375R across total RNA, polysome-associated RNA, TE, and protein-abundance datasets, indicating reproducible remodeling at each layer (Fig. S2C-F).

**Fig. 2.**
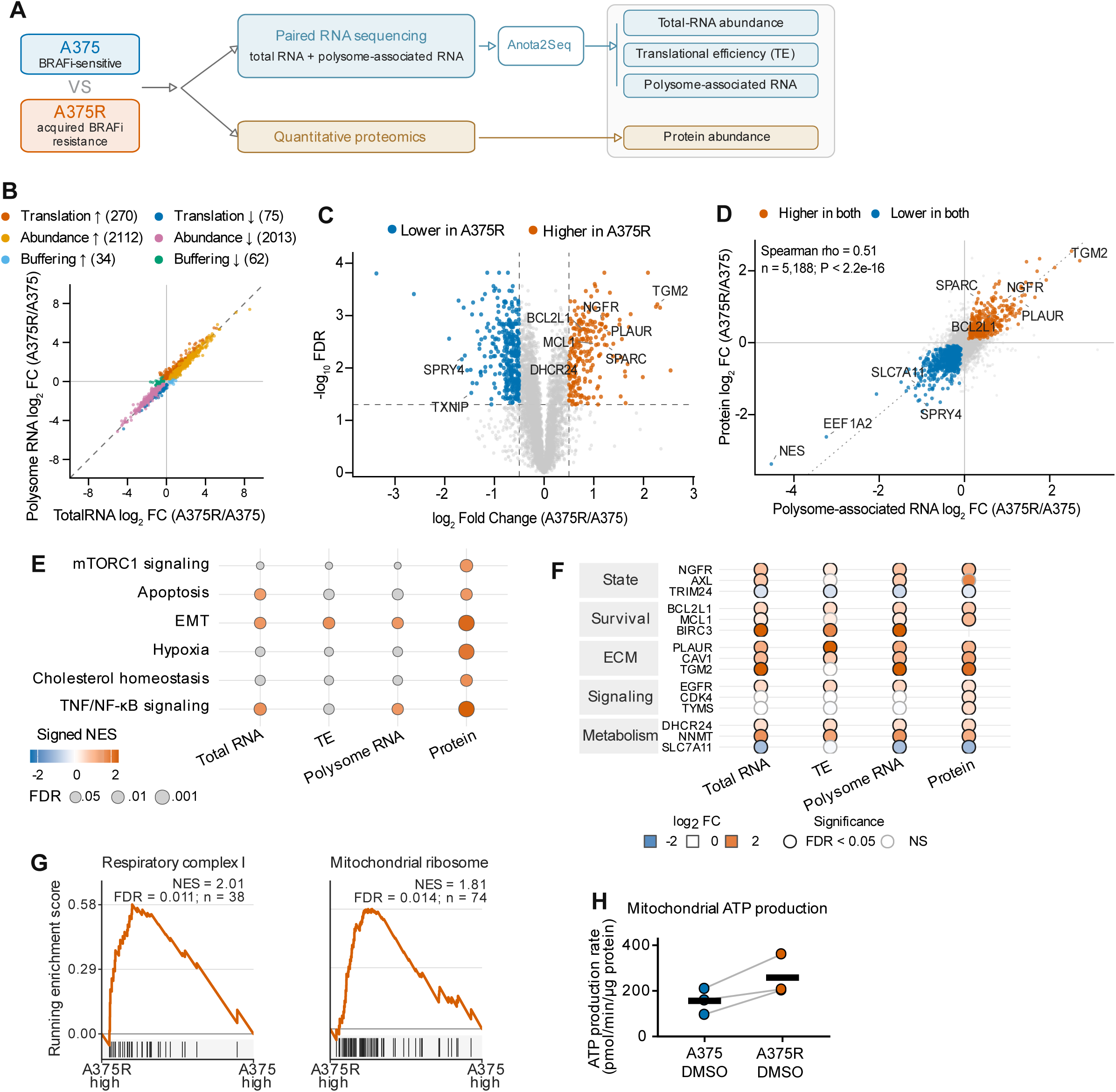
Acquired BRAFi resistance is associated with multilayered transcriptomic, translational, proteomic, and bioenergetic remodeling. **(A)** Experimental design for the basal-state comparison of parental BRAFi-sensitive A375 cells and BRAFi-resistant A375R cells generated by chronic PLX4032 selection. Paired total and polysome-associated RNA sequencing was analyzed with Anota2Seq to resolve total-RNA abundance, translational efficiency (TE), and polysome-associated RNA changes; quantitative proteomics measured protein abundance. RNA analyses included six biological replicates per cell state across two experimental batches, and proteomics included four biological replicates per cell state. **(B)** Anota2Seq regulatory-mode plot showing total-RNA versus polysome-associated RNA log_2_ fold changes in A375R relative to A375. Colored transcripts were assigned to translation, abundance, or buffering modes using the predefined FDR and effect-size criteria described in Methods; numbers indicate transcripts in each directional class. **(C)** Proteomic volcano plot for A375R versus A375. Colored proteins met FDR < 0.05 and |log_2_ fold change| ≥ 0.5; selected melanoma-relevant proteins are labeled. **(D)** Concordance between polysome-associated RNA and protein changes across 5,188 matched genes. Colored genes were significant in both layers and changed concordantly; the dotted line denotes equal fold change. Spearman ρ = 0.51, P < 2.2 × 10⁻¹⁶. **(E)** Hallmark gene-set enrichment across total RNA, TE, polysome-associated RNA, and protein. Color denotes signed normalized enrichment score (NES), dot size denotes -log_10_ FDR, and gray denotes a tested but nonsignificant gene set. **(F)** Selected genes associated with melanoma state, survival, extracellular matrix (ECM), signaling, and metabolism. Fill denotes log_2_ fold change in A375R relative to A375; black outlines denote FDR < 0.05, gray outlines denote nonsignificant changes, and x denotes not detected. **(G)** MitoCarta 3.0 preranked GSEA of the A375R-versus-A375 proteome for respiratory complex I and mitochondrial-ribosome gene sets. Proteins were ranked by limma log_2_ fold change; positive enrichment indicates higher abundance in A375R. (H) Mitochondrial ATP-production rates in vehicle-treated A375 and A375R cells measured using the Seahorse ATP Rate Assay. Lines connect paired biological experiments (n = 3); black bars indicate means.

Anota2Seq showed that established resistance was dominated by abundance-mode regulation: total and polysome-associated RNA increased concordantly for 2,112 transcripts and decreased concordantly for 2,013 transcripts. Nevertheless, 270 transcripts showed increased TE and 75 showed decreased TE, identifying changes in polysome engagement that were not explained by corresponding changes in transcript abundance. Additional transcripts exhibited translational buffering, indicating that some total-RNA changes were counteracted at the level of polysome association (Fig. 2B; Fig. S2G). Quantitative proteomics identified broad remodeling of the resistant-cell proteome, including increased NGFR, associated with neural crest-like melanoma states, and increased PLAUR, SPARC, and TGM2, proteins linked to extracellular-matrix remodeling, adhesion, and invasive plasticity, together with altered abundance of survival and metabolic proteins (Fig. 2C). Thus, the resistant state combines a broad abundance-driven RNA program with a smaller but distinct contribution from selective translational regulation and buffering, with both components reflected in altered protein output. Complete resistance-state multi-omic results are provided in Supplementary Table S1.

To determine how changes in polysome-associated RNA relate to protein abundance, we compared fold changes at these two levels across 5,188 genes quantified in both datasets (Fig. 2D). The two layers were positively correlated (Spearman ρ = 0.51, P < 2.2 × 10⁻^16^), and concordantly increased genes included NGFR (neural crest-like state), PLAUR and SPARC (extracellular-matrix remodeling and adhesion), and BCL2L1 (anti-apoptotic survival) (Fig. 2D). TE changes were also correlated with protein abundance, although more modestly (Spearman ρ = 0.45, P < 2.2 × 10⁻^16^; Fig. S2H). Thus, polysome-associated RNA and TE provide complementary information about how regulatory changes beyond total-RNA abundance are reflected in protein output.

Pathway-level analysis revealed convergence on programs previously implicated in melanoma persistence and therapeutic adaptation, including phenotype switching and survival rewiring^5,6,17,56^ (Fig. 2E). Epithelial–mesenchymal-transition (EMT) signatures were enriched across several regulatory layers, whereas hypoxia, mTORC1 signaling, apoptosis/survival, cholesterol homeostasis and TNF/NF-κB signaling showed layer-selective enrichment, particularly at the protein level^25,57^ (Fig. 2E). Gene-level inspection supported this broader organization: melanoma-state regulators such as NGFR and AXL were altered at one or more layers; BCL2L1, MCL1 and BIRC3 marked survival rewiring; PLAUR, CAV1 and TGM2 reflected extracellular-matrix and adhesion remodeling; and EGFR, CDK4 and TYMS represented signaling and proliferative outputs (Fig. 2F). Metabolic changes were similarly heterogeneous, including increased DHCR24, an enzyme involved in cholesterol biosynthesis, and NNMT, a nicotinamide methyltransferase, together with reduced SLC7A11, the substrate-specific subunit of the cystine/glutamate antiporter. The absence of uniform regulation across all layers argues against a single linear resistance pathway and instead supports a coordinated state assembled through several regulatory routes.

Mitochondrial remodeling emerged as a particularly coherent feature of acquired BRAFi resistance. Respiratory-complex-I and mitochondrial-ribosome proteins were positively enriched in the A375R proteome (NES = 2.01 and 1.81, respectively; Fig. 2G). Consistent with this proteomic signature, mitochondrial ATP-production rates were higher in A375R cells in each of three paired experiments (Fig. 2H). Mito Fuel Flex analysis further revealed increased glucose-oxidation capacity in A375R cells, accompanied by reduced capacities for fatty-acid and glutamine oxidation (Fig. S2I,J). These observations indicate that acquired resistance is associated not simply with globally increased substrate flexibility, but with a redistribution toward greater glucose-supported mitochondrial capacity.

Together, these data characterize the acquired BRAFi-resistant state in A375R cells as multilayered, with extensive RNA-abundance changes reinforced by selective translational regulation and reflected in coordinated proteomic and bioenergetic outputs. Given this multilayered remodeling and the vulnerability of A375R cells to eIF4A inhibition, we next examined how eIF4A inhibition affected these regulatory layers at selected post-treatment intervals.

### eIF4A inhibition produces distinct translational, RNA-abundance and proteomic responses in BRAFi-resistant melanoma

Having defined the multilayered programs associated with acquired BRAFi resistance in A375R cells, we next asked how eIF4A inhibition remodels this state across translational, RNA-abundance and protein layers. A375R cells were treated with CR-1-31-B (10 nM) or DMSO and analyzed by paired total and polysome-associated RNA sequencing at 1 h, independent total RNA sequencing at 24 h, and quantitative proteomics at 36 h (n = 6, n = 6 and n = 4 biological replicates per condition, respectively; Fig. 3A). Representative polysome profiles from one of the two experimental batches (three of six biological replicates per condition) showed relative accumulation of 80S monosomes and marked depletion of heavy polysomes after CR-1-31-B treatment (Fig. S3A). Across both batches, quantification confirmed a lower polysome/80S ratio following CR-1-31-B treatment (Fig. S3B; P < 0.001). Because CR-1-31-B directly targets eIF4A-dependent mRNA translation, the 24-h total RNA-seq layer was used to capture secondary RNA-abundance remodeling downstream of the early translational response, whereas the 36-h proteome represents the integrated molecular output. PCA showed that the 1-h input-RNA and polysome-associated-RNA samples separated by treatment (Fig. S3C,D). To assess whether the inferred translational-efficiency changes were robust to the analytical framework, we compared the Anota2Seq estimates with a DESeq2 interaction analysis. The two approaches were strongly concordant (Spearman ρ = 0.88 across all genes and 0.95 among Anota2Seq-selected genes), and 2,214 of 2,678 Anota2Seq-selected transcripts were also significant by DESeq2 (Fig. S3E). Complete 1-h CR-1-31-B-response multi-omic results are provided in Supplementary Table S2.

**Fig. 3.**
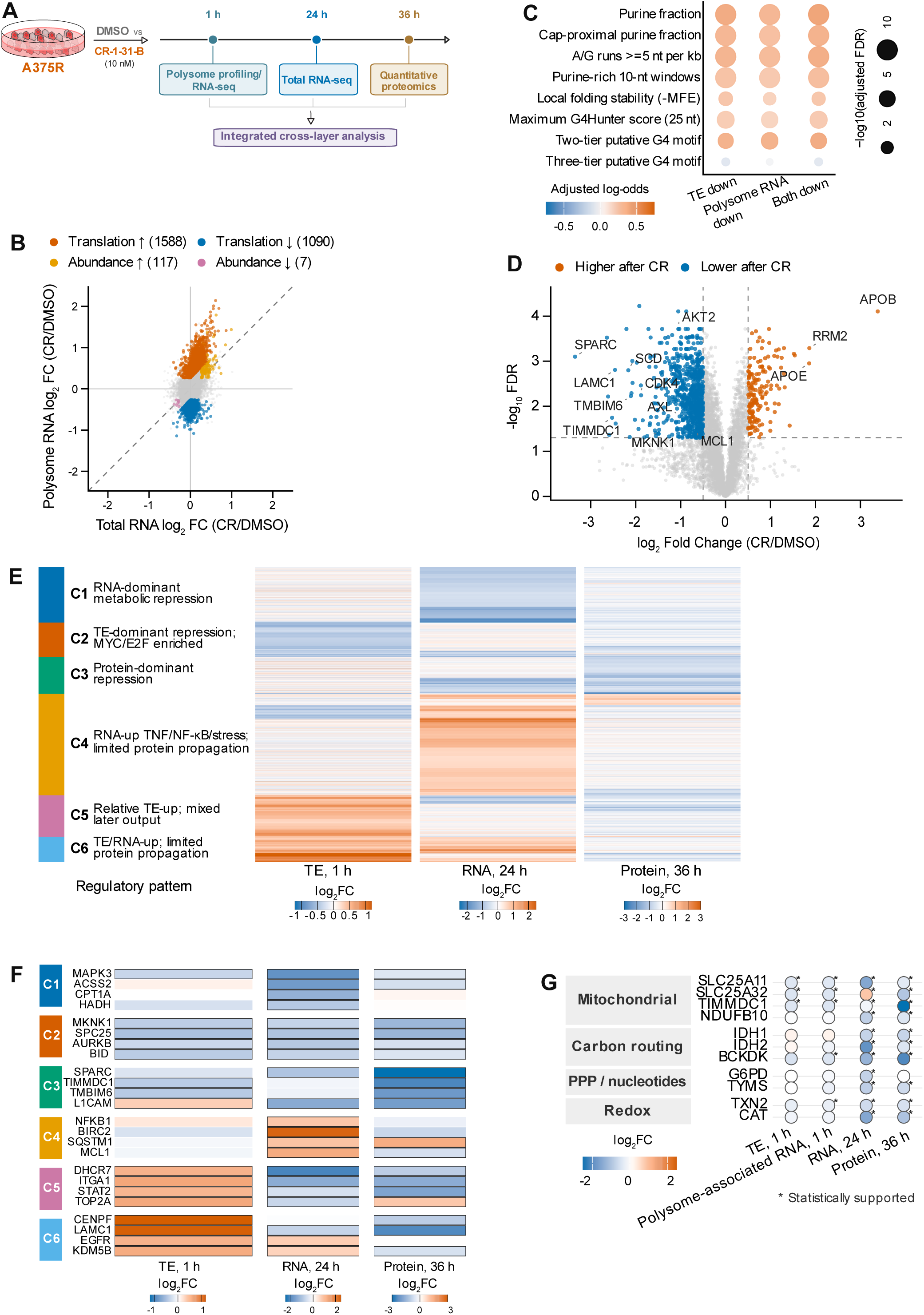
CR-1-31-B elicits distinct translational, RNA-abundance, and proteomic responses in A375R cells. **(A)** Experimental design. A375R cells were treated with CR-1-31-B (10 nM) or DMSO and analyzed by paired input/polysome-associated RNA sequencing at 1 h, total RNA sequencing at 24 h, and quantitative proteomics at 36 h. The respective sample sizes were n = 6, n = 6, and n = 4 biological replicates per condition. **(B)** Anota2Seq regulatory-mode plot showing total-RNA versus polysome-associated RNA log_2_ fold changes after 1 h CR-1-31-B treatment. Colored transcripts were classified as translation- or abundance-regulated using the predefined FDR and effect-size criteria described in Methods; numbers indicate transcripts in each directional class. **(C)** Covariate-adjusted associations between selected features of one representative, publicly annotated 5′UTR per gene and transcript classes showing decreased TE, decreased polysome-associated RNA, or concordant decreases at both levels. Color denotes adjusted log-odds for membership in the corresponding CR-1-31-B-down class: orange indicates a positive association or enrichment and blue indicates a negative association or depletion. Point size denotes −log_10_(Benjamini–Hochberg FDR), capped at 10 for display. The local-folding coefficient was oriented so that orange indicates stronger predicted stability. **(D)** Proteomic volcano plot at 36 h. Colored proteins met FDR < 0.05 and |log_2_ fold change| ≥ 0.5; selected signaling, survival, extracellular-matrix, mitochondrial, and metabolic proteins are labeled. **(E)** Heatmap of 3,241 genes quantified across all three layers and meeting CR-response criteria in at least one layer. Hierarchical clustering grouped genes into six response patterns (C1–C6); colors show log₂ fold changes using layer-specific scales. **(F)** Representative genes from each cluster in E. Values are log_2_ fold changes across TE, total-RNA, and protein layers. **(G)** Selected mitochondrial, carbon-routing, pentose-phosphate/nucleotide, and redox genes across TE, polysome-associated RNA, total RNA, and protein. Fill denotes log_2_ fold change after CR-1-31-B treatment; asterisks indicate statistically supported effects in the corresponding layer.

At 1 h, CR-1-31-B predominantly altered translational efficiency: 1,588 transcripts showed increased and 1,090 showed decreased translational efficiency, whereas only 117 and 7 transcripts were assigned to increased and decreased abundance modes, respectively (Fig. 3B). Because TE effects are relative, the TE-increased class represents transcripts whose polysome association was preferentially maintained or increased during CR-1-31-B treatment, rather than increased global protein synthesis. Thus, before extensive remodeling of RNA abundance, eIF4A inhibition rapidly redistributed polysome association across a large transcript set. Pathway analysis indicated that the TE-decreased group included MYC targets and fatty-acid, amino-acid and translation-related programs, whereas TE-increased transcripts were enriched for cell-cycle, integrin/extracellular-matrix and chromatin-related programs (Fig. S3F). These findings support transcript-selective translational reprogramming rather than uniform suppression of translation across all mRNAs.

Because transcript sensitivity to eIF4A can be influenced by 5′ untranslated region (5′UTR) sequence architecture, we examined publicly annotated representative 5′UTRs of CR-1-31-B-responsive transcripts. Transcripts with decreased TE and/or polysome-associated RNA were associated with purine-rich, locally structured 5′UTRs, including greater cap-proximal purine content, longer purine tracts, and stronger predicted local folding stability (Fig. 3C; Fig. S3G,H). Predicted G4 features contributed to this association but were not consistently supported across definitions, indicating that G4 potential is one component of a broader sequence architecture. Because public transcript annotations were used, this analysis identifies sequence propensities and does not define the exact 5′UTR isoforms expressed in A375R cells. Complete adjusted 5′UTR feature-association results are provided in Supplementary Table S3A; the corresponding complete gene-level 1-h CR-1-31-B response data and representative-transcript 5′UTR annotations are provided in Supplementary Table S3B.

For the 36 h proteomic analysis, PCA showed that the quantitative proteomes separated by both cell state and CR-1-31-B exposure (Fig. S3I). The proteomic response was more strongly weighted toward reduced protein abundance: 752 proteins decreased and 162 increased using the predefined significance and effect-size thresholds (Fig. 3D). Reduced proteins included signaling and proliferation regulators (AKT2, AXL, CDK4 and MKNK1), survival proteins (MCL1 and TMBIM6), extracellular-matrix components (SPARC and LAMC1), and mitochondrial or metabolic proteins (TIMMDC1 and SCD). A smaller set increased, including APOB, APOE and RRM2. Consistent with these gene-level effects, protein-level GSEA showed depletion of PI3K-AKT-mTOR, receptor-tyrosine-kinase/GPCR and extracellular-matrix programs. Conversely, enrichment of selected MYC and mRNA-translation-related protein sets indicated possible compensatory responses (Fig. S3J). Thus, the later proteome reflected broad but non-uniform suppression of melanoma-fitness programs.

To resolve how early translational regulation and later RNA-abundance changes combine to shape protein output, we clustered 3,241 CR-1-31-B-responsive genes quantified across the TE, 24-h total-RNA and 36-h protein layers. Six regulatory patterns emerged (Fig. 3E,F). These included RNA-dominant metabolic repression (C1; exemplified by MAPK3, ACSS2, CPT1A and HADH), TE-dominant repression enriched for MYC/E2F-linked programs (C2; MKNK1, SPC25, AURKB and BID), and protein-dominant repression (C3; SPARC, TIMMDC1, TMBIM6 and L1CAM). A distinct cluster showed increased stress- and TNF/NF-κB-linked RNA abundance with limited propagation to the proteome (C4; NFKB1, BIRC2 and SQSTM1), while two clusters showed relative TE increases with mixed or limited later protein output (C5-C6). These patterns show that changes at one regulatory layer do not necessarily predict the final protein response: CR-1-31-B-sensitive protein outputs can arise through early selective translation, later RNA-abundance remodeling, or protein-dominant regulation.

A focused analysis further identified coordinated effects on metabolic control points spanning mitochondrial transport and respiratory function (SLC25A11, SLC25A32, TIMMDC1 and NDUFB10), carbon routing (IDH1, IDH2 and BCKDK), pentose-phosphate/nucleotide metabolism (G6PD and TYMS), and redox control (TXN2 and CAT) (Fig. 3G). For several genes, reduced TE or polysome-associated RNA was already evident at 1 h and was accompanied by reduced RNA abundance and/or protein abundance at later times. Other changes emerged predominantly at the RNA or protein layer, and not every RNA decrease propagated to protein abundance, as illustrated by G6PD (Fig. 3G). Together, these data indicate that eIF4A inhibition initiates an early transcript-selective response that is reinforced and reshaped across regulatory layers, ultimately constraining multiple signaling, survival and metabolic programs in BRAFi-resistant cells.

### eIF4A inhibition preferentially suppresses proteins acquired during BRAFi resistance

Because several proteins suppressed by CR-1-31-B were elevated in A375R cells, we asked whether eIF4A inhibition preferentially opposes proteomic changes acquired during chronic BRAFi selection. We aligned the A375R-versus-A375 proteomic comparison with the CR-1-31-B-versus-DMSO response in A375R cells. Of the 225 proteins significantly increased in A375R relative to A375, 65 (28.9%) were significantly reduced by CR-1-31-B (FDR < 0.05; log2FC ≤ −0.5). More broadly, 168 (74.7%) showed a negative CR-1-31-B-associated fold change, indicating a strong directional bias toward suppression of resistance-up proteins. By contrast, only 92 of 346 resistance-down proteins (26.6%) shifted upward after CR-1-31-B, of which 13 (3.8%) met the strict threshold (Fig. 4A; Fig. S4A). This marked asymmetry indicates that CR-1-31-B preferentially suppresses proteins acquired in the resistant state.

**Fig. 4.**
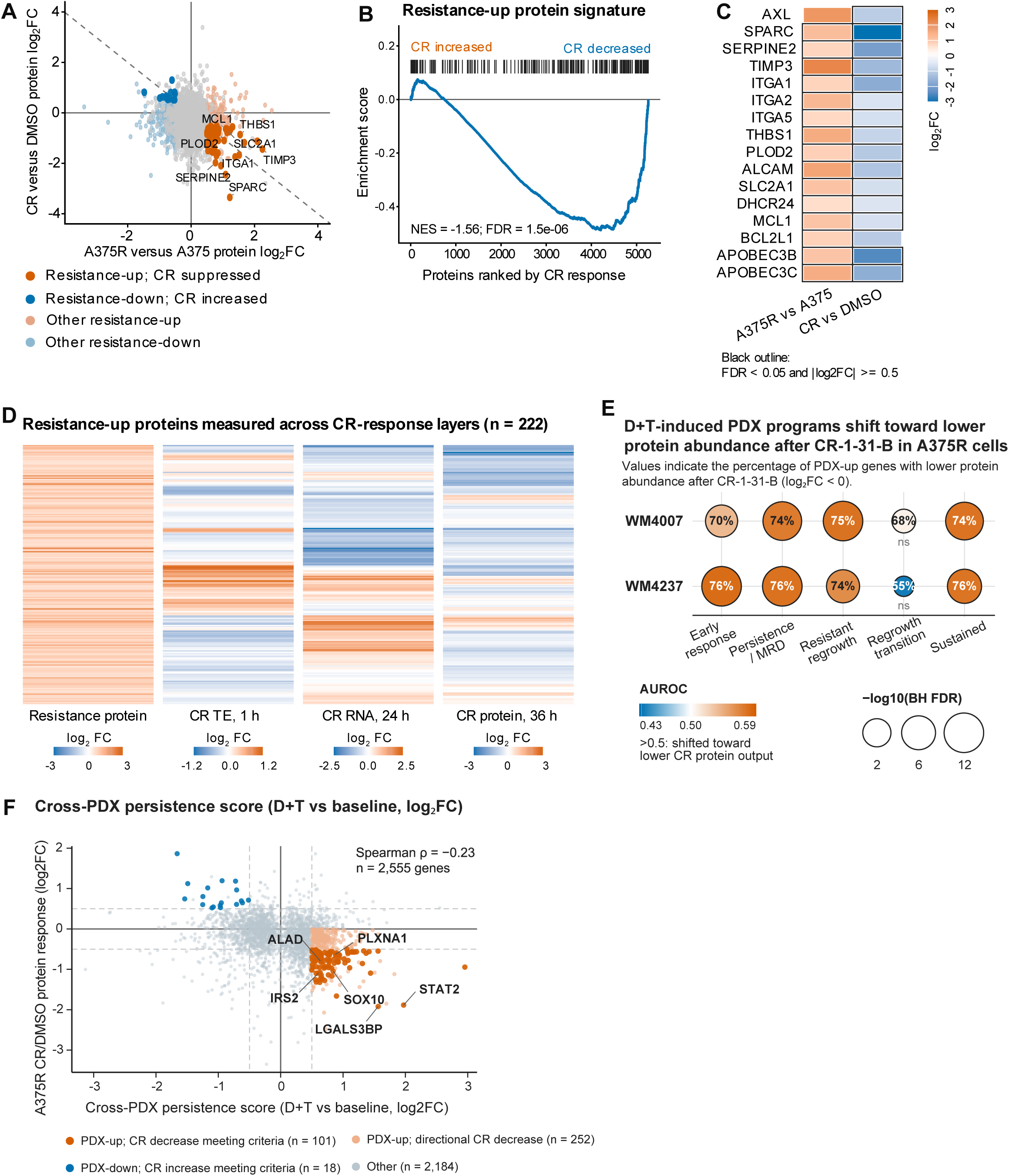
CR-1-31-B preferentially suppresses proteins acquired during BRAFi resistance. **(A)** Protein log_2_ fold changes in A375R relative to A375 are plotted against protein changes in A375R cells treated with CR-1-31-B versus DMSO. Resistance-associated and CR-1-31-B-responsive proteins were defined by FDR < 0.05 and |log_2_ fold change| ≥ 0.5. The dashed line denotes equal-magnitude opposition (y = −x); selected resistance-up proteins shifted downward by CR-1-31-B treatment are labeled. **(B)** Preranked GSEA of the 225-protein resistance-up signature against the complete ranked CR-1-31-B-responsive proteome. Proteins are ordered from CR-1-31-B-increased to CR-1-31-B-decreased, and vertical marks identify members of the resistance-up set. **(C)** Selected proteins associated with melanoma state/plasticity, ECM, adhesion, survival, glucose transport, lipid metabolism, and APOBEC-family proteins linked to mutagenesis. Fill denotes log_2_ fold change; black outlines denote FDR < 0.05 and |log_2_ fold change| ≥ 0.5 within the corresponding comparison. **(D)** The 222 resistance-up proteins quantified across the CR-1-31-B-response layers, shown for resistance-associated protein abundance and CR-1-31-B effects on TE at 1 h, total RNA at 24 h, and protein abundance at 36 h. Rows were grouped by similarity across CR-response layers; colors show raw effects using layer-specific scales. **(E)** Stage-resolved rank enrichment of genes induced in each PDX during continuous dabrafenib plus trametinib (D+T) treatment. PDX programs were defined independently in WM4007 and WM4237 and evaluated against the ranked A375R CR-1-31-B-responsive proteome. Color denotes rank enrichment (AUROC), where 0.5 indicates no shift and values > 0.5 indicate a shift toward lower protein abundance after CR-1-31-B treatment. Dot size denotes −log_10_(Benjamini–Hochberg-adjusted FDR), inset numbers show the percentage of PDX-up genes with CR-1-31-B protein log_2_ fold change < 0, and ns denotes Benjamini–Hochberg-adjusted FDR ≥ 0.05. **(F)** Relationship between the cross-PDX persistence score and the 36-h A375R protein response to CR-1-31-B across 2,555 matched genes. The score captures the shared direction and smaller absolute log₂ fold change across WM4007 and WM4237. Orange identifies persistence-induced genes whose proteins decreased after CR-1-31-B, blue identifies persistence-decreased genes whose proteins increased, and darker colors indicate effects meeting the predefined significance and fold-change criteria; category sizes are shown in the panel. Dashed lines mark |log₂ fold change| = 0.5. Spearman ρ = −0.23, P = 4.75 × 10⁻³¹. Selected genes are labeled. PDX analyses are descriptive because one tumor-bearing mouse was profiled per model and stage.

Consistent with this interpretation, the complete 225-protein resistance-up signature was negatively enriched across the ranked CR-1-31-B-responsive proteome (NES = −1.56; FDR = 1.5 × 10^-6^), indicating a collective shift of resistance-acquired proteins toward lower abundance after CR-1-31-B treatment (Fig. 4B). The proteins displayed in Fig. 4C span melanoma-state and extracellular-matrix regulation (AXL, SPARC, SERPINE2, TIMP3, THBS1 and PLOD2), integrin/adhesion signaling (ITGA1, ITGA2, ITGA5 and ALCAM), survival (MCL1 and BCL2L1), metabolic support (SLC2A1 and DHCR24), and APOBEC-family proteins linked to mutagenesis (APOBEC3B and APOBEC3C). The panel illustrates the functional breadth of resistance-associated proteins that shifted downward after CR-1-31-B. Pathway analysis of the strict 65-protein resistance-up/CR-1-31-B-down subset reinforced this organization. The suppressed set was enriched for EMT and coagulation signatures, together with Reactome pathways related to extracellular-matrix (ECM) organization, ECM proteoglycans, integrin and non-integrin cell-surface interactions, and L1CAM interactions (Fig. S4B). Immunoblotting confirmed the resistance-up/CR-down pattern for AXL: its abundance was markedly higher in A375R than A375 cells and decreased after 24 h of CR-1-31-B exposure (Fig. S4C). These findings identify ECM, adhesion and state-plasticity programs as prominent components of the resistance-acquired proteome opposed by eIF4A inhibition.

Having established protein-level opposition, we next mapped CR-1-31-B regulation of the resistance-acquired set across molecular layers. Of the 225 proteins increased in A375R cells, 222 could be linked to corresponding CR-response measurements of TE at 1 h, total RNA at 24 h, and protein abundance at 36 h (Fig. 4D). Many were reduced at one or more layers, but the pattern varied among genes, with suppression evident in TE at 1 h, total-RNA abundance at 24 h, protein abundance at 36 h, or combinations of these layers—consistent with the multilayered response defined in Fig. 3. These data demonstrate preferential suppression of a coordinated resistance-acquired module enriched for melanoma plasticity, extracellular-matrix/adhesion, survival and metabolic functions. Complete resistance-program analyses are provided in Supplementary Table S4.

To determine whether the CR-1-31-B-responsive proteome identified in A375R cells was relevant to treatment-associated states in patient-derived models, we analyzed two publicly available BRAF^V600E^ melanoma PDX spatial-transcriptomic trajectories profiled during continuous dabrafenib/trametinib treatment^20^ (Fig. S4D). Stage-specific programs were defined independently in WM4007 and WM4237 and evaluated against the ranked A375R CR-1-31-B-responsive proteome. In each PDX model, genes induced during early response, persistence, or resistant regrowth—and genes whose induction was sustained from persistence through regrowth—were shifted toward lower protein abundance after CR-1-31-B in A375R cells (Fig. 4E). Across these programs, 70–76% of induced genes had a negative CR-1-31-B-associated protein fold change, and each showed significant rank enrichment after Benjamini–Hochberg correction. By contrast, genes newly induced during the transition from persistence to regrowth were not significantly shifted in either model, indicating stage-selective rather than uniform opposition.

The strongest convergence occurred during persistence: the cross-PDX persistence score was inversely correlated with the 36-h A375R CR-1-31-B response (Spearman ρ = −0.23, P = 4.75 × 10⁻³¹; Fig. 4F). Of 467 genes induced during persistence in both PDXs, 353 (75.6%) showed lower protein abundance after CR-1-31-B in A375R cells, including 101 (21.6%) that met the stringent protein-down criterion. Representative genes labeled in Fig. 4F included STAT2, LGALS3BP, IRS2, SOX10, PLXNA1, and ALAD; a broader curated set spanning signaling/stress survival, melanoma-state/adhesion, metabolism, and cell-cycle/regrowth functions is shown in Fig. S4E. These cross-model results support a broad inverse rank shift between programs emerging during BRAF/MEK inhibition and the A375R protein response to CR-1-31-B, rather than complete reversal of a fixed resistance signature. We therefore next asked whether one recurrently affected program—bioenergetics and carbon metabolism—showed corresponding functional alterations.

### eIF4A inhibition constrains bioenergetic and glutamine-carbon metabolism in BRAFi-sensitive and -resistant melanoma cells

The integrated analyses identified coordinated effects on mitochondrial, carbon-handling and nucleotide-entry programs, prompting us to determine whether this regulatory remodeling produced a functional metabolic phenotype (Fig. 5A). A375 and A375R cells were exposed for 24 h to DMSO, PLX4032, CR-1-31-B, or their combination and analyzed by Seahorse bioenergetic profiling. In BRAFi-sensitive A375 cells, PLX4032 reduced energetic output, predominantly through suppression of glycolytic ATP production (Fig. 5B). In A375R cells, which displayed higher baseline glycolytic and mitochondrial ATP-production rates than A375 cells, PLX4032 had little effect. By contrast, CR-1-31-B-containing treatments reduced bioenergetic output in both cell states and shifted A375R cells toward a low-output metabolic state (Fig. 5B). Complete extracellular-acidification and oxygen-consumption profiles confirmed broad suppression of glycolytic and respiratory parameters by CR-1-31-B (Fig. S5A,B). Consistent with this functional response, OXPHOS programs were negatively enriched in polysome-associated RNA at 1 h and total RNA at 24 h, with particularly strong depletion of complex-I genes at 24 h (Fig. S5C).

**Fig. 5.**
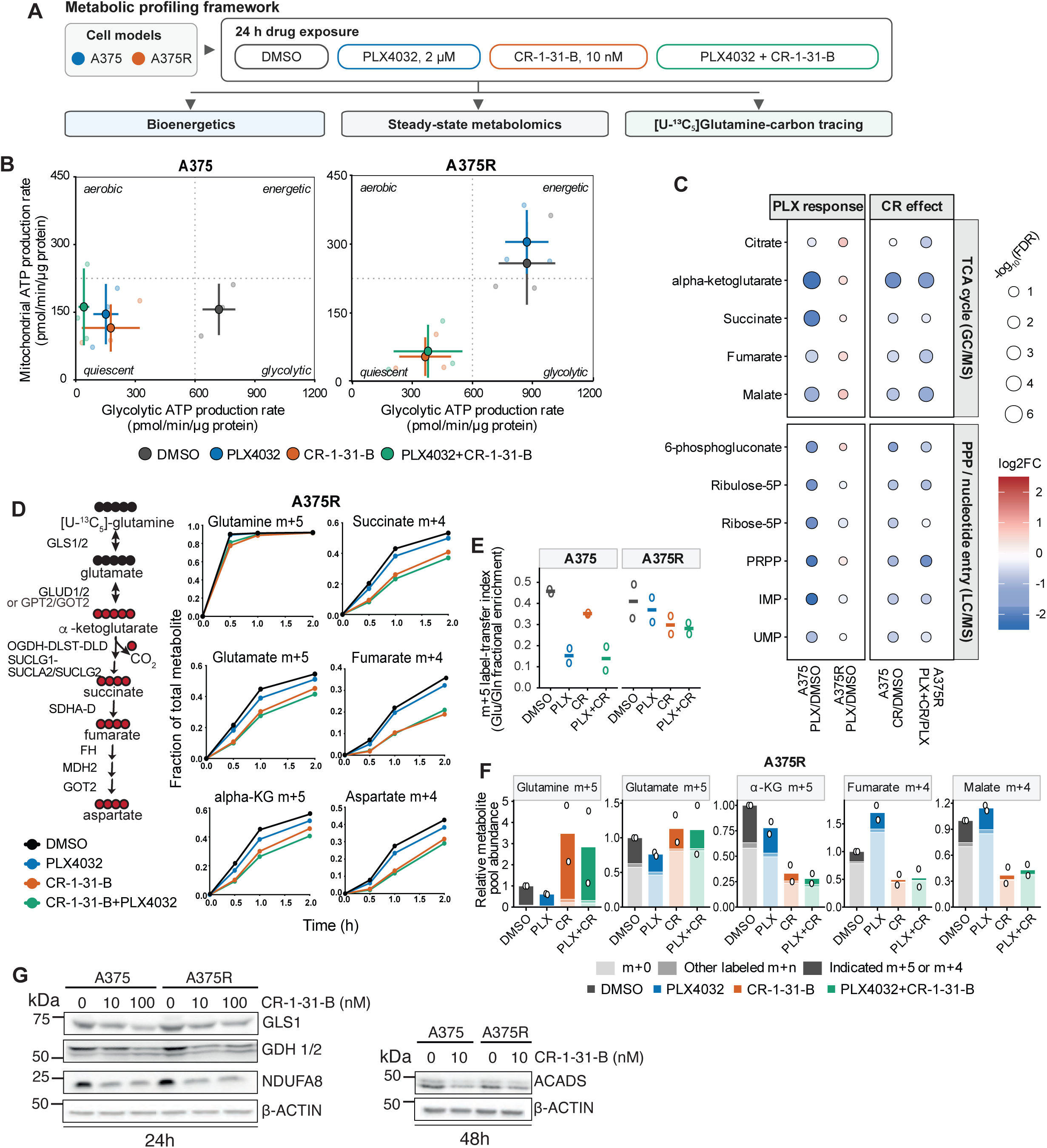
CR-1-31-B remodels bioenergetics and glutamine-carbon metabolism in BRAFi-sensitive and -resistant melanoma cells. **(A)** Seahorse XFe96 and [U-¹³C₅]glutamine tracing included DMSO, PLX4032 (2 µM), CR-1-31-B (10 nM), and PLX4032 plus CR-1-31-B in both A375 and A375R cells. Steady-state GC-MS/LC-MS included DMSO, PLX4032, and CR-1-31-B in A375 cells and DMSO, PLX4032, and PLX4032 plus CR-1-31-B in A375R cells. Seahorse and steady-state metabolomics used 24-h drug exposures. For isotope tracing, cells were pretreated for 24 h, and drugs remained present during the subsequent 2-h equilibration and tracing period. **(B)** Energetic maps showing mitochondrial and glycolytic ATP-production rates after the indicated treatments. Small translucent symbols represent three independent biological experiments; large symbols show means and horizontal and vertical bars show SD. Quadrant labels are descriptive guides. **(C)** Selected steady-state TCA-cycle and pentose-phosphate/nucleotide-entry metabolites. For the PLX4032 response, comparisons are A375 PLX4032/DMSO and A375R PLX4032/DMSO; for the CR-1-31-B effect, comparisons are A375 CR-1-31-B/DMSO and A375R PLX4032+CR-1-31-B/PLX4032. Color denotes mean log₂ fold change and dot size denotes −log₁₀ FDR, capped at 6 for display. GC-MS and LC-MS analyses each included three independent biological experiments per condition; statistical testing and FDR correction were performed separately within each platform and contrast, as described in Methods. **(D)** [U-¹³C₅]glutamine tracing through oxidative TCA metabolism in A375R cells. The schematic shows the expected isotopologues, and plots show fractional enrichment of glutamine m+5, glutamate m+5, α-ketoglutarate m+5, succinate m+4, fumarate m+4, and aspartate m+4 over 0.5-2 h. Lines show means from two independent biological experiments. **(E)** Glutamine-to-glutamate m+5 transfer index at 1 h, calculated as glutamate m+5 fractional enrichment divided by glutamine m+5 fractional enrichment. Symbols represent two independent biological experiments and horizontal bars show means. **(F)** Relative metabolite-pool abundance and isotopologue composition after 1 h tracing in A375R cells. Total bar height represents mean metabolite abundance after internal-standard and cell-number normalization, expressed relative to DMSO. Shading partitions m+0, other labeled isotopologues, and the indicated m+5 or m+4 isotopologue; open symbols represent two independent biological experiments. **(G)** Representative immunoblots of GLS1, GDH1/2, and NDUFA8 in A375 and A375R cells treated for 24 h with 0, 10, or 100 nM CR-1-31-B, and of ACADS after 48 h treatment with 0 or 10 nM CR-1-31-B. β-actin served as a loading control. Immunoblots are representative of three independent experiments.

Steady-state metabolomics showed that CR-1-31-B remodeled central carbon-metabolic pools in both BRAFi-sensitive and -resistant cells. In A375 cells, CR-1-31-B decreased TCA-cycle and pentose-phosphate/nucleotide-entry metabolites, including α-ketoglutarate (α-KG), succinate, fumarate, malate, ribulose-5-phosphate, ribose-5-phosphate, PRPP and IMP; in PLX4032-treated A375R cells, CR-1-31-B decreased the steady-state α-KG, fumarate and malate pools (Fig. 5C). In contrast, PLX4032 alone produced substantially weaker metabolic changes in A375R than in A375 cells, consistent with their differential BRAFi responsiveness. Across the broader GC-MS and LC-MS datasets, the CR-1-31-B-containing responses were strongly correlated between A375 and A375R cells (Spearman ρ = 0.83 for both platforms), with 94% and 85% directional concordance, respectively (Fig. S5D). Thus, CR-1-31-B imposes a broadly shared metabolic response despite the different sensitivities of the two cell states to BRAF inhibition.

The reduction in α-KG, a key entry point for glutamine-derived carbon into the TCA cycle, prompted us to examine whether CR-1-31-B restricts glutamine-carbon flow into downstream oxidative TCA metabolism. In A375R cells exposed to [U-¹³C₅]glutamine for 0.5, 1, 2 or 6 h, intracellular glutamine m+5 labeling was rapidly established and remained broadly similar among treatment conditions during the early 0.5–2 h interval (Fig. 5D), and this pattern persisted through 6 h (Fig. S5E). However, CR-1-31-B and PLX4032-plus-CR-1-31-B reduced the appearance of glutamate m+5 and α-KG m+5, followed by lower labeling of oxidative TCA products, including succinate, fumarate and aspartate m+4 (Fig. 5D). The glutamine-to-glutamate m+5 transfer index was likewise lower under CR-1-31-B-containing conditions (Fig. 5E). At 1 h, CR-1-31-B-treated cells retained or accumulated glutamine and glutamate pools while α-KG and downstream TCA pools were depleted (Fig. 5F). Extended tracing in both A375 and A375R cells supported reduced transfer of glutamine-derived carbon into oxidative TCA intermediates rather than loss of intracellular glutamine labeling itself (Fig. S5E-I). Moreover, cell-number-normalized glutamine consumption was not significantly reduced by CR-1-31-B in either cell state (Fig. S5J). Together, these findings indicate that eIF4A inhibition constrains intracellular glutamine-carbon processing downstream of uptake.

Targeted immunoblotting showed that CR-1-31-B reduced the abundance of GLS1 and GDH1/2, which convert glutamine to glutamate and glutamate to α-ketoglutarate, respectively, as well as the complex-I subunit NDUFA8 at 24 h; ACADS, a mitochondrial short-chain fatty-acid-oxidation enzyme, was reduced at 48 h in A375 and/or A375R cells (Fig. 5G). GLS2 and ATP6V1F were also reduced at 24 h (Fig. S5K). To assess whether this remodeling extended beyond the proteins examined by immunoblotting, we used the 36-h proteomic dataset to survey a pathway-defined group of 18 enzymes and transporters involved in glutamine utilization, TCA-cycle reactions and associated carbon-shuttle pathways. Proteins were included for biochemical relevance independently of response direction or statistical significance, and their effects are displayed alongside the 24-h steady-state metabolite changes for descriptive pathway context (Fig. S5L; Supplementary Table S5). Significant decreases affected glutamate–α-KG routing (GPT2), mitochondrial α-ketoglutarate–malate exchange (SLC25A11), α-ketoglutarate-dehydrogenase and succinate-dehydrogenase/complex-II functions (DLD, SDHA and SDHB), and isocitrate/α-ketoglutarate metabolism (IDH1 and IDH2). Conversely, proteins involved in other TCA-cycle and malate–aspartate-shuttle reactions increased (SUCLG2, FH, MDH2 and GOT2). Other quantified proteins did not meet the predefined significance criteria; GLUD2 was not detected. Thus, CR-1-31-B induced selective, node-specific protein remodeling rather than uniform suppression of the glutamine–TCA pathway. RT-qPCR showed that GLS1, NDUFA8 and ACADS transcripts were not lower at 48 h, and GDH1/2 transcripts were not lower at 72 h, arguing against sustained concordant mRNA loss at the assayed later time points (Fig. S5M,N). Consistent with the functional importance of this axis, reducing extracellular glutamine depleted glutamine-linked metabolite pools and reduced the number of viable cells accumulated over 72 h in both A375 and A375R cultures (Fig. S5O,P). The difference in viable-cell number between vehicle- and low-dose-CR-1-31-B-treated cultures narrowed as glutamine availability declined, suggesting that glutamine limitation and eIF4A inhibition constrain an overlapping component of cell fitness (Fig. S5P).

Collectively, these data show that eIF4A inhibition induces a common low-output metabolic state in BRAFi-sensitive and -resistant melanoma cells by constraining bioenergetic capacity and glutamine-carbon transfer into the TCA cycle.

### Combined BRAF and eIF4A inhibition prolongs tumor control in A375 xenografts

Based on the activity of CR-1-31-B in BRAFi-resistant cells and its suppression of resistance-associated molecular and metabolic programs (Figs. 1, 3–5), we next tested whether combined eIF4A and BRAF inhibition could improve the depth and durability of tumor control *in vivo*. A375 cells were implanted subcutaneously into nude mice, and established tumors were randomized to vehicle, PLX4720, CR-1-31-B, or PLX4720 plus CR-1-31-B (Fig. 6A). Vehicle-treated tumors grew rapidly, while CR-1-31-B monotherapy delayed tumor growth relative to vehicle during the evaluable comparison window (treatment-by-time interaction, P = 0.0011). PLX4720 initially suppressed tumor growth but was followed by progressive regrowth in multiple animals. The combination produced deeper and more sustained control, with substantially lower tumor volumes than PLX4720 alone over the prespecified evaluable window through day 49 (treatment-by-time interaction, P = 6.59 × 10⁻⁵; Fig. 6B). Body-weight monitoring showed an early decrease in the combination group followed by recovery during continued treatment, without progressive weight loss (Fig. S6A).

**Fig. 6.**
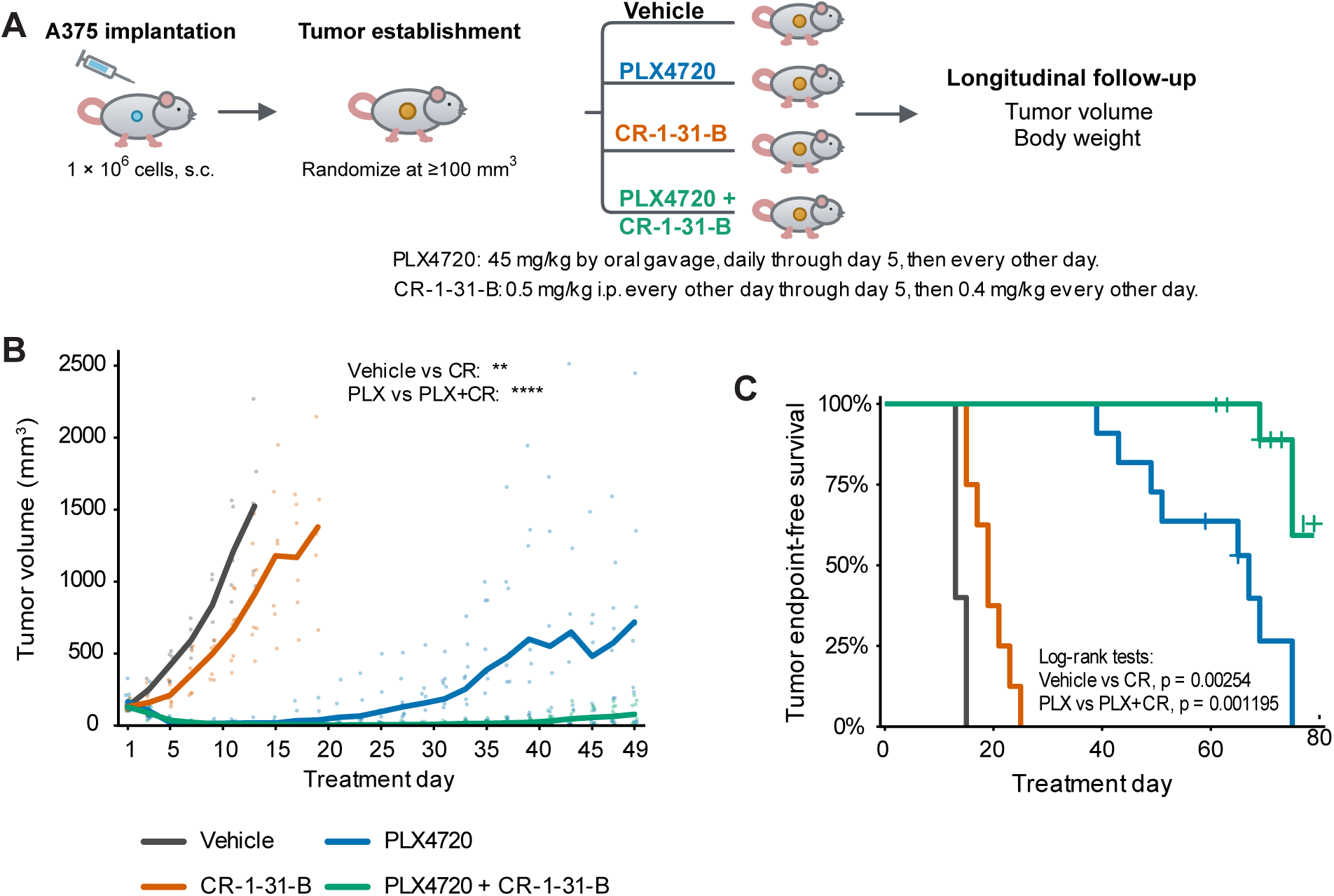
Combined BRAF and eIF4A inhibition prolongs tumor control in A375 xenografts. **(A)** Experimental design. Nude mice were implanted subcutaneously with 1 × 10⁶ A375 cells and randomized when tumors reached approximately 100–150 mm³. Mice were treated with vehicle, PLX4720, CR-1-31-B, or PLX4720 plus CR-1-31-B. PLX4720 was administered at 45 mg/kg by oral gavage daily through treatment day 5 and every other day thereafter. CR-1-31-B was administered intraperitoneally at 0.5 mg/kg every other day through day 5 and at 0.4 mg/kg every other day thereafter. **(B)** Tumor volumes through treatment day 49. Small symbols show individual measurements, and lines show observed group means. Group means were displayed only while at least five mice contributed. Longitudinal comparisons used comparison-specific evaluable windows: through day 13 for vehicle versus CR-1-31-B, the last time point at which all five vehicle-treated mice remained under observation, and through day 49 for PLX4720 versus the combination; at day 49, nine PLX4720-treated and 11 combination-treated mice remained under observation. Stars denote treatment-by-time interactions from longitudinal mixed-effects models comparing vehicle with CR-1-31-B through day 13 (P = 0.0011) and PLX4720 with PLX4720+CR-1-31-B through day 49 (P = 6.59 × 10⁻⁵). **(C)** Tumor endpoint-free survival in the same cohorts. An event was defined as attainment of a humane endpoint; mice remaining below endpoint at administrative study termination were retained as right-censored observations. Pairwise log-rank tests compared vehicle with CR-1-31-B (P = 0.00254) and PLX4720 with PLX4720+CR-1-31-B (P = 0.001195). Cohorts comprised vehicle, n = 5; PLX4720, n = 11; CR-1-31-B, n = 8; and PLX4720 plus CR-1-31-B, n = 11.

The individual trajectories showed that PLX4720 responses were heterogeneous and frequently followed by regrowth, whereas combination-treated tumors generally remained controlled for longer, although a subset eventually progressed (Fig. S6B). Consistent with these growth patterns, CR-1-31-B prolonged tumor endpoint-free survival relative to vehicle (P = 0.00254), and PLX4720 plus CR-1-31-B prolonged endpoint-free survival relative to PLX4720 alone (P = 0.001195; Fig. 6C).

Together, these results show that combined BRAF and eIF4A inhibition provides more durable control of A375 xenografts than BRAF inhibition alone and identifies eIF4A as a preclinical therapeutic vulnerability in this model.

## Discussion

This study identifies eIF4A-dependent mRNA translation as a vulnerability in melanoma models with acquired resistance to BRAF inhibition and links it to the molecular and metabolic organization of the resistant state. Across BRAFi-sensitive and -resistant models, multiple rocaglate eIF4A inhibitors reduced cell accumulation at low-nanomolar concentrations. CR-1-31-B rapidly reduced nascent protein synthesis, lowered selected cell-cycle and survival proteins, induced apoptosis, and impaired clonogenic recovery. In the A375/A375R model, resistance involved coordinated changes in RNA abundance, selective mRNA translation, protein output, cell-state and extracellular-matrix programs, survival signaling, and mitochondrial metabolism. CR-1-31-B acted across these layers, preferentially lowering the levels of resistance-acquired proteins and imposing a similar low-output metabolic state in A375 and A375R despite their different responses to PLX4032. It was also associated with reduced downstream labeling of glutamine-derived carbon and, in A375 xenografts, deepened and prolonged tumor control when combined with BRAF inhibition. Collectively, the data support a model in which eIF4A-dependent translation sustains several outputs of resistant-cell fitness rather than acting as a single, linear resistance mechanism (Fig. 7).

**Fig. 7.**
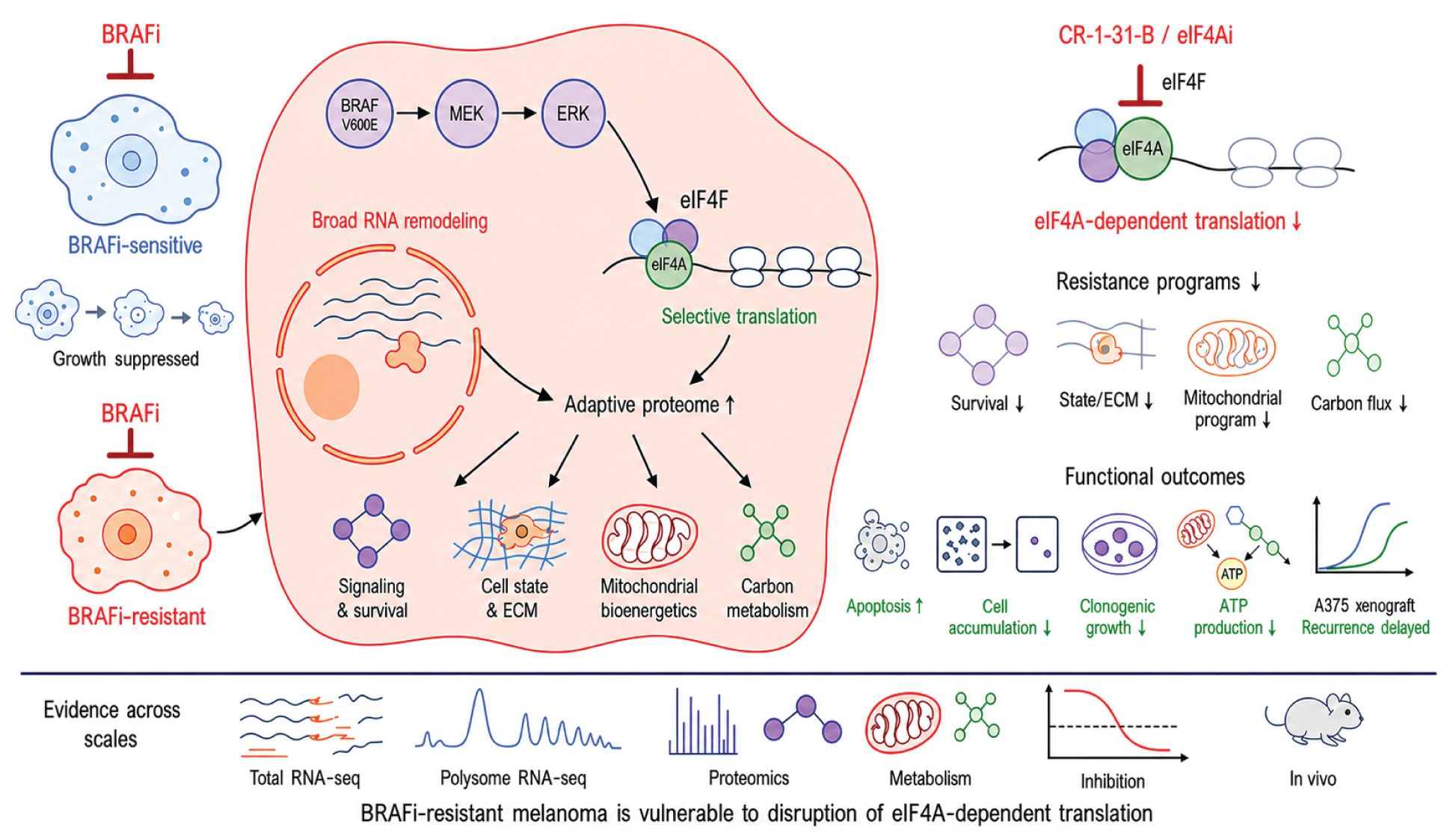
Model of eIF4A-dependent adaptation in BRAFi-resistant melanoma. BRAF–MEK–ERK signaling converges on eIF4F. In BRAFi-resistant cells, broad RNA remodeling and selective translation shape an adaptive proteome supporting survival, cell-state/ECM programs, mitochondrial bioenergetics, and carbon metabolism. Pharmacologic eIF4A inhibition suppresses these programs, increases apoptosis, reduces cell accumulation, clonogenic growth, and ATP production, and delays A375 xenograft regrowth. The model summarizes integrated transcriptomic, translatomic, proteomic, metabolic, pharmacologic, and in *vivo* evidence from this study. The illustration was generated using OpenAI’s image-generation tool within Codex under author direction; all scientific content and labels were reviewed and verified by the authors.

Established BRAFi-resistant melanoma cells remained sensitive to eIF4A inhibition. Comparable activity of CR-1-31-B, SDS-1-021, silvestrol, and CMLD012612 indicates that this response is not unique to CR-1-31-B, although individual compounds can differ in RNA-clamping and translational selectivity^40^ and in pharmacokinetic and *in vivo* properties^42^. Activity in A375R and LU1205R cells, and in MDA-MB-231R cells selected for resistance to combined BRAF/MEK inhibition, shows retained cellular activity after resistance to MAPK-directed therapy. Braf^V600E^-driven YUMMER1.7 and parental MDA-MB-231 cells extend the response across distinct backgrounds. This is consistent with work identifying eIF4F as a convergence point for BRAF/MEK-inhibitor resistance and eIF4A-dependent translation as a requirement for melanoma persister-cell survival^14,18^. Recent studies further show that eIF4F-dependent DUSP6 synthesis shapes ERK-feedback control and eIF4A-dependent 53BP1 synthesis can promote adaptive mutability during resistance acquisition^37,50^. In addition, eIF4F inhibition can activate AMPK through an LKB1-independent mechanism involving reduced protein phosphatase 2A abundance, directly connecting translation initiation to metabolic signaling in melanoma^58^. Our findings extend this framework to established resistance and identify CR-1-31-B-sensitive outputs spanning extracellular-matrix and adhesion programs, cell-state plasticity, survival, mitochondrial bioenergetics, and glutamine-carbon metabolism. Thus, eIF4A-dependent translation represents a vulnerability shared by several programs that sustain resistant-cell fitness.

The A375R/A375 comparison showed that established resistance was characterized largely by RNA-abundance remodeling, with selective changes in translation efficiency and translational buffering superimposed. Polysome-associated RNA tracked protein changes more closely than translation efficiency alone, consistent with protein output integrating transcript availability and ribosome engagement. This multilayered remodeling encompassed neural crest-like or dedifferentiated states, extracellular-matrix and adhesion programs, anti-apoptotic signaling, and mitochondrial functions. NGFR, AXL, SPARC, and BCL2L1 exemplify these programs but are not proposed as a universal resistance marker set. This pattern accords with the state plasticity, matrix remodeling, and survival rewiring reported in drug-tolerant and resistant melanoma populations^5,6,17,22^.

Direct alignment of the resistance and CR-1-31-B-response proteomes showed that eIF4A inhibition preferentially opposes proteins gained during resistance. This asymmetric response was concentrated in extracellular-matrix, adhesion, state-plasticity, survival, and metabolic programs. It was nevertheless partial: eIF4A inhibition attenuated a coordinated resistance-acquired module, including AXL- and matrix-associated outputs, without reverting cells to the parental state or reversing every upstream alteration.

A cross-dataset comparison with two public longitudinal melanoma PDX trajectories suggested concordance with treatment-associated states beyond A375. Programs induced during early response, persistence/MRD, or resistant regrowth—and those sustained from persistence through regrowth—were modestly shifted toward lower corresponding protein abundance in the A375R CR-1-31-B response. Convergence was strongest during persistence, whereas the regrowth-transition program was not significantly shifted. The overlapping genes spanned signaling and stress survival, melanoma state and adhesion, metabolism, and cell-cycle/regrowth functions (Fig. S4E). These associations remain hypothesis-generating and require direct testing of eIF4A inhibition in additional patient-derived models.

CR-1-31-B produced distinct responses across regulatory levels. At 1 h, altered polysome association occurred with limited RNA-abundance change, whereas the 24-h RNA and 36-h proteomic profiles captured broader remodeling. Because these layers were generated in independent experiments at single intervals, they do not establish a strict temporal sequence. Their discordance nevertheless indicates that protein changes can be associated with selective translation, later RNA-abundance remodeling, or effects detected primarily at the protein level. MYC/translation-related protein enrichment and stress/TNF–NF-κB RNA responses may represent compensatory activity, although their functional role remains untested. This possibility is consistent with reported ERK hyperactivation following loss of eIF4F-dependent DUSP6 and NRF2-mediated resistance to rocaglates^37,48^.

The 5′UTR analysis provides a candidate link between transcript selectivity and rocaglate-induced eIF4A clamping. CR-1-31-B-sensitive transcripts were enriched for purine-rich, locally structured leaders, consistent with rocaglate stabilization of eIF4A on polypurine RNA^40–43^. Predicted G4 features contributed but were not consistently supported across stringent definitions, suggesting that G4 potential is one element of a broader sequence and structural context. These associations do not identify direct targets, particularly because public rather than A375R-specific transcript annotations were used^30,32^. Mapping transcription start sites and testing selected leaders by reporter mutagenesis will be needed to establish isoform-specific causality.

Metabolic remodeling was prominent in the resistant state. A375R cells were enriched for respiratory-complex-I and mitochondrial-ribosome proteins, produced more mitochondrial ATP, and showed greater glucose/pyruvate oxidation capacity, consistent with the oxidative phenotype reported in a subset of MAPKi-resistant melanomas^23–25,59,60^. Fatty-acid and glutamine oxidation capacities were lower, indicating redistribution toward glucose-supported respiration rather than uniformly increased fuel flexibility. Thus, A375R represents one oxidative resistance state rather than a universal metabolic template. CR-1-31-B produced a similar lower-output metabolic response in A375 and A375R cells despite their different responses to PLX4032, reducing glycolytic and mitochondrial ATP production and selected TCA-cycle and pentose-phosphate/nucleotide pools. Coordinated changes in mitochondrial transport, complex-I assembly, carbon routing, redox, and biosynthetic proteins support a broad metabolic response consistent with established links between eIF4A/eIF4F-dependent translation and bioenergetic control^32,38,45,61^. Reduced biosynthetic demand following growth inhibition may contribute, but the cell-number-normalized bioenergetic measurements and accompanying protein and isotope-tracing changes argue against cell number alone explaining the effect.

Glutamine tracing pointed to reduced carbon processing downstream of uptake: extracellular consumption was not consistently reduced and intracellular glutamine m+5 labeling was preserved, whereas labeling of glutamate, α-KG, and downstream oxidative TCA products decreased. Changes in GLS1, GDH1/2, NDUFA8, and other nodes may contribute, but no single bottleneck was identified. Lower acute glutamine-oxidation capacity and robust isotope labeling are not contradictory because the assays measure potential capacity and realized carbon transfer, respectively. Reduced cell accumulation under glutamine limitation supports the importance of this axis, although the smaller difference between vehicle and low-dose CR-1-31-B under these conditions does not establish direct inhibition of glutaminolysis. This interpretation is consistent with reports that BRAFi-resistant or high-OXPHOS melanomas can rely on glutamine for TCA-cycle and redox metabolism^26,60,62^. Metabolite-rescue and targeted protein-restoration experiments will be needed to test causality.

In BRAFi-sensitive A375 xenografts, PLX4720 initially controlled tumors but was followed by heterogeneous regrowth; adding CR-1-31-B deepened and prolonged tumor control and extended endpoint-free survival, consistent with evidence that targeting eIF4F/eIF4A can enhance MAPK-directed therapy^14,18,50^. The *in vitro* data suggest several tumor-intrinsic programs that may contribute, but treated tumors were not molecularly profiled and the mechanism of benefit remains unresolved. Regrowth in some combination-treated tumors indicates delayed rather than uniform prevention of escape. Testing eIF4A inhibition with a combined BRAF/MEK backbone in resistant and patient-derived models will determine the generality of this effect.

## Limitations of the study

Several limitations constrain interpretation. The integrated omics centered on one chronically selected A375/A375R pair; eIF4A-inhibitor activity was confirmed in additional models, but they were not profiled at equivalent depth, and A375R represents only one resistance trajectory. Molecular layers were generated in independent experiments at single intervals, supporting cross-layer comparison but not a formal temporal sequence or direct causality. The isotope-tracing dataset was descriptive (n = 2). The public PDX analysis used whole-section pseudobulk profiles from one tumor per model and stage and therefore cannot distinguish tumor-intrinsic regulation from lineage/state composition, spatial sampling, or tumor purity. Public 5′UTR annotations may not reflect A375R-specific isoforms. Finally, the xenograft study tested upfront treatment of BRAFi-sensitive A375 tumors rather than established resistant disease; the use of athymic mice limited assessment of immune contributions, and treated tumors were not molecularly profiled.

## Conclusion

In summary, acquired BRAFi resistance in A375R cells involved coordinated changes in RNA abundance, selective translation, and protein abundance across signaling, survival, cell-state/ECM, and metabolic programs. CR-1-31-B was active in BRAFi-resistant models, preferentially suppressed resistance-acquired proteins, reduced bioenergetic output, and limited glutamine-carbon transfer downstream of uptake. Combining eIF4A and BRAF inhibition improved control of A375 xenografts, supporting evaluation of eIF4A inhibitors alongside combined BRAF/MEK therapy in patient-derived models. Together, these findings identify eIF4A-dependent translation as a vulnerability shared by multiple programs supporting resistant melanoma-cell fitness.

## Materials and Methods

### Cell lines

A375 and A375R human melanoma cells were kindly provided by Dr. Ze’ev Ronai^51^. The acquired BRAF-inhibitor-resistant derivative A375R was generated from A375 cells by chronic stepwise selection with PLX4032. A375R stock cultures were maintained in complete medium containing PLX4032 (10 µM) and were cultured without the selecting drug for one passage before each experiment.

LU1205R melanoma cells were kindly provided by Dr. Ze’ev Ronai^51^. YUMMER1.7 murine melanoma cells, carrying Braf^V600E^ together with Pten and Cdkn2a loss, were kindly provided by Dr. Sonia del Rincón (McGill University); the model was originally developed in Dr. Marcus Bosenberg’s laboratory (Yale University)^63^.

MDA-MB-231 breast cancer cells carrying BRAF^G464V^ and the matched encorafenib/binimetinib-resistant derivative (MDA-MB-231R) were provided by April A. N. Rose (McGill University). MDA-MB-231R stocks were maintained in complete medium containing encorafenib (1,000 nM) and binimetinib (200 nM), and cultured without these inhibitors for one passage before experiments.

A375, A375R, LU1205R, MDA-MB-231, and MDA-MB-231R cells were cultured in Dulbecco’s modified Eagle medium (DMEM; Wisent BioProducts, 319-005-CL) supplemented with 10% heat-inactivated HyClone fetal bovine serum (Cytiva, SH30396.03) and 1% penicillin-streptomycin (Wisent BioProducts, 450-201-EL). YUMMER1.7 cells were cultured in DMEM/F12 supplemented with 10% fetal bovine serum, 1% nonessential amino acids, and 1% penicillin-streptomycin. Cells were maintained at 37°C in a humidified atmosphere containing 5% CO₂ and used within 20 passages after thawing.

Mycoplasma contamination was monitored using a PCR-based detection kit (Applied Biological Materials, G238) according to the manufacturer’s instructions. Human cell-line identity was assessed by short-tandem-repeat profiling through the Génome Québec Cell Line Authentication Service. The LU1205R STR profile used in this study was consistent with that of the Wistar-derived culture.

### Compounds and treatment preparation

SDS-1-021, silvestrol, and CMLD012612 were generous gifts from Dr. Jerry Pelletier, McGill University. CR-1-31-B (MedChemExpress, HY-136453), PLX4032 (vemurafenib; Selleck Chemicals, S1267), encorafenib (ChemieTek, CT-LGX818), and binimetinib (ChemieTek, CT-A162) were obtained from the indicated suppliers. Puromycin was from Sigma-Aldrich (5088380001). The final DMSO concentration did not exceed 0.15% (v/v) and was matched between treatment and vehicle-control conditions. Treatment concentrations and durations are stated below and in the corresponding figure legends.

### Viable-cell counting and concentration-response analyses

For A375 and A375R endpoint viable-cell counts, 3 × 10⁴ cells were seeded per well in six-well plates (Sarstedt, 83.3902) in 3 mL complete medium. After 24 h, cells were exposed to the indicated compounds for 72 h. Cells were collected by trypsinization (Wisent BioProducts, 325-043-EL), mixed with trypan blue (Corning, 25-900-CI), and counted using a DeNovix CellDrop automated cell counter. Viable-cell numbers were expressed relative to the matched vehicle control. The A375 and A375R PLX4032 curves and CR-1-31-B curves were derived from independent experiments as specified in the figure legends.

For LU1205R concentration-response assays, 1.5 × 10⁵ cells were seeded per well in six-well plates (Sarstedt, 83.3902) containing 3 mL complete medium. After 24 h, cells were exposed to the indicated compounds for 72 h and viable cells were quantified by trypan-blue exclusion using a DeNovix CellDrop automated cell counter. For the MDA-MB-231/MDA-MB-231R CR-1-31-B treatment, 1 × 10⁴ MDA-MB-231 cells or 2 × 10⁴ MDA-MB-231R cells were seeded per well in six-well plates containing 3 mL complete medium. After 24 h, cells were treated for 72 h and viable cells were quantified by trypan-blue exclusion using the CellDrop counter.

YUMMER1.7 cells were seeded at a density of 1 × 10⁴ cells per well in 12-well plates (Corning Costar, Cat. No. 3513) in 1 mL of culture medium and allowed to adhere for 24 h prior to treatment. Cells were then treated with CR-1-31-B at final concentrations of 0.1, 0.5, 1, 2.5, 5, 10, or 100 nM, with DMSO-treated cells serving as the vehicle control (0 nM). Each treatment condition was performed in triplicate. Cells were treated for 72 h, and cell accumulation was quantified using Incucyte segmentation-based cell counting. Three independent biological experiments were performed.

Biological experiments were analyzed independently; technical wells or counts were averaged before biological-level summaries.

Concentration–response curves were fitted in R (version 4.5.1; R Foundation for Statistical Computing) by nonlinear least squares using stats::nls. Curves were plotted using ggplot2 (version 4.0.2). Four-parameter logistic models were used when both asymptotes could be estimated; for normalized datasets in which the upper response was constrained by vehicle normalization, a variable-slope logistic model with the upper asymptote fixed at 100% was used. IC_50_ values were calculated from the fitted log_10_ IC_50_ parameter. Vehicle observations were displayed but were not log-transformed. For A375 treated with PLX4032, three independently fitted experiments were summarized as the arithmetic mean ± SD; A375R PLX4032 treatment was reported as IC_50_ >10 µM because 50% inhibition was not reached within the tested range.

### Apoptosis analysis

A375 and A375R cells were seeded at 3 × 10⁴ cells per well in six-well plates and treated for 72 h. Adherent and nonadherent cells were collected, washed, and stained for 15 min in the dark with APC-conjugated Annexin V (1:400; BioLegend, 640941) and DAPI (1:32,000) in binding buffer. Samples were acquired on a BD FACSCelesta flow cytometer, with 50,000 events collected per sample, and analyzed in FlowJo v10.7.1. After exclusion of debris and doublets, viable cells were defined as Annexin V−/DAPI−. The apoptotic fraction was calculated as the sum of Annexin V+/DAPI− and Annexin V+/DAPI+ populations. Three independent biological experiments were performed, with three technical replicates each.

### Clonogenic assays

For post-treatment clonogenic recovery assays, A375 and A375R cells were exposed to the indicated compounds or combinations for 72 h, collected, counted, and replated at 1,000 viable cells per well in six-well plates containing 3 mL drug-free complete medium. Medium was replaced every 2 days, and colonies were allowed to form for 6 days after replating. Colonies were fixed and stained with a solution containing 50% ethanol, 2.5 mg/mL NaCl, 1.78% formaldehyde (Fisher Scientific, F79500), and 7.5 mg/mL crystal violet (Sigma-Aldrich, 6158), and scanned under identical settings. Colony-associated crystal-violet intensity was quantified in ImageJ v1.54b and normalized to the matched vehicle control. Three independent biological experiments were analyzed; technical wells were averaged within experiment.

MDA-MB-231 and MDA-MB-231R cells were seeded at 1,000 viable cells per well in six-well plates containing 3 mL complete medium. After 24 h, cells were exposed to vehicle, CR-1-31-B (2.5, 5, or 10 nM), or encorafenib/binimetinib combinations (0.125/0.05, 0.5/0.1, 2/0.4, 4/0.8, 6/1.2, 8/1.6, or 16/3.2 µM, respectively). Drug-containing medium was replaced every 72 h, and colonies were allowed to form for 6 days. Colonies were then fixed, stained, scanned, and quantified as described above. Three independent biological experiments were performed, and technical wells were averaged within each experiment.

### Puromycin-incorporation assay

To assess nascent protein synthesis, 1 × 10⁶ cells were seeded in 100-mm dishes containing 10 mL complete medium. After 24 h, cells were treated with vehicle, CR-1-31-B (10 nM), PLX4032 (2 µM), or their combination for 1 or 24 h, as indicated. Puromycin (10 µg/mL; Sigma-Aldrich, 5088380001) was added for the final 15 min. Cells were lysed as described below, and incorporated puromycin was detected by immunoblotting. Total puromycin signal across each lane was quantified in ImageJ, normalized to β-actin, and expressed relative to the corresponding vehicle control. The displayed experiments represent three independent biological experiments.

### Immunoblotting

Cells were lysed on ice in buffer containing 50 mM Tris-HCl (pH 7.5), 150 mM NaCl, 0.1% SDS, 1% NP-40, 0.5% sodium deoxycholate, 1× cOmplete EDTA-free protease-inhibitor cocktail (Roche, 05056489001), and 1× PhosSTOP phosphatase-inhibitor cocktail (Roche, 04906837001). Protein concentrations were determined with the Pierce BCA Protein Assay Kit (Thermo Fisher Scientific, 23225) using a TECAN Infinite M1000 PRO plate reader.

Equal amounts of protein were separated by SDS-PAGE and transferred to nitrocellulose membranes (Cytiva, 45004002). Membranes were blocked in 5% nonfat dry milk or 5% bovine serum albumin in TBST and incubated with primary antibodies overnight at 4°C. Horseradish-peroxidase-conjugated anti-rabbit IgG (1:4,000; Cell Signaling Technology, 7074) or anti-mouse IgG (1:4,000; Cell Signaling Technology, 7076) was applied for 1 h at room temperature. Signal was developed using ECL reagent (PerkinElmer, 0RT2755) or SuperSignal West Femto substrate (Thermo Fisher Scientific, 34096) and acquired with an Azure 600 imager.

Primary antibodies used included mouse monoclonal β-actin (1:5,000; Sigma-Aldrich, A5441), anti-puromycin clone 12D10 (1:1,000; Millipore, MABE343), rabbit monoclonal BCL-2 clone D55G8 (1:1,000; Cell Signaling Technology, 4223), mouse monoclonal cyclin D3 (1:1,000; Cell Signaling Technology, 2936), rabbit polyclonal CDK4 (1:1,000 in 2% skim milk; Abcam, ab137675), rabbit monoclonal AXL clone C89E7 (1:1,000; Cell Signaling Technology, 8661), mouse monoclonal total RPS6 clone C-8 (1:2,000; Santa Cruz Biotechnology, sc-74459), rabbit polyclonal phospho-RPS6 Ser240/244 (1:1,000; Cell Signaling Technology, 2215S), rabbit polyclonal total ERK1/2 (1:1,000; Cell Signaling Technology, 9102S), and rabbit polyclonal phospho-ERK1/2 Thr202/Tyr204 (1:1,000; Cell Signaling Technology, 9101S).

Metabolic targets were detected using rabbit monoclonal GLS1 clone E9H6H (1:1,000; Cell Signaling Technology, 56750S), rabbit polyclonal GLS2 (1:1,000; Abcam, ab113509), rabbit monoclonal GDH1/2 clone D9F7P (1:500; Cell Signaling Technology, 12793), rabbit monoclonal NDUFA8 clone EPR15303 (1:1,000; Abcam, ab184952), rabbit monoclonal ACADS/SCAD clone EPR10861(B) (1:1,000; Abcam, ab154823), and rabbit monoclonal ATP6V1F clone EPR15053 (1:2,000; Abcam, ab190789). The complete antibody inventory is provided in Supplementary Table S6.

Immunoblots are representative of at least three independent experiments unless stated otherwise. Where quantified, band intensity was measured from unsaturated images in ImageJ, normalized to β-actin, and then normalized to the matched vehicle control.

### RNA extraction, reverse transcription, and quantitative PCR

Total RNA was isolated using TRIzol Reagent (Invitrogen/Life Technologies, 15596018) according to the manufacturer’s instructions. RNA concentration and purity were assessed using a NanoDrop spectrophotometer, and cDNA was synthesized from 1 µg total RNA using the All-in-One 5X RT MasterMix (abm, G592). Quantitative PCR was performed using BlasTaq 2X qPCR MasterMix (abm, G892) on a QuantStudio 12K Flex Real-Time PCR System. Relative transcript abundance was calculated using the 2⁻ΔΔCt method. Primer sequences and amplification efficiencies are provided in Supplementary Table S6. For validation of GLS1, NDUFA8, and ACADS, A375 and A375R cells were treated with vehicle or CR-1-31-B (10 nM) for 48 h. GLS1 was analyzed in four independent biological experiments, whereas NDUFA8 and ACADS were analyzed in three. Expression was normalized separately to ACTB and TUBB, and the two normalized values were combined as their geometric mean within each experiment. Values were expressed as log₂ fold change relative to the matched DMSO control. In a separate 72-h experiment, GDH1/2 mRNA abundance was measured following treatment with CR-1-31-B (7 nM), normalized to ACTB, and expressed relative to DMSO. Three independent biological experiments were performed. The primer assay does not distinguish GLUD1 from GLUD2.

### Polysome Profiling and RNA Sequencing

The polysome/RNA-sequencing data reported in this study were generated in an experiment comprising six independent biological replicates per condition across two experimental batches, with two predefined comparisons: DMSO-treated A375R versus DMSO-treated A375 cells, and A375R cells treated for 1 h with CR-1-31-B (10 nM) versus DMSO.

For each replicate, 4 × 10⁶ cells were seeded in 150-mm dishes (Corning, 430599) containing 20 mL complete medium and allowed to attach for 48 h, at which time treatment was added as indicated above. Medium was changed twice, 24 h and 47 h post-seeding. Cycloheximide (100 µg/mL; Sigma-Aldrich, 01810) was added for the final 5 min of treatment. Cells were washed twice with 10 mL ice-cold PBS containing cycloheximide (100 µg/mL) before lysis in ice-cold hypotonic buffer containing 5 mM Tris-HCl (pH 7.5), 2.5 mM MgCl₂, 1.5 mM KCl, 100 µg/mL cycloheximide, 2 mM dithiothreitol, 0.5% Triton X-100, and 0.5% sodium deoxycholate^64^.

Clarified lysates were layered onto 5–50% sucrose gradients and centrifuged at 150,000 × g for 2 h in a Sorvall WX+ ultracentrifuge (Thermo Fisher Scientific, 75000100) at 4°C, with acceleration and deceleration disabled. Gradients were generated and fractionated with a BioComp piston-gradient fractionator while absorbance at 260 nm was monitored with a BioComp Triax flow cell. Fractions starting with and including the peak corresponding to four ribosomes were pooled as the polysome-associated sample (the 80S monosome peak corresponds to one ribosome). An aliquot of the same clarified lysate loaded onto the gradient was retained as the paired input sample. RNA was purified from the input aliquot and pooled polysome fractions using the Direct-zol RNA MicroPrep kit (200-preparation format; Zymo Research, R2062).

#### Library preparation, sequencing, alignment, and count generation

RNA integrity was assessed by the IRIC Genomics Platform before library preparation. Stranded, ribosomal RNA-depleted libraries were sequenced as 84-bp single-end reads on an Illumina NextSeq 500/550 instrument. Reads were aligned to the human GRCh38 reference genome using STAR v2.7.9a and Ensembl release 110 annotations. Gene-level counts were generated using featureCounts v2.0.3.

#### Anota2Seq regulatory-mode analysis

Paired input and polysome-associated RNA counts were analyzed with Anota2Seq v1.30.0 in R v4.5.1, following the framework described by Oertlin et al.^55^. Low-abundance features were filtered independently in the input and polysome compartments using edgeR filterByExpr^65^, and only genes retained in both compartments were analyzed. TMM-normalized log_2_ expression values were modeled without pre-shifting, with experimental batch included in the design.

Anota2Seq distinguishes changes in RNA abundance, changes in translational efficiency (TE; polysome-association changes beyond those explained by input RNA), and translational buffering. Translation-mode calls required Benjamini-Hochberg-adjusted P < 0.05, APV slope from −1 to 2, |APV effect| ≥ log_2_(1.2), and a concordant |deltaPT| ≥ log_2_(1.2). Buffering-mode calls required adjusted P < 0.05, APV slope from −2 to 1, |APV effect| ≥ log_2_(1.2), and a concordant |deltaTP| ≥ log_2_(1.2). Genes were classified as mRNA-abundance regulated by anota2seqRegModes when concordant changes in total and polysome-associated RNA each met an absolute APV effect threshold of ≥log_2_(1.2) (0.263), an RVM-adjusted Benjamini–Hochberg FDR < 0.05, and the corresponding directional delta thresholds (selDeltaT and selDeltaP) of log_2_(1.2). The same harmonized Anota2Seq selection criteria were applied to the untreated A375R-versus-A375 and A375R CR-1-31-B-versus-DMSO comparisons.

For PCA visualization, input RNA and polysome-associated RNA counts were variance-stabilized separately using DESeq2^66^. Relative TE was represented as the difference between matched polysome-associated and input RNA variance-stabilized values. Batch effects were removed from these matrices using limma::removeBatchEffect, while preserving biological condition in the design matrix; this adjustment was used only for visualization, whereas batch was included in the differential models. During quality control, concordant clustering of both input and polysome-associated RNA profiles identified a reciprocal labeling error affecting A375 and A375R libraries in Batch A. Sample labels were reassigned before downstream analyses; the original sequencing-library identifiers were retained in the GEO metadata to preserve traceability to the raw FASTQ files. As an alternative-method sensitivity analysis of relative TE, raw input and polysome-associated RNA counts were analyzed using DESeq2 v1.48.2 with the design ∼ batch + condition + fraction + condition:fraction. The condition-by-fraction interaction tested whether the polysome-versus-input change differed between conditions.

### 24-h total RNA sequencing

A375R cells were treated for 24 h with CR-1-31-B (10 nM) or DMSO in six independent biological replicates per condition across two experimental batches. RNA extraction, library preparation, sequencing, alignment, and counting were performed as described above for the input-RNA libraries. Genes with at least 10 counts in at least three libraries were retained. Differential expression was analyzed with DESeq2 v1.48.2 in R v4.5.1 using the design ∼ batch + condition. P values were adjusted by the Benjamini-Hochberg method. Both unshrunk log_2_ fold changes and normal-prior-shrunken log_2_ fold changes were retained; shrunken effects were used where indicated for visualization and ranked analyses.

### Quantitative Proteomics

#### Experimental design, sample preparation, and acquisition

Quantitative proteomics included four biological replicates in each of four groups: A375-DMSO, A375-CR-1-31-B, A375R-DMSO, and A375R-CR-1-31-B. Cells were treated for 36 h with CR-1-31-B (10 nM) or matched DMSO.

Sample preparation was performed by the Université de Sherbrooke Mass Spectrometry Proteomics Platform. Cell pellets were suspended in 50 mM triethylammonium bicarbonate (TEAB; pH 8.5), mixed 1:1 with 2× lysis buffer (10% SDS, 100 mM TEAB, pH 8.5), heated (95°C, 5 min), sonicated, and centrifuged (16,000 × g, 5 min). Protein was quantified by BCA assay. Each 50-µg aliquot was brought to 92 µL with 1× lysis buffer (5% SDS, 50 mM TEAB, pH 8.5), reduced with 4 µL of 120 mM tris(2-carboxyethyl)phosphine (5 mM final; 55°C, 15 min), and alkylated with 4 µL of 500 mM chloroacetamide (20 mM final; 10 min, room temperature, dark).

Samples were acidified with 15 µL of 27% phosphoric acid, diluted with 690 µL of 90% methanol containing 100 mM TEAB, and loaded onto Monarch S1A spin columns (New England Biolabs). After three 150-µL washes, proteins were digested overnight at 37°C with 5 µg Pierce MS-grade trypsin in 20 µL of 200 mM TEAB. Peptides were sequentially eluted with 60 µL of 50 mM TEAB, 60 µL of 0.2% formic acid, and 60 µL of 50% acetonitrile. Pooled eluates were vacuum-dried, reconstituted in 200 µL of 1% formic acid in LC–MS-grade water, quantified at 205 nm by NanoDrop, and stored at −80°C.

Peptides (375 ng/injection) were separated on a Bruker nanoElute 2 HPLC with an Acclaim PepMap 100 C18 trap (0.3 × 5 mm) and PepSep C18 analytical column (1.9 µm, 75 µm × 25 cm) using a 2-h 5–37% acetonitrile gradient in 0.1% formic acid at 400 nL/min, and analyzed on a Bruker timsTOF Pro with a CaptiveSpray source. diaPASEF acquisition used a target intensity of 20,000 and an intensity threshold of 2,500. Each acquisition cycle comprised one MS1 TIMS frame followed by 11 diaPASEF frames containing 27 contiguous 50-Da isolation windows spanning m/z 113.5–1,463.5 and arranged along the peptide m/z–mobility diagonal. TIMS accumulation and ramp times were 100 ms, the cycle time was approximately 1.27 s, and collision energy varied with ion mobility from approximately 21.8 to 54.5 eV.

Raw DIA files were processed using DIA-NN v2.2.0 Academia^67^ in a two-pass workflow employing an initial predicted spectral library, followed by generation of an experiment-specific spectral library from all 16 runs and reanalysis of the data. Searches were performed against the UniProt human reference proteome UP000005640 downloaded on August 3, 2025. In silico digestion specified cleavage after lysine or arginine, with up to one missed cleavage; peptides of 7–30 amino acids and precursor charge states of 2–4 were considered. Cysteine carbamidomethylation was specified as a fixed modification, and N-terminal methionine excision was enabled. Match-between-runs was enabled. Final quantitative matrices were generated using 1% precursor- and protein-group FDR thresholds. Protein inference used proteotypic peptides, and protein groups and gene annotations were assigned by DIA-NN using the UniProt FASTA annotations.

#### Proteomic differential-abundance analysis

Gene-level protein-abundance values exported from DIA-NN were analyzed in R. Intensities were transformed as log_2_(intensity + 1), and features detected in at least 50% of samples were retained (6,015 of 6,266 gene groups). Missing values were not imputed for differential-abundance analysis. Differential abundance was assessed using limma^68^ for A375R versus A375 under DMSO and for CR-1-31-B versus DMSO in A375R cells. P values were adjusted using the Benjamini–Hochberg method. Unless otherwise specified, features with FDR < 0.05 were considered differentially abundant; volcano-plot annotation and strict cross-comparison subsets additionally required |log_2_ fold change| ≥ 0.5. For principal-component analysis, features quantified in all samples included in each plot were retained, log_2_-transformed, and median-centered within each sample. PCA was performed using prcomp with feature centering but without unit-variance scaling. No missing-value imputation or batch correction was applied to the PCA.

### Public-annotation 5′UTR feature analysis

To determine whether 5′UTR architecture was associated with the response to CR-1-31-B, we selected one representative publicly annotated protein-coding transcript per gene, measured sequence and predicted structural features of its 5′UTR, and compared these features among transcripts with different 1-h translational responses.

Transcript annotations and sequences were obtained from the GENCODE v50 GTF and transcript FASTA files. Representative transcripts were selected using the following priority order: MANE Select, Ensembl canonical, GENCODE Primary, APPRIS principal, basic, MANE Plus Clinical, and other protein-coding transcripts. Ties were resolved using GENCODE evidence level, coding-sequence length, and transcript identifier. A representative 5′UTR was identified for 12,145 of the 13,548 genes in the 1-h CR-1-31-B translatomic dataset; 12,110 of these representative transcripts were MANE Select. Because transcription-start-site usage was not measured directly in A375R cells, this analysis describes features of reference 5′UTRs and not necessarily the exact transcript isoforms expressed in these cells. Complete gene-level 1-h CR-1-31-B response data and the corresponding representative-transcript annotations are provided in Supplementary Table S3B.

For each representative 5′UTR, we calculated length, G and GC content, total purine content, and purine content within the first and last 50 nucleotides. We also measured the length and density of A/G tracts and the fraction of overlapping 10-nucleotide windows containing at least 80% purines. Overlapping occurrences of (AG)4, (GA)4, (GGC)4, and (CGG)4 were counted. Two permissive G-quadruplex motif definitions were evaluated: four G tracts containing at least two or at least three consecutive G residues, with adjacent tracts separated by loops of 1–7 nucleotides. G4Hunter scores^69^ were calculated in 25-nucleotide sliding windows; the maximum score and the fractions of windows reaching thresholds of 1.2 or 1.5 were retained. Additional annotations included upstream AUGs, complete candidate upstream open reading frames, candidate open reading frames extending beyond the annotated 5′UTR, and 5′TOP-like starts, defined as an initial cytosine followed by a pyrimidine tract of at least five nucleotides.

Predicted conventional RNA secondary structure was evaluated using ViennaRNA 2.7.2^70^. Minimum free energy (MFE) was calculated in 100-nucleotide windows advanced in 50-nucleotide steps, with an additional terminal window ending at the 3′ end when needed. For 5′UTRs shorter than 100 nucleotides, the complete sequence was folded. The MFE of the first and last windows and the minimum and mean MFE across all windows were recorded. For 5′UTRs of 500 nucleotides or less, we also calculated full-sequence MFE and MFE normalized per 100 nucleotides. ViennaRNA RNA.fold was run with default thermodynamic settings. This analysis did not model pseudoknots or G-quadruplexes; G-quadruplex propensity was evaluated separately using the sequence-based measures described above.

Genes were then grouped according to their 1-h response to CR-1-31-B. TE-down and TE-up genes were defined using the Anota2Seq regulatory-mode criteria described above. Genes with decreased or increased polysome-associated RNA required FDR < 0.05 and an effect of at least log2(1.2) in the corresponding direction. A high-confidence downregulated group comprised genes that decreased at both the TE and polysome-associated-RNA levels. We compared TE-down genes with TE-up and TE-nonresponsive genes, polysome-associated-RNA-down genes with polysome-associated-RNA-up and polysome-nonresponsive genes, and the high-confidence downregulated group with genes stable at both levels. Genes assigned to the opposite response direction were excluded from the corresponding nonresponsive comparator.

We first compared continuous 5′UTR features between response groups using two-sided Wilcoxon rank-sum tests and summarized effect size using rank-biserial correlation. Sparse sequence motifs were analyzed primarily as present or absent using Fisher’s exact tests. We then used binomial logistic regression (stats::glm in R 4.5.1) to determine whether each feature remained associated with response-group membership after accounting for potential confounders. Models of TE regulation were adjusted for mean baseline DMSO total-RNA abundance, whereas models of polysome-associated RNA or the combined TE-plus-polysome response were adjusted for mean baseline DMSO polysome-associated-RNA abundance. Models also included log1p-transformed 5′UTR length and GC fraction, except when length or GC fraction was the feature being tested. Continuous predictors were standardized to z scores, binary motifs were coded as present or absent, and complete-case observations were used. Benjamini–Hochberg correction was applied separately across features within each comparison. Feature extraction was performed in Python 3.9.6; data processing and visualization used dplyr 1.2.0 and ggplot2 4.0.2. Complete adjusted association results are provided in Supplementary Table S3A.

### Gene-set enrichment and over-representation analyses

We used preranked gene-set enrichment analysis (GSEA) to identify pathways associated with each regulatory layer. Preranked GSEA^71^ was performed in R using fgsea 1.34.2 and the fgseaMultilevel algorithm. Hallmark and Reactome gene sets from MSigDB 2026.1.Hs^72^ were retrieved using msigdbr 26.1.0. Full, unthresholded gene lists were ranked using Anota2Seq APV effect estimates for the input-RNA abundance, translation-efficiency, and polysome-associated-RNA layers; DESeq2 shrunken log_2_ fold changes for the independent 24-h total RNA-seq dataset; or limma log_2_ fold changes for proteomics. Unless otherwise specified, gene-set sizes were restricted to 10–500 genes and eps was set to 0. Benjamini–Hochberg FDR < 0.05 defined significant enrichment. To reduce redundancy for visualization, significant Reactome pathways were collapsed using fgsea::collapsePathways with a p-value threshold of 0.05 and 2,000 permutations. Focused mitochondrial analyses used the MSigDB Hallmark oxidative-phosphorylation gene set and respiratory-complex-I genes from MitoCarta 3.0 ^73^.

Resistance-associated protein signatures were also evaluated as custom gene sets. Resistance-up and resistance-down signatures comprised 225 and 346 proteins, respectively, that differed between A375R and A375 cells at FDR < 0.05 and log_2_ fold change ≥ 0.5 or ≤−0.5. These signatures were tested against the complete proteomic response to CR-1-31-B using fgseaMultilevel with gene-set sizes of 10–5,000 and eps = 0.

For over-representation analysis, the query set comprised 65 proteins that were increased in A375R relative to A375 cells and decreased following CR-1-31-B treatment, with both changes meeting FDR < 0.05 and |log2 fold change| ≥ 0.5. The background comprised all 5,259 proteins quantified in both proteomic comparisons. Hallmark and Reactome pathways were tested using the upper-tail hypergeometric distribution. Pathways were required to contain at least five background proteins and overlap the query set by at least two proteins. P values were adjusted separately within each collection using the Benjamini–Hochberg method, and FDR < 0.05 defined significant over-representation.

### Cross-layer integration and resistance-program analyses

To compare the RNA and protein datasets, we first harmonized gene identifiers at the gene-symbol level. Ensembl gene IDs in the input and polysome-associated RNA count matrices were mapped using the external_gene_name annotation supplied with the RNA-seq analysis, and counts from multiple Ensembl IDs mapping to the same gene symbol were summed before Anota2Seq analysis. The corresponding external_gene_name annotation was used for the independent total RNA-seq dataset. Proteomic gene groups were assigned gene symbols from the UniProt human reference proteome UP000005640, downloaded on August 3, 2025. For cross-omic comparisons, gene symbols were converted to uppercase and matched exactly; entries without a valid gene symbol were excluded. No additional gene-alias remapping database was applied.

Cross-layer relationships were evaluated using Spearman rank correlations across all genes quantified in both layers. The A375R-versus-A375 analyses included 5,188 matched genes for both the polysome-associated-RNA/protein and TE/protein comparisons. Statistical significance was determined independently within each layer for display, but the correlations were calculated across all matched genes rather than only the significant subsets.

To visualize the CR-1-31-B response across regulatory layers, we constructed a heatmap of 3,241 genes that were quantified for TE at 1 h, total RNA at 24 h, and protein at 36 h and met the selection criteria in at least one layer. We applied the Anota2Seq criteria described above for TE, FDR < 0.05 and |shrunken log₂ fold change| ≥ 0.5 for total RNA, and FDR < 0.05 and |log₂ fold change| ≥ 0.5 for protein. For clustering, each effect was divided by its layer-specific effect-size threshold [log₂(1.2) for TE and 0.5 for total RNA and protein]. Values were set to zero when a gene did not meet the selection criteria in a given layer, and scaled values were capped at −6 and +6 before clustering. Six clusters were defined using Euclidean distance and Ward.D2 linkage.

To determine whether CR-1-31-B opposed protein changes associated with acquired resistance, we restricted the analysis to proteins quantified in both the A375R-versus-A375 comparison and the CR-1-31-B-versus-DMSO comparison in A375R cells. Resistance-up and resistance-down proteins required FDR < 0.05 and a log₂ fold change ≥ 0.5 or ≤ −0.5, respectively; CR-1-31-B-responsive proteins required FDR < 0.05 and |log₂ fold change| ≥ 0.5. Of the 225 resistance-up proteins, 222 had corresponding measurements in all three CR-1-31-B-response layers—TE at 1 h, total RNA at 24 h, and protein at 36 h. These 222 genes were ordered by hierarchical clustering of their threshold-scaled CR-1-31-B effects.

### Public melanoma PDX treatment-trajectory analysis

To determine whether the A375R response to CR-1-31-B opposed gene programs that emerge during BRAF/MEK-targeted therapy, we analyzed processed spatial-transcriptomic data from the BRAF^V600E^ melanoma PDX models WM4007 and WM4237. Data were obtained from GSE245582 and the associated Visium objects deposited at Zenodo (doi: 10.5281/zenodo.10287542)20. The dataset included 24 tissue sections. Twenty sections represented five treatment stages (T0–T4) in each PDX model, with two nearby 10x Visium sections (S1 and S2) obtained from each tumor block. The remaining four were vehicle-control (TC) sections: two from a WM4007 vehicle-control tumor collected on day 38 and two from a WM4237 vehicle-control tumor collected on day 42. Each PDX model and time point was represented by one tumor from one mouse.

The author-provided objects contained log1p-transformed, sample-scaled expression values after mouse/stromal spots had been removed and the data restricted to human genes. We recovered linear values using expm1, summed expression across the retained human-tumor spots within each section, and normalized the resulting values to counts per million (CPM). Expression estimates from the two nearby sections of each tumor were assessed for concordance and then averaged with equal weight to obtain one tumor-level estimate; the sections were not treated as independent biological replicates. We did not calculate spot- or section-level differential-expression P values.

The TC tumor-level profiles were used only in sensitivity analyses. Vehicle-control drift was assessed separately within each model by comparing the WM4007 TC tumor at day 38 with WM4007 pretreatment T0 at day 0 and the WM4237 TC tumor at day 42 with WM4237 pretreatment T0 at day 0. Genes with an absolute TC-versus-T0 change ≥ 0.75 log₂ units were classified as showing large vehicle-control drift.

The WM4007 trajectory comprised pretreatment (T0, day 0), early response (T1, day 14), maximal response/persistence-MRD (T2, day 44), early regrowth (T3, day 73), and resistant regrowth (T4, day 133). The WM4237 trajectory comprised pretreatment (T0, day 0), early treated/early persister (T1, day 14), early and late persistence/MRD (T2, day 42; T3, day 70), and resistant regrowth (T4, day 94). Both trajectories were sampled during continuous dabrafenib/trametinib treatment after baseline. Stage effects were calculated as log₂ ratios using a 0.5-CPM pseudocount. For WM4237, the T2 and T3 profiles were averaged with equal weight to define persistence before comparison with T0. Resistant-regrowth-state effects compared T4 with T0, whereas regrowth-transition effects compared T4 with the model-specific persistence profile. Sustained effects captured changes shared by persistence and resistant regrowth relative to baseline: when both effects had the same direction, we retained the smaller absolute effect with its sign; when their directions differed, the sustained effect was set to zero.

Genes were included in ranked analyses if they reached at least 1 CPM and were detected in at least 5% of retained spots at one or more included stages. A stage-induced program comprised genes with a log₂ effect ≥ 0.5 and a concordant effect direction in the paired S1 and S2 tissue sections from the same tumor block. For each PDX model separately, we then asked whether stage-induced genes were shifted toward lower protein abundance in the A375R CR-1-31-B response. One-sided Mann–Whitney rank tests were performed across the complete 5,259-gene proteomic universe after intersection with genes passing the PDX filters. AUROC was calculated as U/(n1 × n0); 0.5 indicates no shift, whereas values > 0.5 indicate enrichment toward a CR-1-31-B-associated protein decrease. We applied Benjamini–Hochberg correction across the family of stage-program rank-enrichment tests. The fraction of each PDX-up program with a CR-1-31-B protein log₂ fold change < 0 was reported descriptively.

To identify persistence-associated changes shared across the two PDX models, genes were required to pass the expression filters in both models, show direction-consistent effects between the paired S1 and S2 sections from the same tumor, change in the same direction in both models, and be measured in the A375R CR-1-31-B proteome. For direction-concordant genes, the signed conservative persistence score retained the smaller absolute effect observed in the two PDXs. Its association with the 36-h CR-1-31-B protein log₂ fold change was assessed by Spearman correlation. Persistence-up genes required a log₂ effect ≥ 0.5 in each PDX, whereas stringent CR-1-31-B protein decreases required a log₂ fold change ≤ −0.5 and FDR < 0.05. Sensitivity analyses used PDX thresholds ranging from 0.25 to 1.0, alternative methods for summarizing the two PDX effects, exclusion of genes with large vehicle-control drift, and pseudocounts ranging from 0.1 to 1 CPM. Because each PDX stage was represented by one tumor, these analyses describe cross-dataset rank structure and do not provide biological replication of the PDX treatment stages.

For the curated matrix in Fig. S4E, genes were required to show a log₂ effect ≥ 0.5 in both PDX models at one or more treatment stages defined independently within each trajectory and to encode proteins significantly decreased after CR-1-31-B treatment (log₂ fold change ≤ −0.5; FDR < 0.05). Among the 195 proteins meeting these cross-PDX induction and CR-1-31-B-response criteria, 16 representative proteins were manually selected to balance biological interpretability, treatment-stage coverage, and representation across four descriptive functional categories: signaling/stress survival, melanoma state/adhesion, metabolism, and cell cycle/regrowth. The display is non-exhaustive and does not represent a statistical ranking.

### Seahorse extracellular-flux assays

A375 and A375R cells were seeded at 4 × 10³ cells per well in Seahorse XFe96 culture plates in 100 µL complete medium. After 24 h, cells were treated for 24 h with DMSO, PLX4032 (2 µM), CR-1-31-B (10 nM), or the combination, unless stated otherwise. Oxygen-consumption rate (OCR) and extracellular-acidification rate (ECAR) were measured with a Seahorse XFe96 analyzer (Agilent Technologies) according to the manufacturer’s instructions. Normalization was done by protein content.

Mitochondrial-stress assays used oligomycin (1 µM), FCCP (1.5 µM), and rotenone plus antimycin A (0.5 µM each). Glycolytic-stress assays used glucose (10 mM), oligomycin (1 µM), and 2-deoxyglucose (50 mM). ATP production rates were measured using dedicated Seahorse XF Real-Time ATP Rate Assays and calculated as described by Papadopoli et al.^74^ and according to the manufacturer’s instructions. ATP-rate values were normalized to protein abundance. Three independent biological experiments were performed; technical wells (generally six per condition) were averaged within each experiment before statistical analysis.

Mito Fuel Flex assays were performed in untreated A375 and A375R cells using UK5099 (2 µM), BPTES (3 µM), and etomoxir (4 µM) to interrogate glucose/pyruvate, glutamine, and fatty-acid oxidation, respectively. Dependency, capacity, and flexibility were calculated according to the manufacturer’s equations. Paired biological-experiment comparisons between A375 and A375R were evaluated with two-sided paired t tests and Holm adjustment across the three fuels.

Derived parameters were paired by biological experiment to the corresponding DMSO condition and expressed as log_2_ fold changes for the descriptive matrix. One negative spare-respiratory-capacity estimate could not be represented as a log ratio; it was retained in the source table but excluded from that matrix. Total ATP-production-rate data were analyzed in R 4.5.1 using a two-factor repeated-measures ANOVA, with cell line and treatment as within-experiment factors and biological experiment as the blocking factor. Prespecified comparisons were performed using two-sided paired t-tests, with Holm adjustment for multiple comparisons.

### Steady-state GC-MS

#### Steady-state GC-MS sample preparation and acquisition

A375 and A375R cells were seeded at 2.5 × 10⁵ cells per well in six-well plates containing 3 mL complete medium. After 24 h, A375 cells were treated for 24 h with DMSO, PLX4032 (2 µM), or CR-1-31-B (10 nM), and A375R cells were treated for 24 h with DMSO, PLX4032 (2 µM), or PLX4032 plus CR-1-31-B. Cells were washed three times with ice-cold saline (9 g/L NaCl) and quenched twice with 600 µL 80% methanol maintained below −20°C. Cells were scraped into prechilled tubes, sonicated at 4°C for 20 min (30 s on/30 s off, high setting; Bioruptor Plus with Minichiller, Diagenode), and clarified by centrifugation at 14,000 rpm for 10 min at 4°C. Supernatants were dried overnight at −4°C in a CentriVap refrigerated concentrator (Labconco, 7310022).

Dried extracts were resuspended in 30 µL pyridine (Sigma-Aldrich, 270970) containing 10 mg/mL methoxyamine hydrochloride (Sigma-Aldrich, 226904), mixed by sonication and vortexing, clarified, and incubated for 20 min at room temperature. Samples were derivatized with 70 µL N-tert-butyldimethylsilyl-N-methyltrifluoroacetamide (MTBSTFA; Sigma-Aldrich, 394882) for 1 h at 70°C. One microliter was injected into an Agilent 5977B GC/MSD coupled to an Intuvo 9000 GC and 7693A autosampler. Myristic-d27 was used as an internal standard. Chromatographic and selected-ion-monitoring settings were as previously described^45^.

Three independent biological experiments were analyzed, each with three technical injections per condition. SCAN, selected-ion-monitoring (SIM), and diluted-SCAN acquisitions represented alternative measurements of the same samples and were not treated as independent replicates. A feature-level acquisition mode was selected according to prespecified platform quality-control rules: valid SIM was preferred across all experiments, followed by undiluted SCAN, with diluted SCAN used when the undiluted signal was saturated or outside the linear range.

#### Steady-state GC-MS data processing and statistics

Each technical measurement was divided by the matched myristic-d27 signal and the cell number determined from parallel wells, after which technical injections were averaged within each biological experiment. Of 78 initially considered metabolites, 56 passed complete-measurement criteria across the three experiments and were modeled; tentative identities were retained only when explicitly flagged and were excluded from focused analyses. For focused and full supplementary matrices, features additionally required a median technical coefficient of variation ≤30%.

Normalized abundances were log₂-transformed and analyzed with limma using the design ∼ experiment + condition, with experiment included as a fixed effect to account for between-experiment variation. Empirical-Bayes moderation with trend was applied, and P values were adjusted within each contrast by the Benjamini-Hochberg method. Prespecified contrasts were A375 PLX4032/DMSO, A375R PLX4032/DMSO, A375 CR-1-31-B/DMSO, and A375R PLX4032+CR-1-31-B/PLX4032. Because not all four treatment arms were available in both cell lines for steady-state metabolomics, no formal PLX4032-by-CR-1-31-B interaction was estimated. GC-MS PCA used 53 curated, non-tentative features with median technical CV ≤ 30% and complete measurements. Biological-experiment means were log_2_-transformed, mean-centered across conditions within experiment, and autoscaled before PCA.

### Steady-state LC-MS

#### Steady-state LC-MS sample preparation and acquisition

LC–MS-grade water, acetonitrile, methanol, formic acid, ammonium acetate, and ammonium formate were obtained from Fisher Scientific; authentic metabolite standards and N-ethylmaleimide (NEM) were obtained from Sigma-Aldrich. A375 and A375R cells were seeded at 2.5 × 10⁵ cells per well in six-well plates containing 3 mL complete medium and treated for 24 h as follows: A375 cells with DMSO, PLX4032 (2 µM), or CR-1-31-B (10 nM), and A375R cells with DMSO, PLX4032 (2 µM), or PLX4032 (2 µM) plus CR-1-31-B (10 nM).

Cells were washed three times with ice-cold 150 mM ammonium formate containing 1 mg/mL NEM (Sigma-Aldrich, E3876), adjusted to pH 7.4 with ammonium hydroxide. NEM was included to limit oxidation of thiol metabolites, as described previously^75^. Cells were scraped into 380 µL of prechilled 1:1 methanol/water containing 1 mg/mL NEM, after which 220 µL of ice-cold acetonitrile was added. Samples were disrupted for 30 s at 50 Hz in a SpeedMill Plus (Analytik Jena) with four 2-mm ceramic beads.

For liquid–liquid extraction, 600 µL dichloromethane and 300 µL water were added. Samples were centrifuged, and the upper aqueous phase was collected and evaporated in a chilled vacuum concentrator at −4°C (Labconco). Dried extracts were reconstituted in 50 µL chilled water, clarified by centrifugation at 1°C, and 5 µL was injected for each analysis.

Metabolites were analyzed on an Agilent 6470 triple-quadrupole mass spectrometer coupled to an Agilent 1290 Infinity quaternary UHPLC system. Data were acquired in negative-ion mode using a Jet Stream electrospray ionization source with a drying-gas temperature of 150°C, gas flow of 13 L/min, nebulizer pressure of 45 psi, and capillary voltage of 2,000 V. Multiple-reaction-monitoring qualifier and quantifier transitions and retention times were established and optimized using authentic standards.

Chromatographic separation was performed at 35°C on a Zorbax Extend C18 column (1.8 µm, 2.1 × 150 mm) fitted with a matching guard column (1.8 µm, 2.1 × 5 mm). Mobile phase A contained 97% water, 3% methanol, 10 mM tributylamine, 15 mM acetic acid, and 5 µM medronic acid; mobile phase C was methanol containing the same additives; and mobile phase D was 90% acetonitrile. At 0.25 mL/min, the gradient was held at 100% A for 2.5 min, increased to 20% C over 5 min, to 45% C over the next 5.5 min, and to 99% C over the next 7 min, followed by a 4-min hold at 100% C.

The column was washed with 99% D for 3 min at 0.25 mL/min; the flow was then increased to 0.8 mL/min over 0.5 min, held for 3.85 min, and reduced to 0.6 mL/min over 0.15 min. The system was returned to 100% A over 0.75 min while the flow was lowered to 0.4 mL/min, held for 7.65 min, and returned to 0.25 mL/min 1 min before the next injection. Peaks were integrated using MassHunter Quantitative Analysis software (version B.10.00; Agilent).

Three independent biological experiments were analyzed per condition. Peak signals were normalized to cell numbers determined from matched parallel wells before technical cultures were averaged within each biological experiment. The curated analysis input consisted of experiment-matched log_2_ ratios relative to the corresponding DMSO condition.

#### Steady-state LC–MS analysis and visualization

For each LC-MS metabolite and contrast, the mean log_2_ fold change across the three biological experiments was calculated and tested against zero using a two-sided one-sample t test. Benjamini-Hochberg adjustment was performed separately within each contrast. The A375R PLX4032+CR-1-31-B/PLX4032 effect was calculated by subtracting the experiment-matched PLX4032/DMSO log_2_ ratio from the PLX4032+CR-1-31-B/DMSO log_2_ ratio. Directional consistency across experiments was retained in the source tables.

The full supplementary matrices preserve platform-specific feature identities and statistics. Cross-model concordance compared A375 CR-1-31-B/DMSO with A375R PLX4032+CR-1-31-B/PLX4032 using Spearman correlation and directional concordance; this comparison is descriptive because the two contrasts were generated in distinct cellular/treatment contexts.

### Glutamine–TCA pathway protein–metabolite display

For Fig. S5L and Supplementary Table S5, 18 proteins associated with glutamine utilization, TCA-cycle reactions and related carbon-shuttle pathways were included based on biochemical relevance, independently of response direction or statistical significance. Protein log_2_ fold changes and Benjamini–Hochberg-adjusted FDR values were taken from the 36-h A375R CR-1-31-B-versus-DMSO limma analysis (n = 4 biological replicates per condition). Nine measured glutamine–TCA metabolites were included from the 24-h steady-state GC-MS comparison of PLX4032 plus CR-1-31-B versus PLX4032 in A375R cells (n = 3 independent biological experiments per condition); metabolite FDR values were adjusted within the GC-MS platform and contrast. Succinyl-CoA and oxaloacetate were included as unquantified pathway intermediates. Pathway arrows show biochemical topology rather than measured flux. Because the protein and metabolite layers derive from different time points and treatment contrasts, their joint presentation is descriptive and does not establish direct cross-layer mediation.

### [U-¹³C₅]glutamine tracing

For isotope tracing, cells were seeded and pretreated as for steady-state GC-MS. After the 24-h drug exposure, cells were equilibrated for 2 h in complete medium lacking unlabeled glutamine and then supplied with 4 mM [U-¹³C₅]glutamine (Cambridge Isotope Laboratories, CLM-1822-M-0.25). Samples were collected after 0.5, 1, 2, or 6 h and extracted, derivatized, and analyzed by GC-MS as described above. All four treatment conditions were analyzed in both A375 and A375R cells in two independent biological experiments. Drug treatments remained present throughout equilibration and tracing.

Raw isotopologue distributions were corrected for natural isotope abundance using a correction matrix. Fractional enrichment was calculated as the corrected abundance of each isotopologue divided by the sum of all corrected isotopologues for that metabolite. Selected direct-entry trajectories comprised glutamine, glutamate, and α-ketoglutarate m+5; oxidative-TCA output comprised succinate, fumarate, malate, and aspartate m+4; citrate-family m+4 and m+5 were used to summarize oxidative and reductive labeling, respectively.

For metabolite-pool/isotopologue decomposition, raw isotopologue signals were summed for each metabolite, divided by the matched myristic-d27 internal-standard signal and the cell number determined from parallel wells, and expressed relative to the DMSO pool within each biological experiment. Glutamine and glutamate used the matching diluted-SCAN internal-standard measurement; undiluted metabolites used the SCAN internal-standard measurement. The 1-h glutamine-to-glutamate transfer index was calculated within each experiment as glutamate m+5 fractional enrichment divided by glutamine m+5 fractional enrichment. This index reflects transfer of supplied labeled carbon into glutamate and is not a direct measure of absolute glutaminase flux or net glutamine uptake. Because n = 2, tracing results were treated as descriptive and no inferential statistics were applied.

### Extracellular nutrient measurements

Glucose and glutamine concentrations were measured in conditioned medium collected after 24 h using a BioProfile 400 analyzer (Nova Biomedical) at the Rosalind and Morris Goodman Cancer Institute Metabolomics Innovation Resource (MIR), as previously described^76^. Instrument outputs were reported in g/L for glucose and mmol/L for glutamine. Matched cell-free medium controls incubated in parallel were used to calculate nutrient consumption. Concentration changes were converted to micromoles using the 2-mL culture volume and normalized to the logarithmic mean cell number between treatment initiation and sample collection. Three technical wells were averaged within each of six independent biological experiments. Prespecified comparisons—A375 DMSO versus CR-1-31-B and A375R PLX4032 versus PLX4032+CR-1-31-B—were evaluated using two-sided paired t-tests.

### Glutamine-deprivation experiments

Glutamine-deprivation experiments used glutamine-free DMEM (Wisent BioProducts, 319-025-CL) supplemented with 10% fetal bovine serum and 1% penicillin-streptomycin. After attachment in complete medium, cells were washed once with PBS and transferred to medium containing 4, 0.25, or 0 mM glutamine, prepared with L-glutamine (Multicell, 609-065-EL). For metabolomics, cells were cultured under these conditions for 24 h and processed by GC-MS. Mean log2 fold changes relative to the matched 4 mM condition were calculated across three independent biological experiments; two-sided one-sample t tests were adjusted across measured metabolites within each cell-line/concentration comparison by the Benjamini-Hochberg method.

For endpoint viable-cell-number assays, A375 and A375R cells were cultured for 72 h at the indicated glutamine concentrations with vehicle or CR-1-31-B (3.5 nM). Cells were counted by trypan-blue exclusion. Three technical counts were averaged within each of three independent biological experiments, and values were normalized to the matched vehicle-treated 4 mM glutamine condition.

### A375 Xenograft Study

#### Animals

Female athymic nude mice [Crl:NU(NCr)-Foxn1nu; Charles River Laboratories, strain code 490] were 8 weeks old at the start of the study. Mice were housed under standard specific pathogen-free conditions in a temperature- and humidity-controlled facility with a 12-h light/dark cycle and ad libitum access to standard rodent chow and water. Animal studies were conducted at the animal facility of the Maisonneuve-Rosemont Hospital Research Centre and approved by the institutional animal care committee (Comité de protection des animaux, Santé Québec – Est-de-l’Île-de-Montréal) under protocol 2021-2606. A375 cells (1 × 10⁶) were suspended in 200 µL PBS and injected subcutaneously into the right flank. Tumors were monitored until they reached approximately 100-150 mm³, after which mice were allocated randomly to vehicle, PLX4720, CR-1-31-B, or combination treatment.

#### Drug administration and monitoring

PLX4720 (Focus Biomolecules, 10-5113-BULK) was administered at 45 mg/kg in a volume of 100 µL by oral gavage daily on treatment days 1–5 and every other day thereafter. CR-1-31-B (MedChemExpress, HY-136453-10MG) was administered intraperitoneally at 0.5 mg/kg every other day through day 5, followed by 0.4 mg/kg every other day thereafter in a dosing volume of 100 µL. Control animals received matched vehicle administrations. CR-1-31-B was formulated in a vehicle containing 5.2% (v/v) PEG 400 and 5.2% (v/v) Tween 80 in saline. PLX4720 was formulated in a vehicle containing 60% (v/v) Phosal 50 PG, 30% (v/v) PEG 400, and 10% (v/v) ethanol. On combination-treatment days, PLX4720 was administered by oral gavage in the morning, whereas CR-1-31-B was administered by intraperitoneal injection in the afternoon.

Tumor dimensions were measured with digital calipers every other day, and volume was calculated as V = (π/6) × W² × L, where W and L represent tumor width and length, respectively. Body weight and clinical condition were monitored three times per week. Humane endpoints included a maximum tumor volume of 1,800-2,000 mm³, tumor ulceration, or any other protocol-defined welfare criterion requiring euthanasia, as determined by animal-care or veterinary staff. A recorded value of zero represented a nonpalpable tumor or complete regression. Blank observations after an animal left the study were treated as missing and were not imputed.

The displayed efficacy cohorts comprised vehicle n = 5, PLX4720 n = 11, CR-1-31-B n = 8, and PLX4720+CR-1-31-B n = 11. Individual mice were the biological unit; repeated caliper measurements were longitudinal observations and were not treated as independent replicates.

#### Longitudinal tumor and endpoint analyses

Longitudinal comparisons used comparison-specific evaluable windows: through day 13 for vehicle versus CR-1-31-B, the last time point at which all five vehicle-treated mice remained under observation, and through day 49 for PLX4720 versus the combination; at day 49, nine PLX4720-treated and 11 combination-treated mice remained under observation. Individual trajectories were displayed through day 75 without imputation after an animal ceased contributing.

Longitudinal tumor growth was compared separately between vehicle and CR-1-31-B through day 13 and between PLX4720 and combination therapy through day 49. For each prespecified comparison, log1p-transformed tumor volumes were analyzed using a linear mixed-effects model. Treatment, treatment day (categorical), and their interaction were included as fixed effects; a mouse-specific random intercept and an AR(1) correlation structure accounted for repeated measurements within mice. Reported P values test whether the tumor-growth trajectories differed between treatments, based on the treatment-by-day interaction. Analyses were performed in R 4.5.1 using nlme 3.1-168.

Time to tumor endpoint was measured from treatment initiation. An event was defined as euthanasia after reaching a protocol-defined tumor-related humane endpoint, including excessive tumor burden or tumor ulceration. Animals remaining below these criteria at administrative study termination were censored at their last follow-up. Animals lost for reasons unrelated to tumor progression were excluded from the efficacy analysis. Kaplan-Meier curves were compared by two-sided log-rank tests. Body-weight change was calculated relative to each animal’s treatment-day-1 weight; observed group means were shown only while at least five animals contributed.

### Statistical Analysis and Reproducibility

Biological experiments or individual mice were the unit of replication. Technical replicates were averaged within each biological experiment before inferential analysis unless explicitly stated otherwise. Sample sizes, summary statistics, and panel-specific tests are reported in the figure legends. All tests were two-sided unless specified. Multiple-testing correction used the Benjamini-Hochberg false-discovery-rate procedure for omics and metabolomics analyses and the Holm or Tukey method for predefined low-dimensional comparisons, as indicated.

One-way ANOVA followed by Tukey’s honestly significant difference test was used for the displayed dose-versus-vehicle apoptosis and clonogenic comparisons within each cell line. Dose-response curves were fitted by nonlinear least squares using stats::nls. Analyses and figure generation were performed in R version 4.5.1 using ggplot2 4.0.2, svglite 2.2.2, limma 3.64.3, edgeR 4.6.3, DESeq2 1.48.2, anota2seq 1.30.0, fgsea 1.34.2, nlme 3.1-168, and survival 3.8-6. Exact P values are reported where practical; P < 0.05 was considered statistically significant for low-dimensional analyses. No inferential statistics were applied to the n = 2 isotope-tracing experiments.

## Data Availability

The sequencing data generated in this study have been deposited in NCBI’s Gene Expression Omnibus (GEO)^77^ under Series accession GSE344859.

The mass spectrometry proteomics data have been deposited to the ProteomeXchange Consortium via the PRIDE^78^ partner repository with the dataset identifier PXD083257.

## Acknowledgments

This work was supported by the Canadian Institutes of Health Research (CIHR; grant 165901 to L.H.), research funds associated with a Fonds de recherche du Québec – Santé (FRQS) Junior 1 Research Scholar Award to L.H., and start-up funds from the Centre de recherche de l’Hôpital Maisonneuve-Rosemont (CR-HMR) and the Faculty of Medicine, Université de Montréal (to L.H.). L.H. is a Fonds de recherche du Québec – Santé (FRQS) Junior 2 Research Scholar. A.S.C. was supported by postdoctoral fellowships from the Fondation de l’Hôpital Maisonneuve-Rosemont and the FRQS (https://doi.org/10.69777/307595). M.T. was supported by a Canada Graduate Scholarship–Master’s from the Canadian Institutes of Health Research (CIHR), a Scientific Outreach Scholarship and a Merit Scholarship (A21) from the Faculty of Medicine of the Université de Montréal, and a Doctoral Research Award from the Cancer Research Society (award 1511009). A.S.H.N. was supported by a doctoral research scholarship from the Fonds de recherche du Québec (https://doi.org/10.69777/2008870) and a Cole Foundation Scholarship. C.C. was supported by post-doctoral fellowships from the Cole Foundation and from Takeda Canada/CIHR Institute of Genetics Rare Diseases. S.S-A. received studentships from the FRQS (https://doi.org/10.69777/350704), the Foundation of the Maisonneuve-Rosemont Hospital, the Cancer Research Society, and the Leukemia & Lymphoma Society. SDS-1-021, silvestrol, and CMLD012612 were generous gifts from the late Dr. Jerry Pelletier (McGill University). We are very grateful to Ola Larsson, Christian Oertlin, and Kathleen Watt, at the Karolinska Institutet, Sweden, for their guidance on the bioinformatic analysis of the polysome-profiling data. We thank the core facilities at the Maisonneuve-Rosemont Hospital Research Centre for technical support. We acknowledge the Rosalind and Morris Goodman Cancer Institute Metabolomics Innovation Resource (MIR) for metabolomics support. The MIR is supported by the Canada Foundation for Innovation, the Dr. John R. and Clara M. Fraser Memorial Trust, the Terry Fox Foundation (Oncometabolism Team Grant 116128), the Quebec Breast Cancer Foundation, and McGill University.

## Abbreviations

5′UTR: 5′ untranslated region
α-KG: α-ketoglutarate
ANOVA: analysis of variance
APV: analysis of partial variance
AUROC: area under the receiver operating characteristic curve
BRAFi: BRAF inhibitor
CPM: counts per million
DIA: data-independent acquisition
DIA-NN: data-independent acquisition by neural networks
diaPASEF: data-independent acquisition parallel accumulation–serial fragmentation
DMSO: dimethyl sulfoxide
ECAR: extracellular acidification rate
ECM: extracellular matrix
eIF4A: eukaryotic translation initiation factor 4A
eIF4F: eukaryotic translation initiation factor 4F complex
FDR: false discovery rate
G4: G-quadruplex
GC–MS: gas chromatography–mass spectrometry
GSEA: gene set enrichment analysis
IC₅₀: half-maximal inhibitory concentration
LC–MS: liquid chromatography–mass spectrometry
MAPK: mitogen-activated protein kinase
MEK: mitogen-activated protein kinase kinase
MEKi: MEK inhibitor
MFE: minimum free energy
MRD: minimal residual disease
mTOR: mechanistic target of rapamycin
mTORC1: mechanistic target of rapamycin complex 1
NES: normalized enrichment score
OCR: oxygen consumption rate
OXPHOS: oxidative phosphorylation
PCA: principal component analysis
PDX: patient-derived xenograft
PI3K: phosphoinositide 3-kinase
RNA-seq: RNA sequencing
RT-qPCR: reverse-transcription quantitative polymerase chain reaction
SD: standard deviation
SEM: standard error of the mean
TCA: tricarboxylic acid cycle
TE: translational efficiency
TIMS: trapped ion mobility spectrometry.

## Supplementary Figure S1

**Fig. S1. Additional pharmacologic and molecular validation of eIF4A-inhibitor activity across BRAF inhibitor sensitive and resistant models. (A,B)** Puromycin incorporation after 1 h (A) or 24 h (B) in A375 cells treated with vehicle, PLX4032 (2 µM), or CR-1-31-B (10 nM), and in A375R cells treated with vehicle, PLX4032 (2 µM), or PLX4032 (2 µM) plus CR-1-31-B (10 nM). Puromycin was added for the final 15 min; β-actin served as a loading control. Quantification is relative to the corresponding vehicle control. **(C)** Puromycin incorporation and expression of the indicated signaling and survival proteins in LU1205R cells treated for 24 h with vehicle, PLX4032 (2 µM), or PLX4032 plus CR-1-31-B (10 nM). Quantification shows puromycin incorporation relative to vehicle. **(D)** PLX4032 concentration-response curve in LU1205R cells after 72 h. **(E-G)** Concentration-response curves for SDS-1-021 (E), silvestrol (F), and CMLD012612 (G) in A375 and A375R cells after 72 h. **(H-J)** Silvestrol (H), SDS-1-021 (I), and CMLD012612 (J) concentration-response curves in LU1205R cells after 72 h. **(K)** Summary of fitted IC_50_ values. Values are in nM; the A375 PLX4032 value is mean ± SD from three independently fitted experiments. The A375R PLX4032 IC_50_ was not reached within the tested range of the representative experiment. **(L)** CR-1-31-B concentration-response curve in YUMMER1.7 melanoma cells after 72 h. Data are mean ± SD from three independent experiments. **(M,N)** Immunoblot analysis of BCL-2, CDK4, and cyclin D3 in A375 and A375R cells treated for 24 h with SDS-1-021 (M) or CMLD012612 (N). **(O)** Immunoblot analysis of BCL-2, CDK4, and cyclin D3 in LU1205R cells exposed for 24 h to increasing concentrations of CR-1-31-B, silvestrol, SDS-1-021, or CMLD012612. **(P,Q)** Representative clonogenic images (P) and crystal-violet quantification (Q) of MDA-MB-231 and MDA-MB-231R cells exposed to the indicated encorafenib/binimetinib combinations for 6 days (n = 3 independent biological experiments). **(R)** CR-1-31-B concentration-response curves in MDA-MB-231 and MDA-MB-231R cells after 72 h. Viable cell numbers were determined by trypan-blue exclusion and normalized to vehicle. **(S)** Representative clonogenic images and crystal-violet quantification of MDA-MB-231 and MDA-MB-231R cells after 6 days of CR-1-31-B exposure. **(T)** Immunoblot analysis of BCL-2, CDK4, and cyclin D3 after 24 h exposure to increasing CR-1-31-B concentrations. β-actin was used as a loading control. For A–C, symbols represent three independent biological experiments and bars show mean ± SD. Within each panel and cell line, groups were compared by one-way ANOVA with Tukey adjustment; brackets indicate the displayed pairwise comparisons. For S, symbols likewise represent three independent biological experiments, and dose-versus-vehicle comparisons within each cell line were evaluated by one-way ANOVA with Tukey adjustment. Other quantitative panels show biological experiments and mean ± SD. ns, not significant; *P < 0.05; **P < 0.01; ***P < 0.001; ****P < 0.0001.

## Supplementary Figure S2

**Fig. S2. Molecular and bioenergetic features of acquired BRAFi resistance in A375R cells. (A)** Representative normalized polysome profiles from one of two experimental batches, showing three of six biological replicates per cell state for baseline A375 and A375R cells. The 80S monosome peak and heavy-polysome region are indicated. **(B)** Polysome/80S ratios across all six biological replicates per cell state. Symbols represent biological replicates and horizontal lines show means. The P value (P = 0.034) was obtained using an unpaired two-sided Welch t test on log2-transformed ratios. **(C-E)** Principal-component analysis (PCA) of batch-adjusted input/total RNA (C), polysome-associated RNA (D), and relative TE (E) from A375 and A375R cells. Colors denote cell state and symbols denote experimental batch. **(F)** PCA of quantitative proteomes from untreated A375 and A375R cells (n = 4 per cell state). **(G)** Direction-resolved numbers of selected features across RNA-abundance, translation/TE, buffering, polysome-associated RNA, and protein layers. **(H)** Concordance between TE and protein-abundance changes across 5,188 matched genes. Colored genes were significant in both layers and changed concordantly. Spearman ρ = 0.45, P < 2.2 × 10⁻¹⁶. **(I)** Mitochondrial dependency and flexibility for fatty-acid, glucose, and glutamine oxidation in untreated A375 and A375R cells, determined using the Seahorse Mito Fuel Flex Test. Stacked bars show mean dependency and flexibility; open symbols show oxidation capacity in each biological experiment. **(J)** Paired comparison of mitochondrial oxidation capacity for each fuel between A375 and A375R cells. Lines connect three independent biological experiments; black bars indicate means. P values were obtained by paired two-sided t tests and adjusted across the three fuels using the Holm method.

## Supplementary Figure S3

**Fig. S3. Translational, 5′UTR, and proteomic analyses of the A375R response to CR-1-31-B. (A)** Representative normalized polysome profiles from one of two experimental batches, showing three of six biological replicates per condition after 1 h treatment of A375R cells with CR-1-31-B (10 nM) or DMSO. The 80S monosome peak and heavy-polysome region are indicated. **(B)** Polysome/80S ratios across all six biological replicates per condition. Symbols represent biological replicates and horizontal lines show means. The P value was obtained using an unpaired two-sided Welch t test on log2-transformed ratios. **(C,D)** Principal-component analysis (PCA) of batch-adjusted input RNA (C) and polysome-associated RNA (D) from the 1 h experiment. Colors denote treatment and symbols denote experimental batch. **(E)** Comparison of Anota2Seq APV effects with DESeq2 relative-TE interaction effects. Spearman ρ was 0.88 across all genes and 0.95 among Anota2Seq-selected genes; 2,214 of 2,678 Anota2Seq-selected genes (82.7%) also had a significant condition-by-fraction interaction in the DESeq2 sensitivity analysis (FDR < 0.05). **(F)** Hallmark and redundancy-reduced Reactome gene-set enrichment analysis (GSEA) of the 1 h A375R TE response to CR-1-31-B. **(G)** Covariate-adjusted associations between features of one representative, publicly annotated 5′UTR per gene and transcript classes with decreased TE, decreased polysome-associated RNA, or concordant decreases at both levels. Color denotes adjusted log-odds for membership in the corresponding CR-1-31-B-down class: orange indicates a positive association or enrichment and blue indicates a negative association or depletion. Point size denotes −log₁₀ FDR, capped at 10. The local-folding coefficient was oriented so that orange indicates stronger predicted stability. **(H)** Distributions of purine fraction, cap-proximal purine fraction, maximum G4Hunter score in 25-nt windows, and local folding stability (−minimum free energy in 100-nt windows) for unchanged (n = 9,579) and TE-down (n = 1,049) transcripts. The TE-down set is the analyzable subset of the 1,090 translation-mode-decreased transcripts. Violin plots show distributions, and internal boxplots show medians and interquartile ranges. Adjusted association FDR values are shown. For display, distributions were limited to the 1st–99th percentiles; all observations were retained in the analyses. **(I)** PCA of quantitative proteomes from A375 and A375R cells treated for 36 h with CR-1-31-B (10 nM) or DMSO (n = 4 per condition). **(J)** Hallmark and redundancy-reduced Reactome GSEA of the 36 h A375R proteomic response to CR-1-31-B. In F and J, positive NES indicates enrichment after CR-1-31-B and negative NES indicates depletion; dot size denotes −log₁₀ FDR. Only pathways with FDR < 0.05 are shown. Up to eight nonredundant representative pathways are displayed in each direction.

## Supplementary Figure S4

**Fig. S4. Resistance-associated proteomic programs opposed by CR-1-31-B and comparison with melanoma PDX treatment states. (A)** Numbers and fractions of resistance-up and resistance-down proteins, defined by FDR < 0.05 and log_2_ fold change ≥ 0.5 or ≤ −0.5, respectively, in A375R relative to A375, that changed in the opposing direction after CR-1-31-B. The stringent subsets additionally met the predefined CR-response threshold (FDR < 0.05 and |log_2_ fold change| ≥ 0.5). **(B)** Hallmark and Reactome over-representation analyses of the strict 65-protein subset that was increased in A375R relative to A375 and significantly decreased after CR-1-31-B. The analysis background comprised all proteins quantified in both proteomic comparisons. Only pathways with FDR < 0.05 are shown; dot position indicates fold enrichment, dot size indicates the number of overlapping proteins, and color indicates −log10 FDR. **(C)** Representative immunoblot of AXL in A375 and A375R cells treated for 24 h with 0, 10, or 100 nM CR-1-31-B. The approximately 120–140-kDa species corresponding to full-length, glycosylated AXL is shown; β-actin served as the loading control. The resistance-up/CR-1-31-B-down AXL pattern was observed in two independent experiments; the displayed concentration series is representative. The gap between the A375 and A375R samples corresponds to an unloaded lane on the same membrane; the image was not spliced. **(D)** Continuous treatment timelines and stage definitions for the WM4007 and WM4237 BRAF^V600E^ melanoma PDX trajectories in GSE245582. Treatment with the BRAF inhibitor dabrafenib plus the MEK inhibitor trametinib (D+T) began after the pretreatment time point. Each time point represents one tumor-bearing mouse. For each tumor, two nearby Visium sections from the same tissue block were averaged to generate one tumor-level estimate and were not treated as biological replicates. **(E)** Curated matrix of 16 representative genes induced by D+T in both PDX models (log2 effect ≥ 0.5 at one or more independently defined stages) whose corresponding proteins were significantly decreased by CR-1-31-B in A375R cells (log2 fold change ≤ −0.5; FDR < 0.05). PDX columns show stage-specific log2 effects in WM4007 and WM4237; the final column shows the A375R CR-1-31-B-versus-DMSO protein response at 36 h. Colors are capped at ±2, dark corner triangles identify values beyond the color limit, and gray slashes identify stage effects for which adjacent sections were not direction-consistent. Functional categories are descriptive, and the displayed genes constitute a non-exhaustive curated set rather than a statistical ranking.

## Supplementary Figure S5

**Fig. S5. Bioenergetic, proteomic and metabolomic responses to CR-1-31-B associated with glutamine–TCA metabolism. (A)** Extracellular acidification rate (ECAR) and oxygen-consumption rate (OCR) assay profiles in A375 and A375R cells following the indicated 24-h treatments. Lines show means and ribbons show SEM across three independent biological experiments; dashed lines indicate compound injections. **(B)** Derived glycolytic and respiratory parameters expressed as mean log2 fold change relative to the matched DMSO condition. Color is capped at ±3. **(C)** Preranked GSEA of oxidative-phosphorylation (OXPHOS) programs after CR-1-31-B treatment of A375R cells. Ranks show polysome-associated RNA at 1 h and total RNA at 24 h; the complex-I set was obtained from MitoCarta 3.0. NES and FDR are indicated. **(D)** Concordance of the CR-1-31-B-containing metabolic response between A375 (CR-1-31-B/DMSO) and A375R (PLX4032+CR-1-31-B/PLX4032) across GC-MS and LC-MS metabolites. Colors distinguish concordant decreases, concordant increases, and discordant responses; Spearman statistics and directional concordance are shown. **(E)** Fractional enrichment of glutamine m+5, glutamate m+5, α-ketoglutarate m+5, fumarate m+4, and malate m+4 over 0.5-6 h of [U-¹³C₅]glutamine tracing. Faint trajectories show two independent biological experiments and bold trajectories show means. **(F)** Oxidative citrate, isocitrate, and cis-aconitate m+4 labeling. **(G)** Reductive citrate-family m+5 labeling. **(H)** Glutamine-derived 2-hydroxyglutarate m+5 labeling. In F-H, faint trajectories show independent biological experiments and bold trajectories show means. **(I)** Relative metabolite-pool abundance and 1 h isotopologue composition in A375 cells, with A375R succinate shown for comparison. Bars and symbols are displayed as in Fig. 5F. **(J)** Cell-number-normalized glucose and glutamine consumption after 24 h treatment. A375 compares DMSO with CR-1-31-B, whereas A375R compares PLX4032 with PLX4032 plus CR-1-31-B. Lines connect six paired biological experiments; displayed P values are from two-sided paired *t* tests. **(K)** Representative immunoblots of GLS2 and ATP6V1F in A375 and A375R cells treated for 24 h with 0, 10 or 100 nM CR-1-31-B. β-actin served as a loading control. **(L)** Pathway-contextualized view of glutamine–TCA metabolite pools and protein abundance in A375R cells. Metabolite nodes show 24-h log₂ fold changes for PLX4032 plus CR-1-31-B versus PLX4032 (n = 3 independent biological experiments per condition), and the lower matrix shows 36-h protein-abundance log₂ fold changes for CR-1-31-B (10 nM) versus DMSO for 18 proteins selected for biochemical relevance irrespective of response direction or significance (n = 4 biological replicates per condition). Colors encode signed log₂ fold change, and open circles denote Benjamini–Hochberg FDR < 0.05. The crossed cell marks GLUD2 as not detected; dashed nodes mark succinyl-CoA and oxaloacetate as not quantified. Because the layers differ in treatment contrast and time point, the display is descriptive and does not establish cross-layer mediation; arrows show pathway topology, not measured flux. Full results are provided in Supplementary Table S5. **(M)** Total mRNA abundance of GLS1, NDUFA8, and ACADS after CR-1-31-B treatment (10 nM, 48 h), measured by RT-qPCR and expressed as log₂ fold change relative to matched DMSO controls. GLS1, n = 4; NDUFA8 and ACADS, n = 3 independent biological experiments. Symbols represent biological experiments; bars show mean ± SEM. **(N)** GDH1/2 mRNA abundance after CR-1-31-B treatment (7 nM, 72 h), measured by RT-qPCR and expressed as log₂ fold change relative to DMSO. The primer assay does not distinguish GLUD1 from GLUD2. Symbols represent three independent biological experiments; bars show mean ± SEM. **(O)** Selected glutamine-linked metabolites in A375 and A375R cells cultured for 24 h with 4, 0.25, or 0 mM glutamine. Mean log₂ fold changes relative to the matched 4 mM condition were calculated across three independent biological experiments and tested against zero using two-sided one-sample *t* tests, with Benjamini–Hochberg adjustment across measured metabolites within each cell-line/concentration comparison.

Color denotes mean log₂ fold change and is capped at ±4; exact values beyond the color limit are printed within the corresponding points. Dot size denotes −log₁₀ FDR and is capped at 6 for display; black outlines denote FDR < 0.05 and gray outlines denote FDR ≥ 0.05. **(P)** Relative viable-cell numbers in A375 and A375R cells cultured for 72 h at the indicated glutamine concentrations with vehicle or CR-1-31-B (3.5 nM). Values from three independent biological experiments were normalized to the matched vehicle-treated 4 mM glutamine condition. Small symbols represent biological experiments; large symbols and lines show means, and error bars show SD.

## Supplementary Figure S6

**Fig. S6. Body-weight changes and individual A375 xenograft trajectories during BRAF and eIF4A inhibition. (A)** Body-weight change relative to each mouse’s treatment-day-1 value. Small symbols show individual measurements and lines show group means; group-time points are displayed only while at least five mice contributed. Cohorts comprised vehicle n = 5, PLX4720 n = 11, CR-1-31-B n = 8, and PLX4720+CR-1-31-B n = 11. **(B)** Individual A375 xenograft-volume trajectories through treatment day 75, separated by treatment group. No values were imputed after a mouse ceased contributing. Cohorts comprised vehicle n = 5, PLX4720 n = 11, CR-1-31-B n = 8, and PLX4720+CR-1-31-B n = 11.

