## Supplementary Figures for "eIF4A inhibition disrupts resistance-associated translational and metabolic programs in BRAF-mutant melanoma"

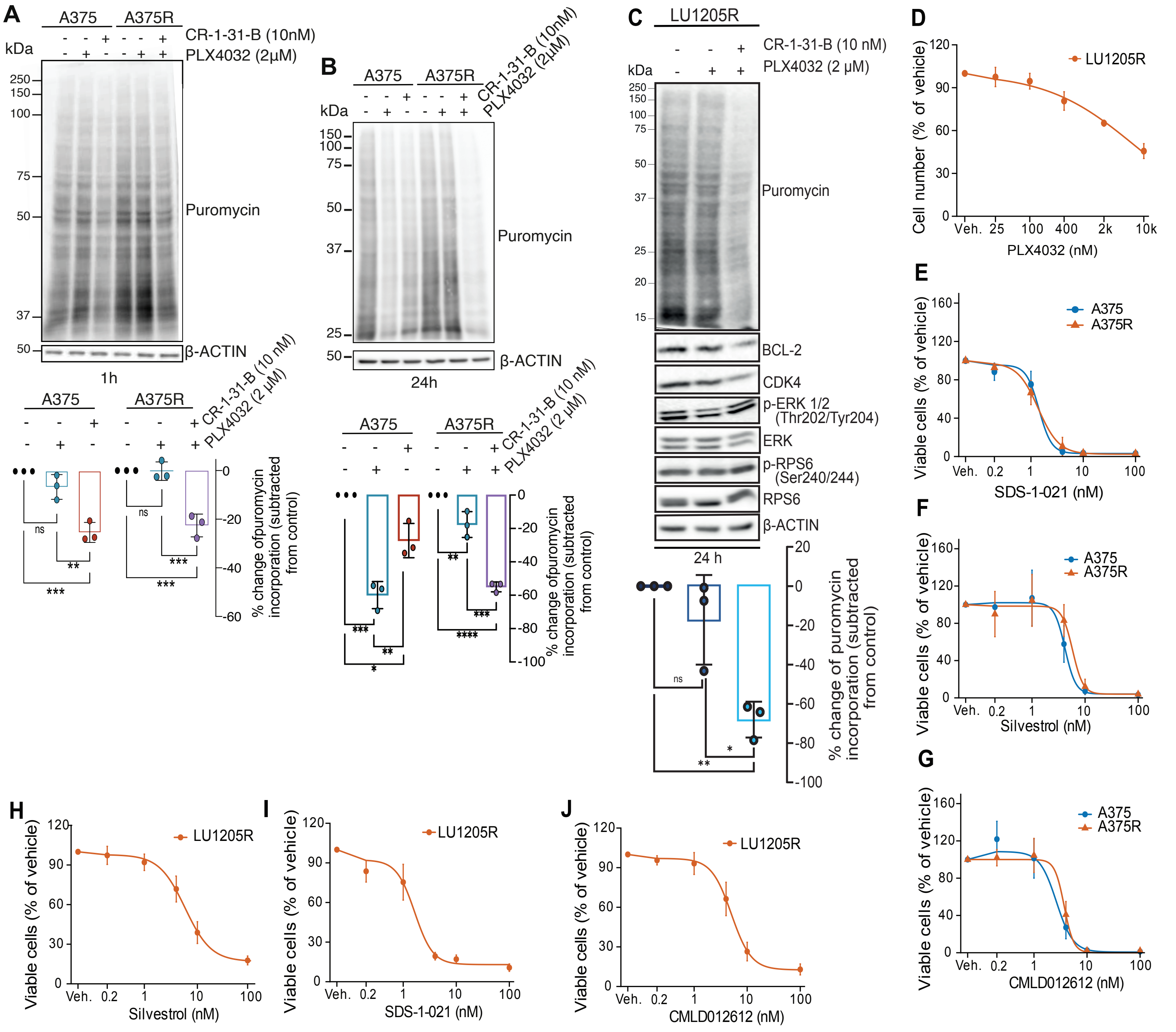


**K** **Summary of fitted $IC_{50}$ values (nM)**

**Melanoma models tested across multiple compounds**

| Compound | A375 | A375R | LU1205R |
|---|---|---|---|
| PLX4032 | 124 ± 27 | >10,000* | 6,711 |
| CR-1-31-B | 3.89 | 3.77 | 6.04 |
| SDS-1-021 | 1.43 | 1.38 | 1.65 |
| Silvestrol | 4.18 | 5.81 | 5.87 |
| CMLD012612 | 2.65 | 3.81 | 5.04 |

**Additional models**

| Compound | YUMMER1.7 | MDA-MB-231 | MDA-MB-231R |
|---|---|---|---|
| CR-1-31-B | 1.67 | 0.712 | 1.377 |

*IC50 was not reached within the tested range in the representative experiment.

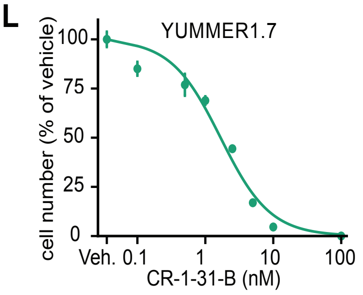


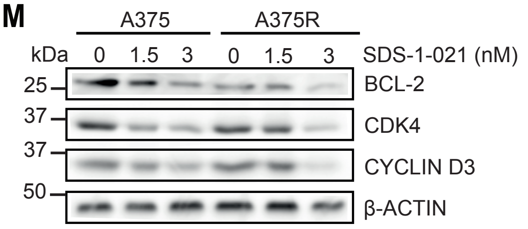


**Fig. S1.** Supporting analyses for Fig. 1. Page 1.

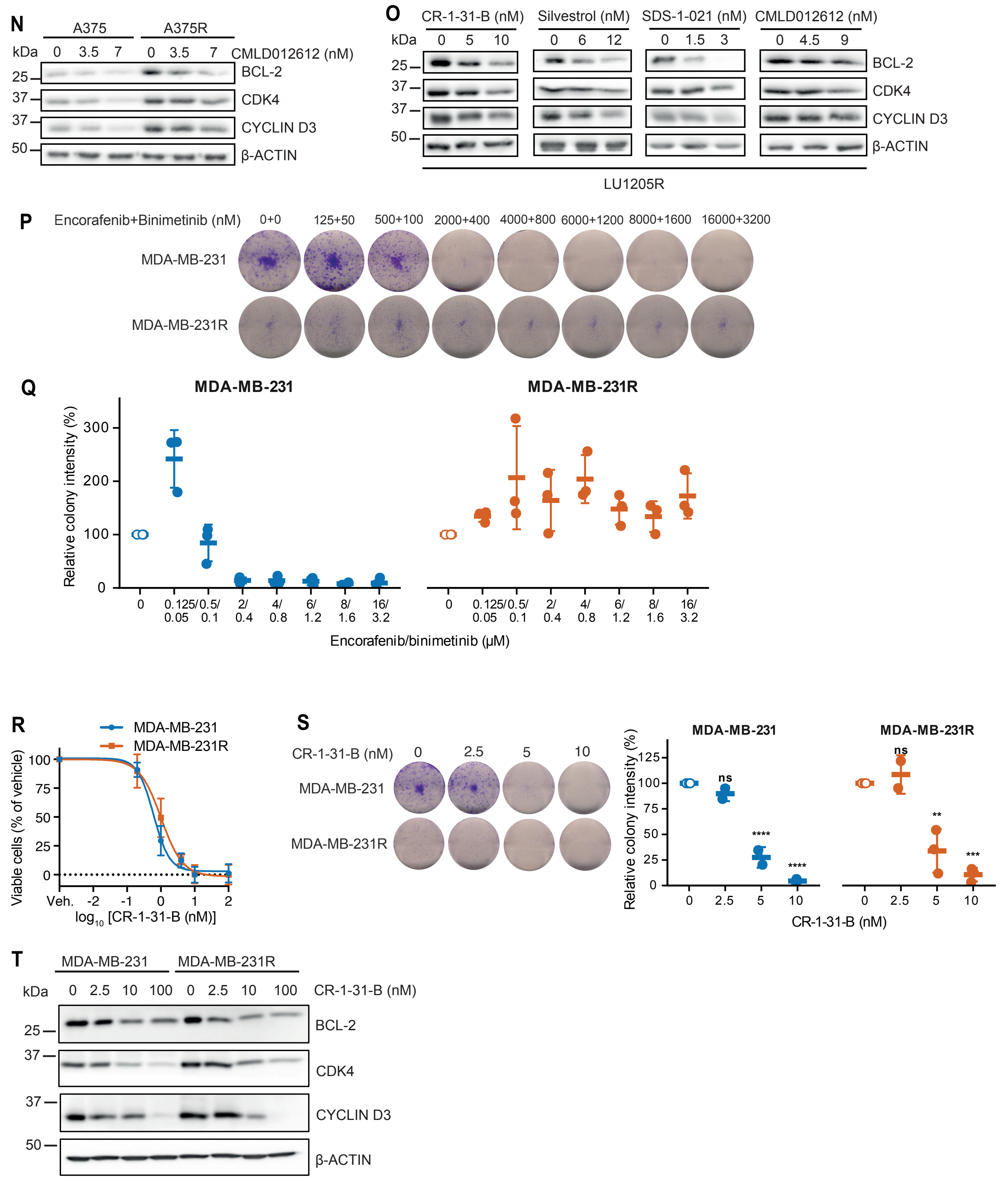


**Fig. S1.** Supporting analyses for Fig. 1. Page 2.

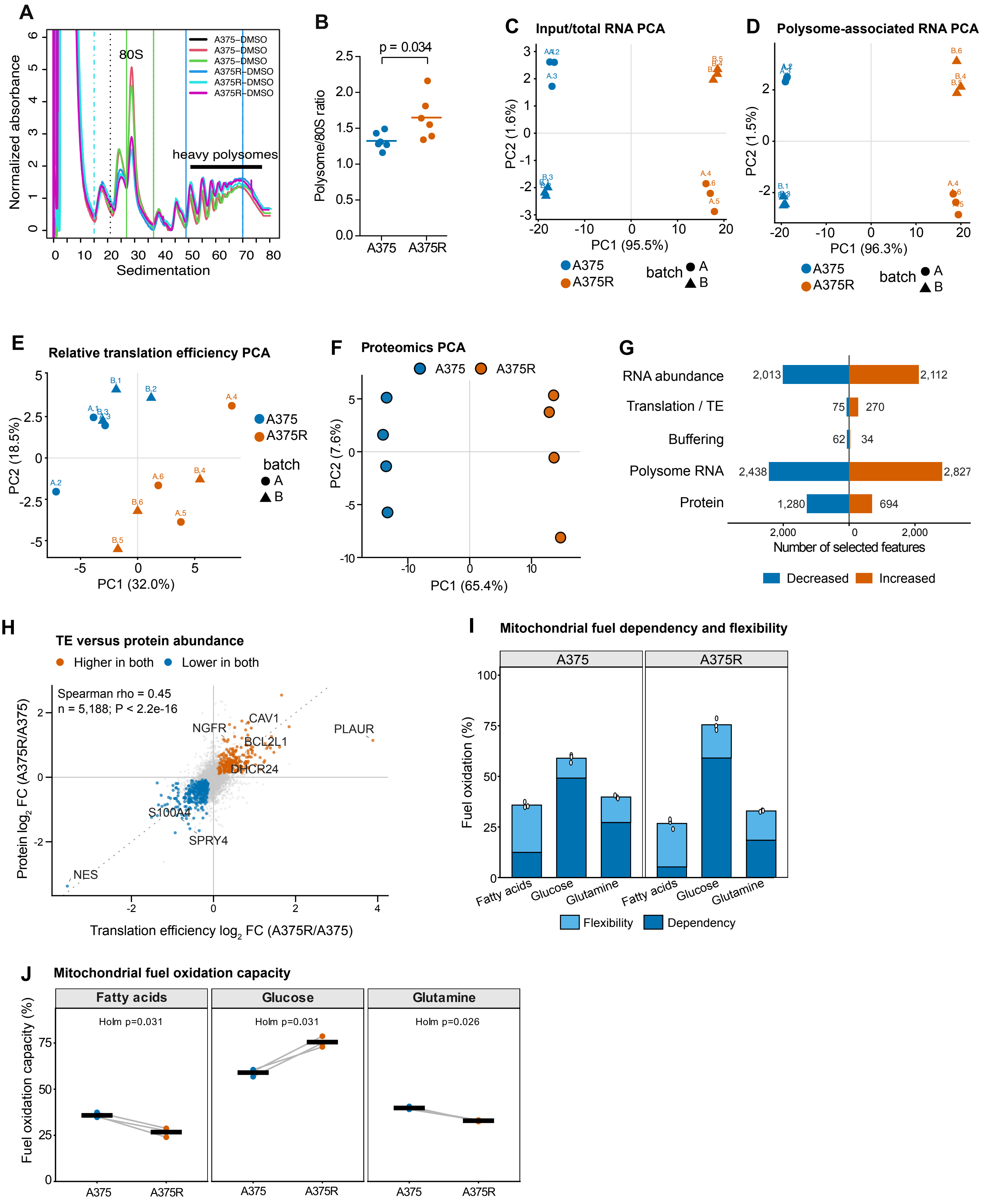


**Fig. S2.** Supporting analyses for Fig. 2.

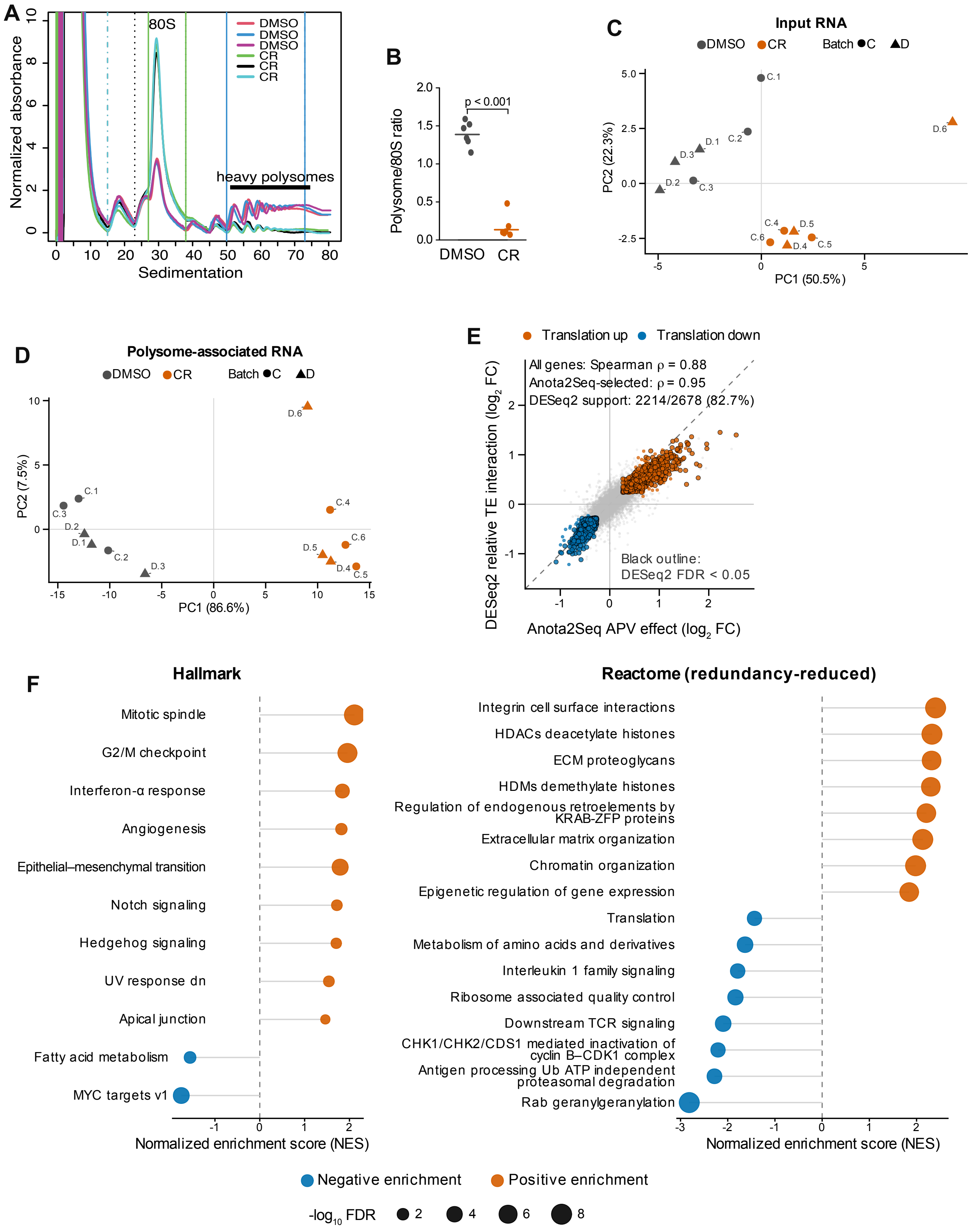


**Fig. S3.** Supporting analyses for Fig. 3. Page 1.

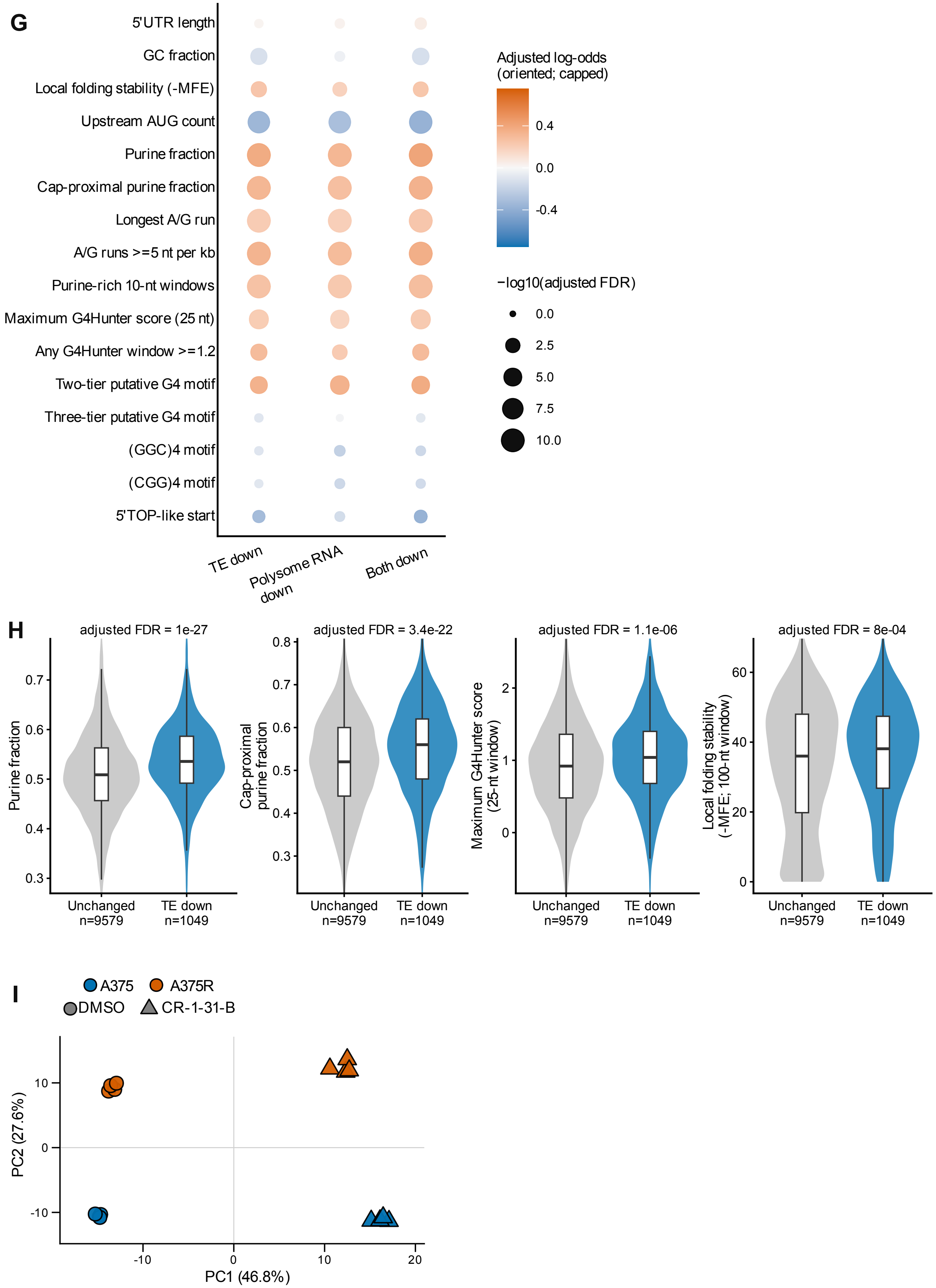


**Fig. S3.** Supporting analyses for Fig. 3. Page 2.

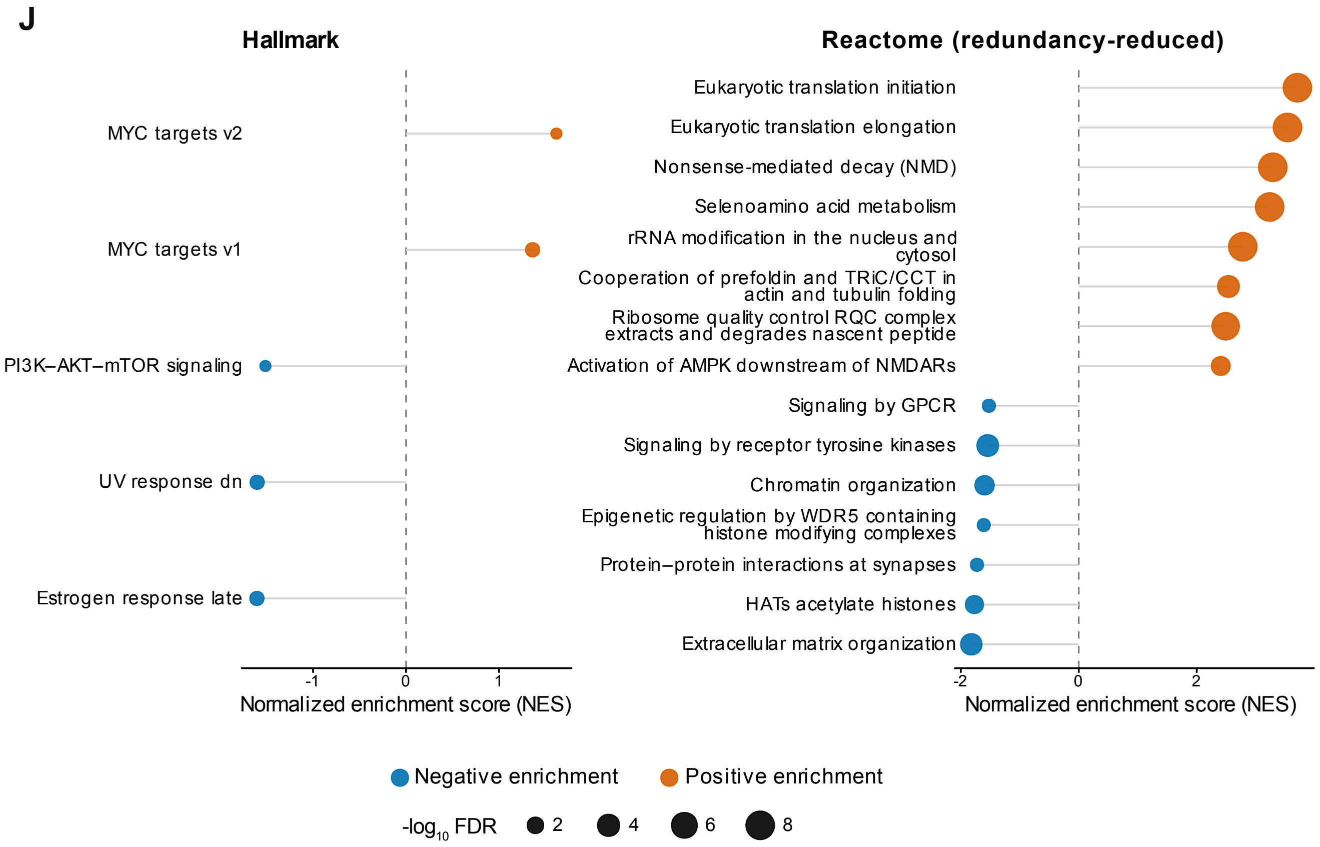


**Fig. S3.** Supporting analyses for Fig. 3. Page 3.

**A375/A375R resistance-associated proteome and AXL validation**
Contrasts: A375R vs A375; CR-1-31-B vs DMSO in A375R cells (36 h)

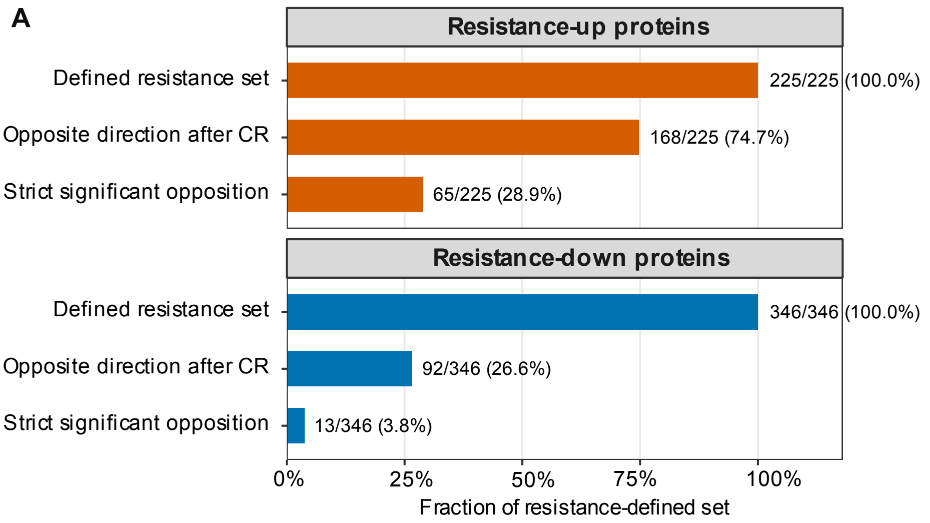


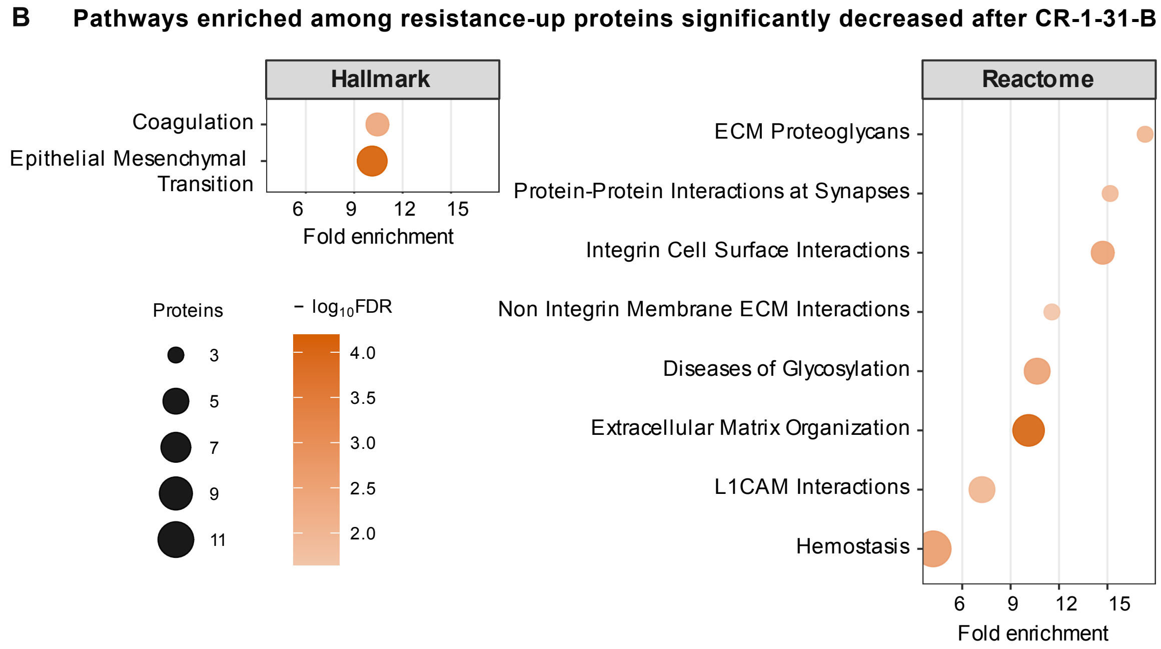


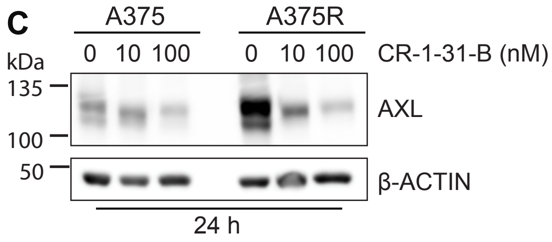


**Fig. S4.** Supporting analyses for Fig. 4. Page 1.

**D** **GSE245582 PDX treatment time course and stage definitions**

BRAF V600E melanoma PDXs; continuous dabrafenib + trametinib (D+T) initiated after T0

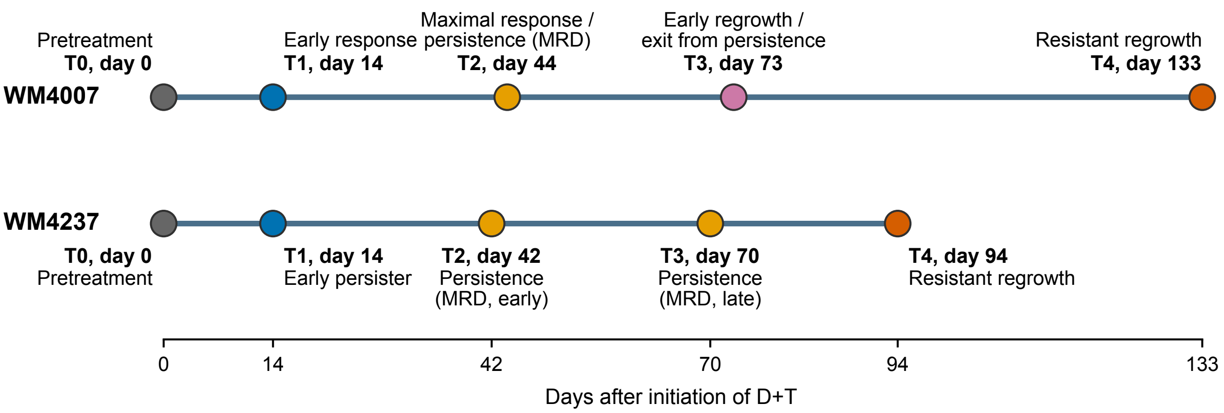


Each time point represents a different tumor-bearing mouse. Two nearby Visium sections from the same tumor block were averaged to generate one tumor-level estimate and were not treated as biological replicates.

**E** **PDX treatment-stage effects and the independent A375R protein response to CR-1-31-B**

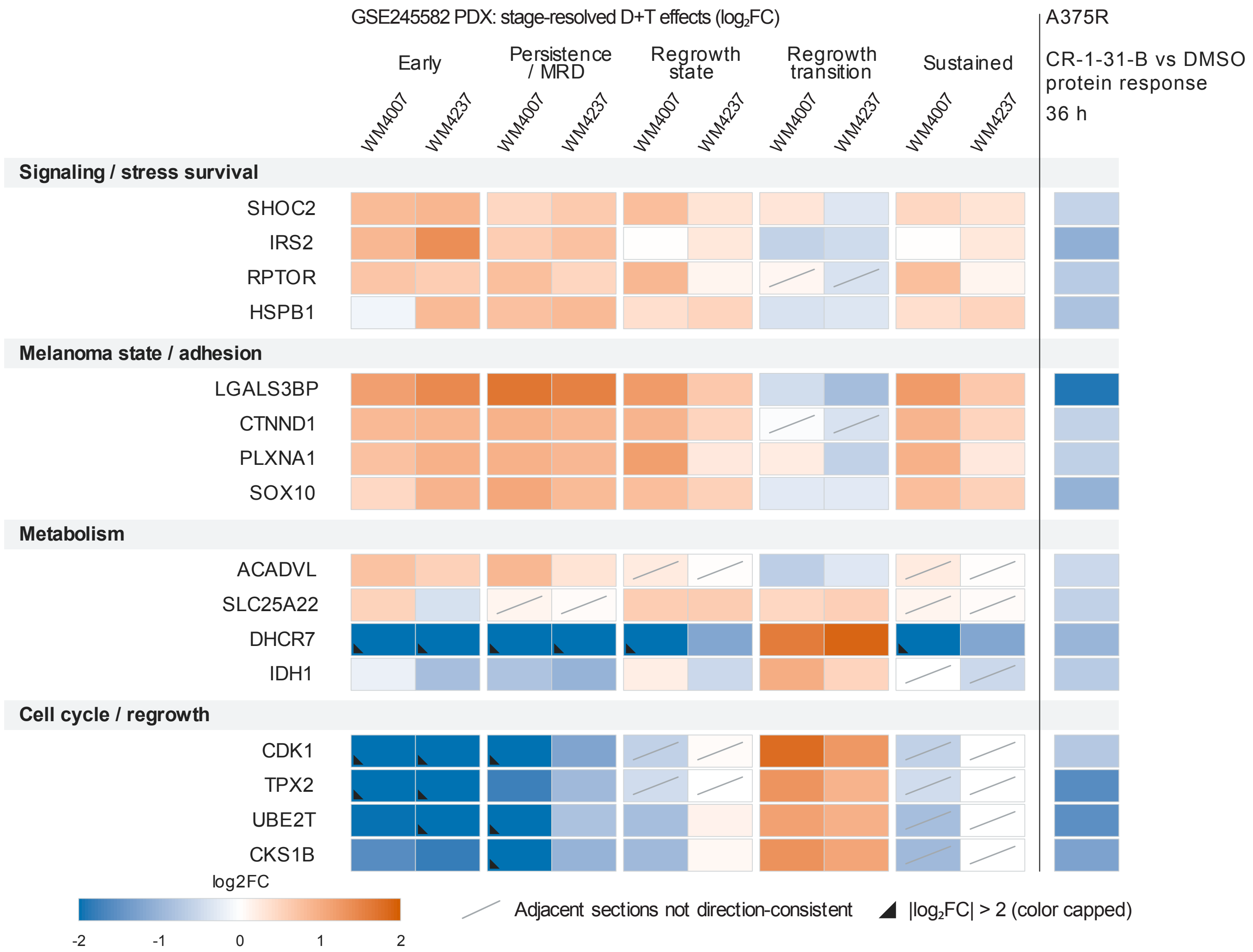


**Fig. S4.** Supporting analyses for Fig. 4. Page 2.

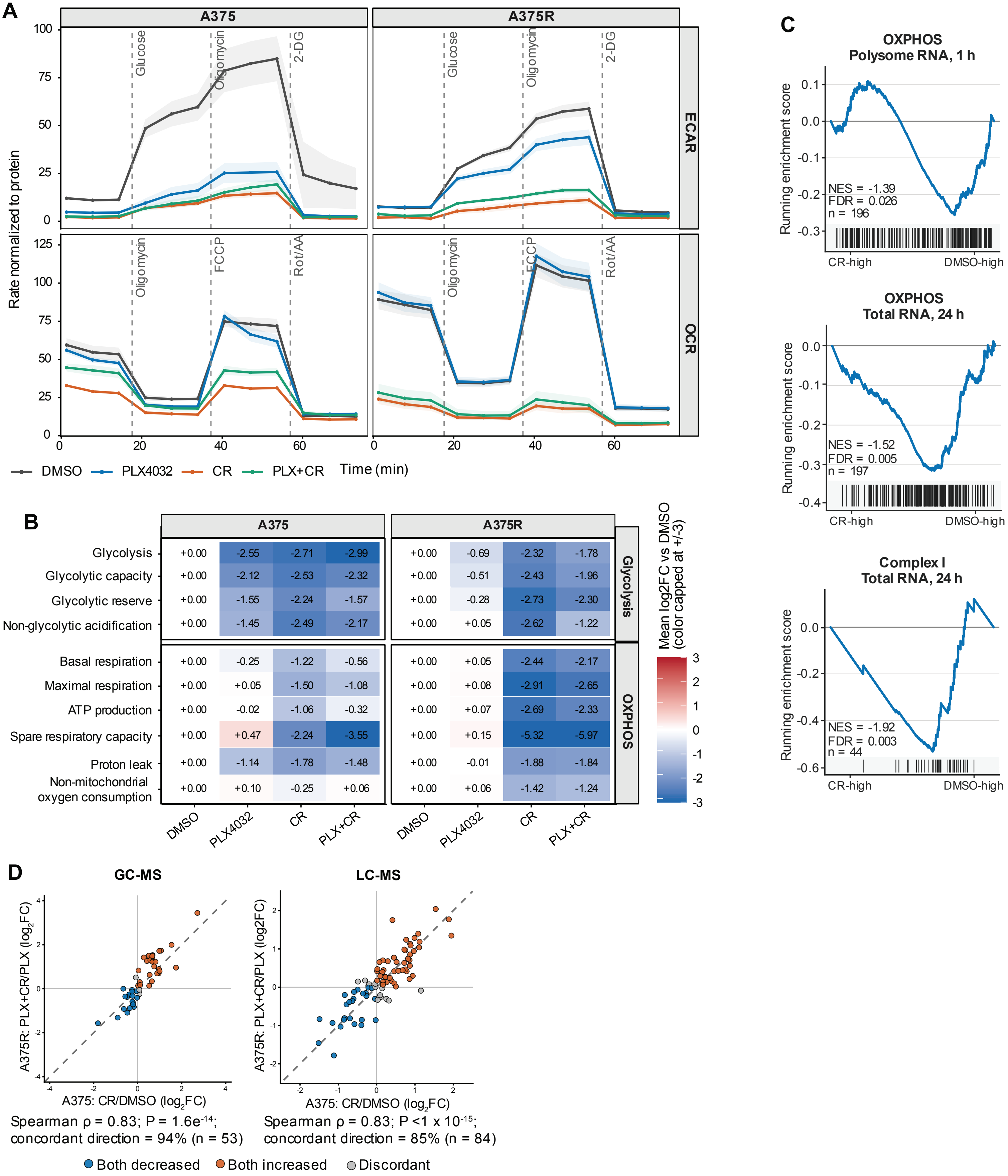


**Fig. S5.** Supporting analyses for Fig. 5. Page 1.

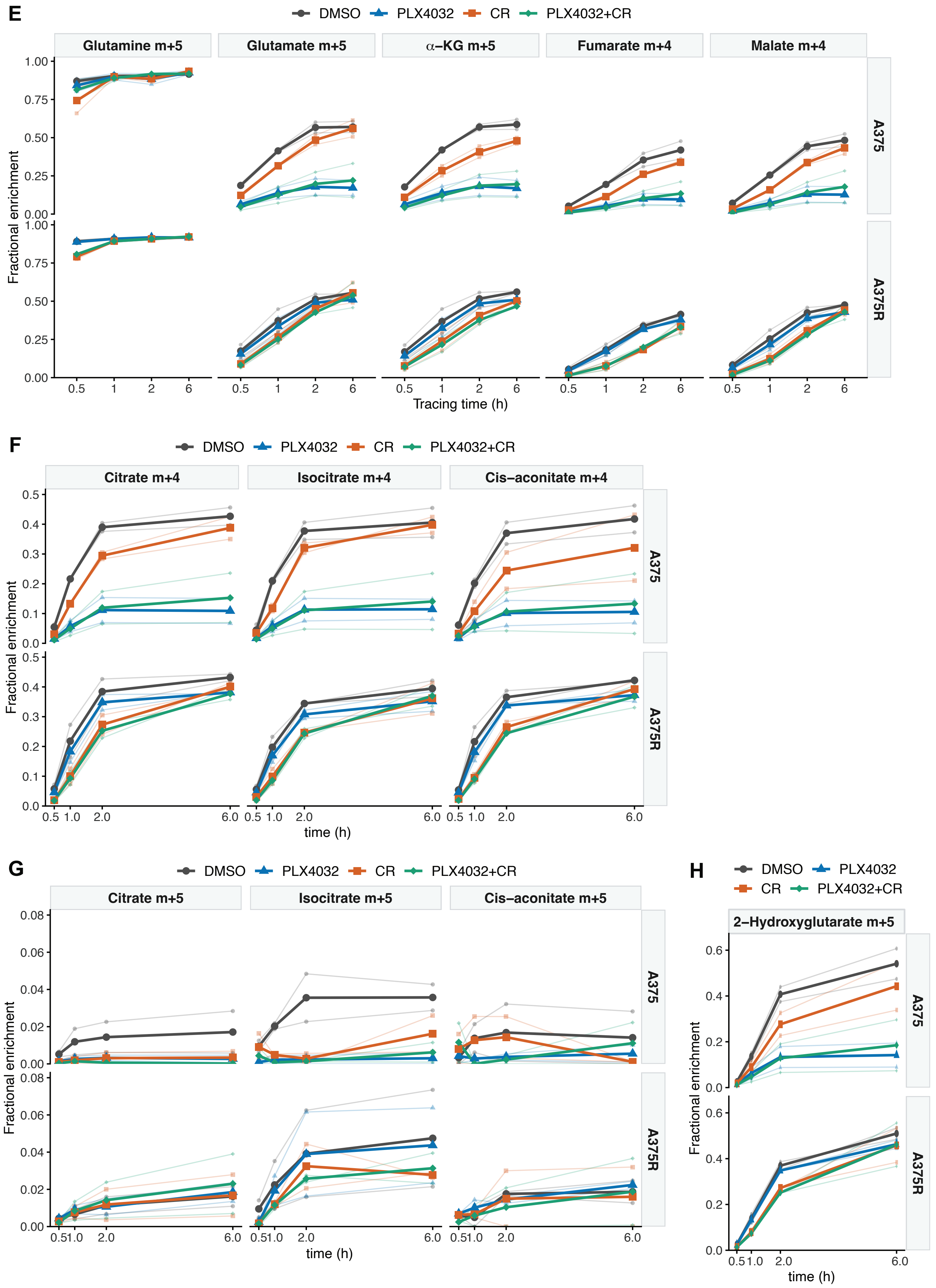


**Fig. S5.** Supporting analyses for Fig. 5. Page 2.

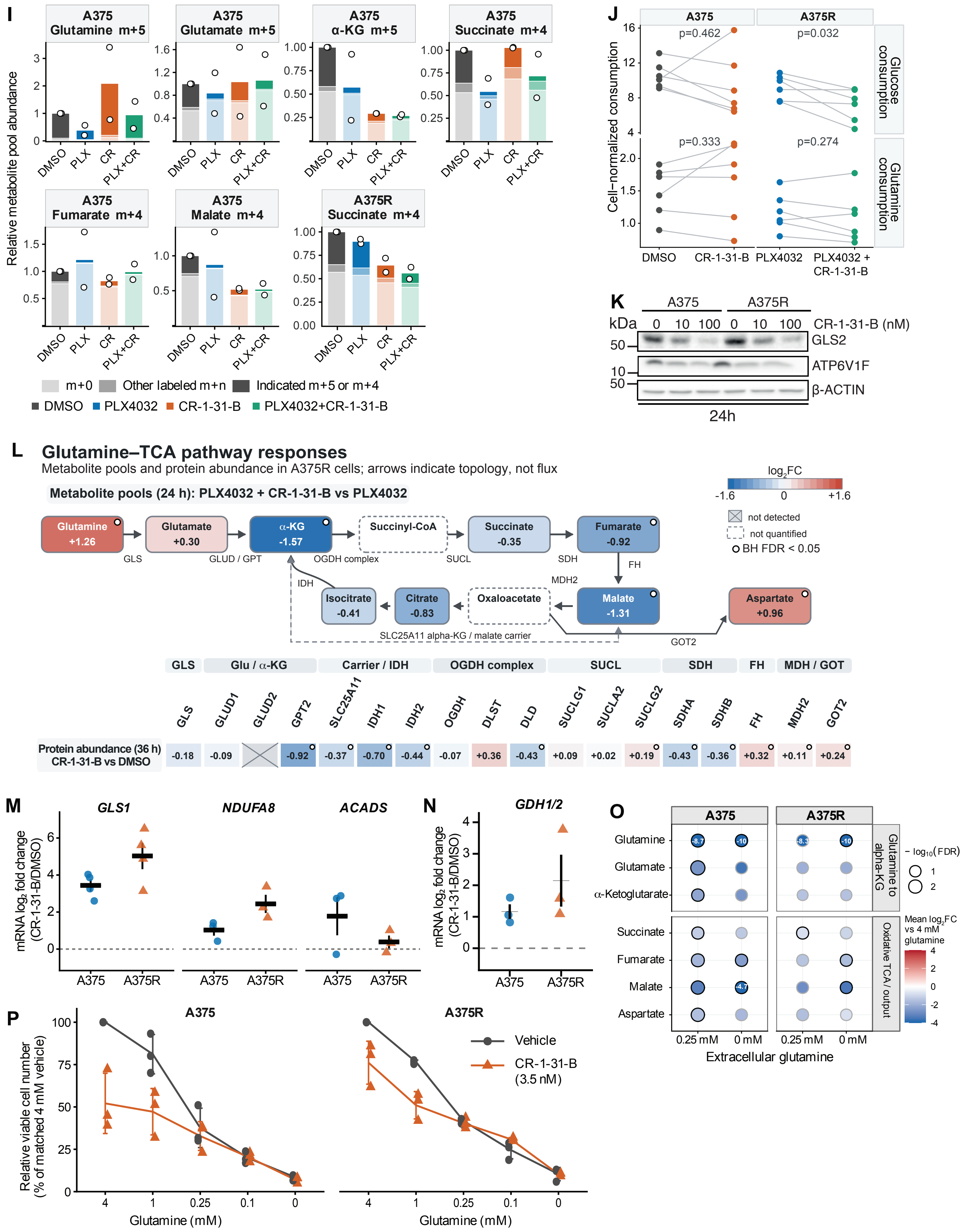


**Fig. S5.** Supporting analyses for Fig. 5. Page 3.

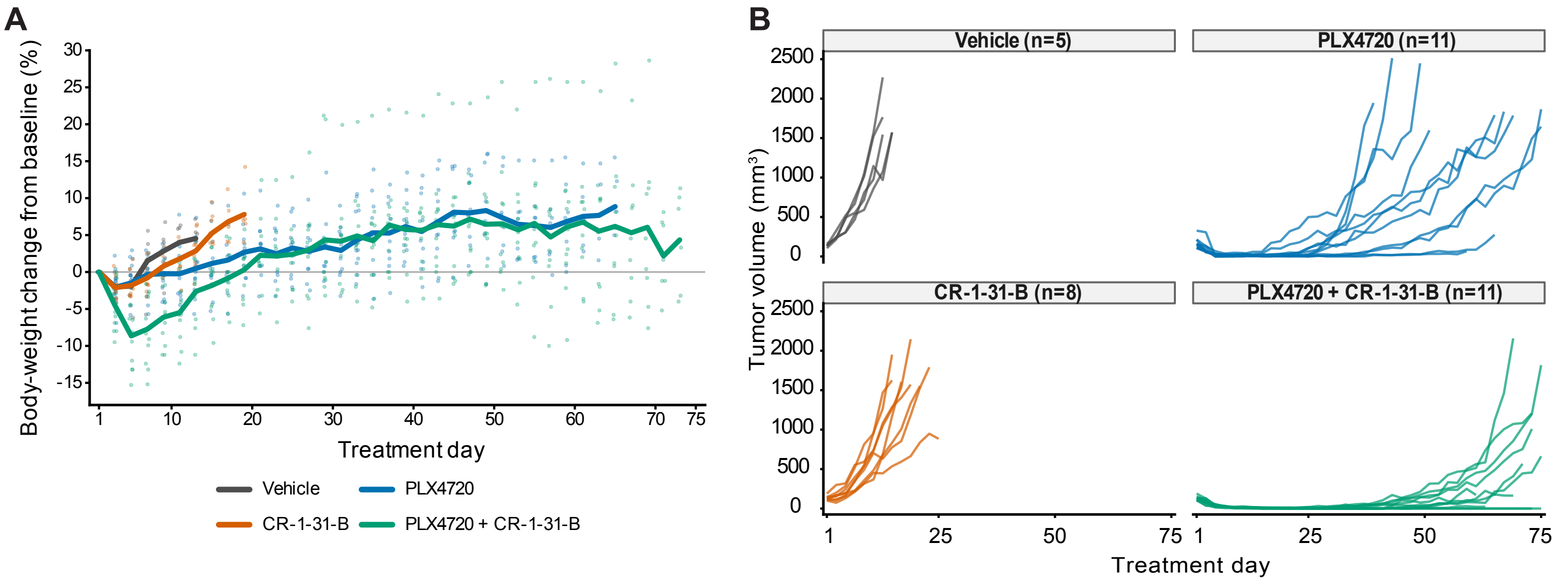


**Fig. S6.** Supporting analyses for Fig. 6.
